# A Single-Cell and Spatial Atlas of MVP-PAN Evolution Reveals a Desmoplastic Dependency to Overcome Glioblastoma Resistance

**DOI:** 10.64898/2025.12.25.696471

**Authors:** Fei Wang, Xin Wu, Guozheng Zhao, Run Huang, Chen Yang, Yuhan Bai, Wenqian Cao, Yue Lu, Guangling Xu, Haohao Qiu, Hongyi Ling, Dengfeng Lu, Youjia Qiu, Juyi Zhang, Bixi Gao, Yanbo Yang, Ting Sun, Zhouqing Chen, Zhong Wang

## Abstract

Therapeutic resistance in IDH-wildtype glioblastoma (GBM) is driven by an intricate interplay between cellular plasticity and protective microenvironmental niches. By constructing the GRIT-Atlas—a massive transcriptomic compendium encompassing nearly one million high-quality single cells across 17 cohorts—and cross-analyzing it with 48 independent Visium spatial transcriptomics sections, we discover a recurrent “Spatial Resistance Triad” comprising mesenchymal-like (MES-like) malignant cells, myeloid-derived suppressor cells (MDSCs), and collagen-secreting cancer-associated fibroblasts (CAFs). This large-scale spatial integration reveals that the triad physically fortifies microvascular proliferation (MVP) and pseudopalisading necrosis (PAN) niches. To substantiate these findings at true single-cell resolution, we deploy high-plex spatial molecular imaging (CosMx SMI) to map over 400,000 single cells across a clinical validation cohort. This high-resolution architecture firmly validates the triad’s geography and, critically, provides structural gradient evidence unmasking a spatiotemporal continuum wherein PAN emerges as a direct functional consequence of MVP advancement and subsequent vascular collapse. Mechanistically, stromal CAF-derived COL6A1 engages CD44 receptors on MES-like cells to accelerate a potent neural stemness cascade within these protective domains. Utilizing a blood-brain barrier (BBB)-penetrant multi-library compound screen, we identify Lacidipine as a multi-modal stromal disruptor that successfully silences CAF activation and halts matrix secretion. Incorporating Lacidipine into the standard chemo-immunotherapy backbone (TMZ + anti-CSF1R) completely dismantles the protective desmoplastic matrix niche, forcing a profound collapse of intracranial tumor burden and dramatically extending overall survival in orthotopic models. Collectively, our study leverages unprecedented single-cell and spatial scale to provide a definitive blueprint of GBM therapeutic evasion, establishing matrix-targeted intervention as a mandatory prerequisite for successful glioblastoma eradication.

## Introduction

IDH-wildtype glioblastoma (GBM) represents the apex of therapeutic resistance among primary brain malignancies^1^. Despite an aggressive standard-of-care (SOC) regimen involving maximal safe resection followed by radiotherapy and temozolomide, recurrence is inevitable, resulting in a dismal median survival of approximately 15 months^2,3^. This intractability stems not merely from a static immunosuppressive tumor microenvironment (TME), but from the profound cellular plasticity and heterogeneity inherent to GBM^4,5^. Malignant cells exist as a dynamic ecosystem, capable of transitioning between functional states—ranging from neurodevelopmental to injury-response phenotypes—to evade therapeutic pressure^6,7^. However, the specific evolutionary trajectories driven by distinct therapeutic modalities, particularly how treatment actively reshapes the cellular hierarchy to favor resistant phenotypes, remain poorly defined.

The promise of immune checkpoint blockade (ICB) in GBM has been tempered by phase III clinical failures, necessitating a deeper interrogation of resistance mechanisms^8,9^. While the genomic landscape of primary versus generic recurrent tumors is well-charted^10,11^, and inquiries into ICB responses are accumulating^12–14^, there remains a paucity of systematic studies that definitively delineate the perturbation landscapes specific to standard of care, ICB, and combinatorial targeted therapies. Crucially, these cellular perturbations do not occur in isolation but within the highly organized spatial architecture of the TME^15–17^. As the TME functions as a structured ecosystem rather than a disordered collection of cells, understanding how diverse therapeutic stresses specifically reconfigure spatial neighborhoods to orchestrate immune exclusion represents a critical knowledge gap.

We hypothesized that the resistance to immunotherapy is physically anchored within specific histological niches. GBM is pathologically defined by two hallmark features: microvascular proliferation (MVP) and pseudopalisading necrosis (PAN).^18,19^ Rather than mere morphological artifacts of rapid growth, we posit that these hypoxic, structurally distinct regions function as ‘immunosuppressive sanctuaries.’ Within these spatial confines, instead of mere immune evasion, specialized malignant subpopulations actively recruit and reprogram stromal and myeloid partners to erect a physical barrier that selectively excludes cytotoxic effectors while nurturing a pro-tumorigenic microenvironment. Unraveling this spatial “code” requires integrating high-resolution single-cell transcriptomics with spatially resolved multi-omics to map the functional geography of resistance.

To address these challenges, we constructed the Glioblastoma Resistance Insights from Treatment Atlas (GRIT-Atlas), the largest and most comprehensive integrated single-cell transcriptomic resource for IDH-wildtype GBM to date, comprising nearly one million high-quality cells from 296 clinical samples. Uniquely, the GRIT-Atlas captures the full spectrum of disease evolution, including primary tumors, recurrences post-standard chemoradiotherapy, and rare cohorts of recurrences following ICB monotherapy and ICB plus anti-angiogenic target combination therapy, stratified by clinical response. By synergizing deep single-cell phenotyping with large-scale spatial transcriptomics across 48 independent Visium sections, we aimed to deconstruct the cellular and spatial logic of therapy-induced evolution.

Here, we report that therapeutic pressure drives a convergent evolution of malignant cells toward a specific, highly aggressive mesenchymal-like (MES-like) state defined by a potent synergy of hypoxia, inflammation, and stemness. By integrating our single-cell atlas with macro-scale spatial transcriptomics, we uncover the physical reservoir of these cells, defining a “Spatial Resistance Triad” composed of MES-like malignant cells, myeloid-derived suppressor cells (MDSCs), and collagen-secreting cancer-associated fibroblasts (CAFs). We demonstrate that this triad specifically colonizes the hypoxic MVP and PAN niches. To substantiate these topological findings at true single-cell resolution, we deployed high-plex spatial molecular imaging (CosMx SMI), mapping over 400,000 single cells across a validation cohort. This high-resolution geography not only firmly solidifies the triad’s microenvironmental alignment but, crucially, provides structural gradient evidence unmasking a spatiotemporal continuum wherein PAN emerges as the direct functional consequence of MVP advancement and subsequent vascular collapse. Mechanistically, stromal CAF-derived COL6A1 acts as the primary architectural driver, constructing a fibrotic scaffold via the COL6A1-CD44 axis that engages MES-like cells to accelerate a neural stemness cascade. Finally, through a blood-brain barrier (BBB)-penetrant compound screen, we identify Lacidipine as a multi-modal stromal disruptor. Combining Lacidipine with standard chemo-immunotherapy (TMZ + anti-CSF1R) completely dismantles this protective desmoplastic fortress, forcing a profound collapse of tumor burden and dramatically extending survival in orthotopic models. Collectively, our findings expose the spatial infrastructure of resistance in GBM and establish matrix-targeted intervention as a mandatory prerequisite for successful glioblastoma eradication.

## Results

### 1.1 Constructing the GRIT-Atlas: Delineating the TME Landscape Across Therapeutic Modalities

To systematically uncover the cellular dynamics driving therapeutic resistance in GBM, we compiled the GRIT-Atlas, an integrated transcriptomic resource spanning 978,065 high-quality cells across 296 clinical specimens (**Figure 1A, Table S1**). The finalized atlas encompasses distinct, treatment-demarcated cohorts: primary GBM (pGBM), post-Stupp protocol recurrences (rGBM-R+C), and two immunotherapy-exposed arms—PD-1 blockade monotherapy (rGBM-ICB) and PD-1 inhibition paired with anti-angiogenic targeted agents (rGBM-ICB+T) (**Figure 1B**). Following batch regularization and unsupervised clustering, we resolved seven fundamental cell lineages based on canonical marker expression (**Figure 1C, Figure S1D**): Myeloid (*CD14/CD68*), Lymphoid (*CD3D/CD8A*), Endothelial (*PECAM1/VWF*), Stromal (*COL1A1/FAP*), Oligodendrocytes (*MBP*), Astrocyte/Neurons (*GFAP/TUBB2B*), and Malignant tumor cells (*EGFR/PTPRZ1*). Due to the transcriptional overlap between neoplastic and normal neural cells, large-scale chromosomal alterations (CNVs) profiling was deployed to definitively segregate malignant cells—characterized by hallmark chromosome 7 amplification and chromosome 10 deletion^20^—from diploid neural populations (**Figure S1E, S1F**).

**Figure 1.**
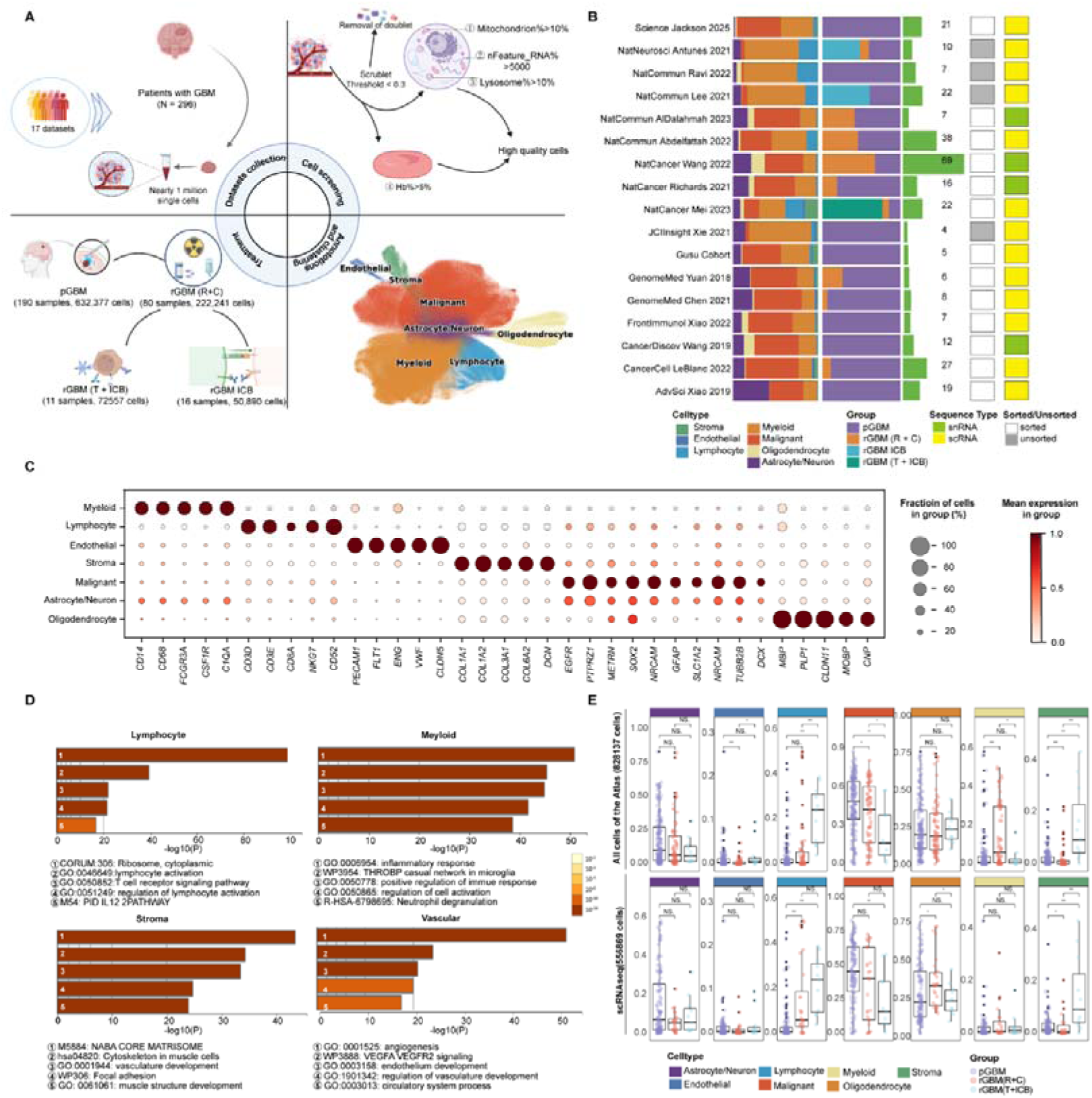
Construction and Single-Cell Landscape of the GRIT-Atlas in IDH-Wildtype Glioblastoma. **(A)** Schematic workflow of the study design, including multi-cohort data collection (17 cohorts), quality control, treatment stratification, and global integration. **(B)** Dataset metadata displaying cell-type composition, clinical treatment groups, cell/sample counts, FACS sorting status, and sequencing modalities across cohorts. **(C)** Dot plot of canonical marker gene expression across seven major cell lineages; dot size indicates the percentage of expressing cells, and color intensity reflects average expression. **(D)** Bar plots of the top 5 enriched biological pathways (Metascape) for the top 100 signature genes in lymphoid, myeloid, stromal, and endothelial populations. **(E)** Box plots comparing cell-type proportions across primary (pGBM), standard recurrent (rGBM-R+C), and combinatorial immunotherapy/targeted recurrent (rGBM-ICB+Target) tumors, presented for the full dataset (upper) and scRNA-seq data only (lower). Statistical significance was determined by the Wilcoxon rank-sum test. See also Figure S1.

Macro-environmental compositional analysis across the treatment continuum unmasked critical microenvironmental remodeling (**Figure 1E**). First, the relative proportion of malignant cells significantly contracted in recurrent tumors, reflecting the decreased tumor purity commonly observed in surgical recurrences^6^. Second, within the immune compartment, rGBM-R+C induced a parallel expansion of both myeloid and lymphoid infiltrates. Strikingly, the addition of combined ICB+Target therapy further amplified the lymphoid compartment while concurrently suppressing the principal immunosuppressive myeloid cell fraction. Finally, we observed a unique divergence within the non-immune compartment. While the endothelial cell fraction exhibited a minor, non-significant stabilization across cohorts after anti-angiogenic therapy, the stromal cell compartment demonstrated a highly robust, non-artifactual hyper-accumulation that persisted significantly in treatment Non-responders (**Figure S1G**). Collectively, these findings indicate that the stromal matrix component, rather than the endothelial vasculature, represents the core microenvironmental shift orchestrating therapy resistance.

### 1.2 High-Resolution Dissection of Malignant Cell Subpopulations Identifies Resistance-Associated States

To resolve the malignant transcriptional architecture driving resistance, consensus Non-negative Matrix Factorization (cNMF) was applied to the neoplastic compartment, partitioning malignant cells into 11 distinct functional states after filtering non-malignant lineage contamination (**Figure 2A, B**). Benchmarking against established classification frameworks categorized these programs into canonical states, including proliferating (cNMF8; driven by *MKI67/TOP2A*), OPC-like (cNMF1), AC-like (cNMF5), and NPC-like (cNMF4/11) states (**Figure S2F, G**). Intriguingly, we identified a malignant Neuron-like (NEU-like) state (cNMF6/10/12) marked by mature synaptic transcripts (*SYT1/GRIA4*). Despite their differentiated neural phenotype, these cells retained patient-specific CNVs and exhibited hyperactivated p53 and PI3K signaling (**Figure 2C, D**), confirming their neoplastic origin and propensity to mediate tumor-neuron synaptic integration within the infiltrating brain parenchyma during recurrence^21–23^.

**Figure 2.**
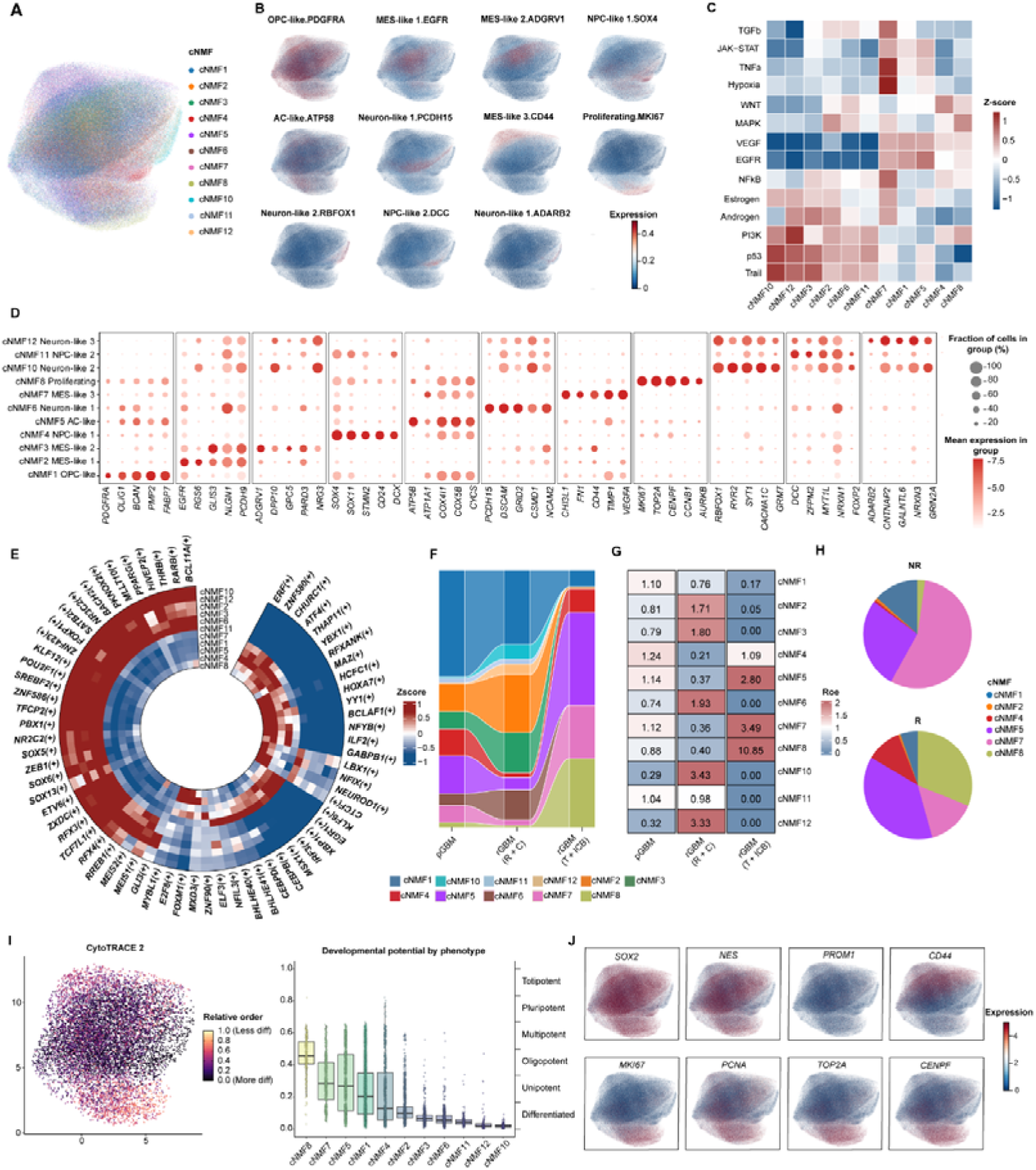
Identification of Therapy-Driven Malignant States and Evolutionary Trajectories in GBM. **(A)** UMAP visualization of malignant cells across all cohorts, colored by 11 consensus Non-negative Matrix Factorization (cNMF) meta-programs. **(B)** Feature plots illustrating the spatial distribution and usage intensity of individual cNMF programs. **(C)** Heatmap of 14 signaling pathway activities quantified by DecoupleR across cNMF programs. **(D)** Dot plot displaying characteristic marker genes defining each cNMF malignant state. **(E)** Circular heatmap of transcription factor regulon activity (pySCENIC) across the 11 cNMF programs (top 10 regulons per program). **(F, G)** Clonal evolution mapping across therapeutic modalities, shown via an alluvial plot depicting proportional shifts of cNMF states **(F)** and a Ratio of Observed to Expected (Ro/e) heatmap quantifying state enrichment across treatment groups **(G)**. **(H)** Pie charts comparing cNMF state composition between non-responders and responders within the rGBM-ICB+Target cohort. **(I)** Differentiation potential analysis using CytoTRACE2, visualized by global UMAP embedding (left) and box plots quantifying stemness scores across malignant clusters (right). **(J)** Feature plots contrasting canonical stemness markers (*SOX2, NES, PROM1, CD44*) with proliferation markers (*MKI67, PCNA, TOP2A, CENPF*). See also Figure S2.

Program cNMF7 was identified as the canonical MES-like state, characterized by profound tissue hypoxia signatures and hyperactivated TNFA, TGFB, and NFKB signaling pathways (**Figure 2D**). This state was molecularly defined by the coordinated upregulation of inflammatory (*CHI3L1/MIF*), invasive (*SPP1/VIM/FN1*), and angiogenic (*VEGFA*) machinery (**Figure S2H**). Gene Regulatory Network (GRN) inference revealed that while cNMF7 shared a basic transcriptional core with developmental states, its unique pathological identity was driven by a distinct master transcription factor triad: **ELF3**, **NFIL3**, and **BHLHE40** (**Figure 2E**).

Quantification of these malignant states across the treatment continuum revealed a severe evolutionary selection landscape (**Figure 2F-H**). Standard chemoradiotherapy preferentially enriched for the NEU-like states, suggesting a genotoxic stress-induced hijacking of neural circuitry to survive clearance. Strikingly, combined ICB and anti-angiogenic targeted therapy forced a massive Darwinian bottleneck, where the highly metabolic AC-like, proliferating, and classic MES-like (cNMF7) states expanded to constitute over 80% of the entire malignant compartment. Crucially, cNMF7 was almost exclusively restricted to treatment Non-responders. Trajectory mapping via CytoTRACE 2 positioned cNMF7 as a highly stem-like, plastic state specifically marked by CD44 expression, whereas the conventional stemness marker *PROM1* (CD133) was largely absent (**Figure 2I, J**). Although cNMF7 exhibited low baseline proliferation, its transcriptional signature robustly predicted dismal clinical survival across seven independent validation cohorts (**Figure S2I**). These data identify this stress-adaptive, quiescent cNMF7 population as the principal malignant engine of clinical non-response and immune evasion.

### 1.3 Treatment Pressure Selects for Immunosuppressive Myeloid Subpopulations Associated with ICB Resistance

Coarse-grained annotation partitioned the myeloid compartment into six major lineages, dominated by microglias (MGs; 54.4%) and bone marrow-derived macrophages (BMDMs; 37.1%) (**Figure 3A, B**). Under ICB or combination ICB+Target therapy, we observed a reciprocal shift characterized by contracted MG fractions and expanded BMDMs (**Figure 3C**), reflecting a mesenchymal-like microenvironmental transformation^6,14^. In contrast, dendritic cells (DCs) expanded across both immunotherapy arms, while neutrophils uniquely surged under combined ICB+Target pressure (**Figure 3C**).

**Figure 3.**
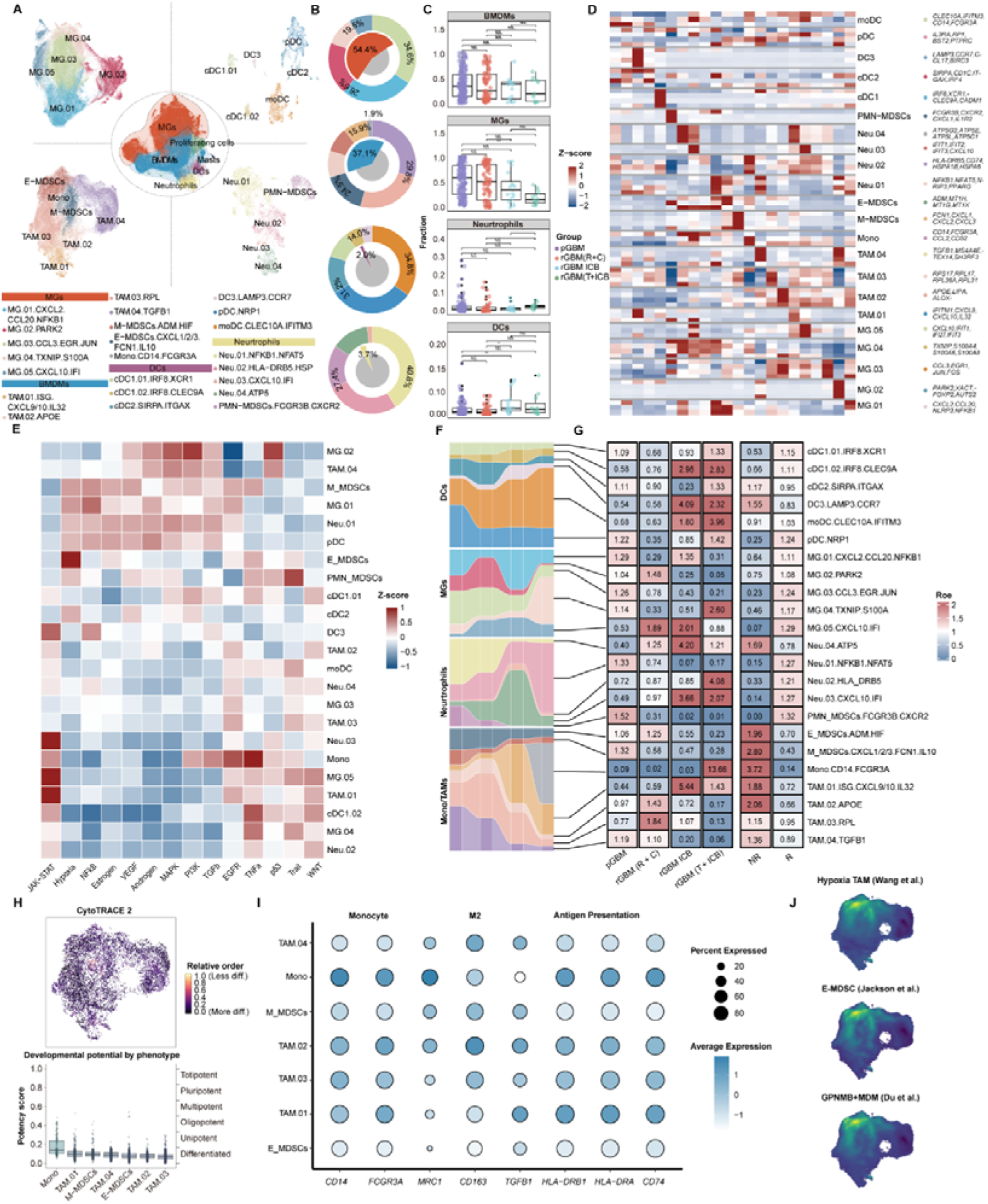
Hierarchy, Functional Heterogeneity, and Therapy-Induced Remodeling of the Myeloid Compartment. **(A)** High-resolution UMAP embedding of the myeloid landscape (center) with detailed sub-clustering of microglia (top-left), dendritic cells (top-right), BMDMs (bottom-left), and neutrophils (bottom-right). **(B)** Nested donut charts illustrating major lineage proportions (inner ring) and specific sub-cluster compositions (outer ring) within the myeloid compartment. **(C)** Box plots of major myeloid lineage abundances (normalized to CD45+ cells) across clinical stages. Wilcoxon rank-sum test. **(D)** Heatmap of top marker genes distinguishing fine-grained myeloid subtypes. **(E)** Heatmap showing DecoupleR activation scores for 14 biological pathways across myeloid subtypes. **(F, G)** Dynamic shifts in myeloid subpopulation proportions across clinical stages tracked by an alluvial plot **(F)** and an aligned Ro/e enrichment heatmap **(G)**. **(H)** CytoTRACE2 differentiation potential mapped across myeloid sub-clusters. **(I)** Dot plot of BMDM phenotypic states categorized by monocyte, M2-like, and antigen-presentation markers. **(J)** Nebulosa kernel density estimation plots visualizing the expression weights of E-MDSC, Hypoxia-TAM, and GPNMB+ MDM signatures overlaid on the BMDM UMAP space. See also Figure S5.

High-resolution subclustering resolved 23 distinct myeloid subpopulations across these lineages (**Figure 3A**). Rather than uniform activation, these subsets exhibited highly polarized responses to immunotherapy (**Figure 3E-G**). ICB exposure drove widespread interferon-mediated JAK-STAT pathway activation across five distinct sub-lineages, signaling successful immune engagement (**Figure 3E**). However, therapeutic resistance was tightly anchored to specific non-responsive subsets. Crucially, intermediate monocytes (Mono.CD14.FCGR3A) were almost exclusively enriched in ICB+Target Non-responders, uniquely displaying hyperactivated oxidative stress, complement, and epithelial-mesenchymal transition (EMT) pathways (**Figure S3D**). Similarly, while most DC subsets correlated with treatment response, the mature DC3.LAMP3.CCR7 subset was paradoxically enriched in Non-responders, indicating a hijacked, immune-tolerant phenotype^24^. Overall, a clear lineage dichotomy emerged: MG and neutrophil subsets tracked with therapeutic response, whereas BMDM and monocyte progenitor subsets formed the core immunosuppressive architecture of non-response.

To trace the ontogeny of this immunosuppressive axis, CytoTRACE analysis was performed, revealing a clear developmental trajectory where peripheral monocytes (Mono) possess high differentiation potential, tracking into differentiation-arrested myeloid-derived suppressor cells (MDSCs; comprising E-MDSC and M-MDSC subsets) (**Figure 3H**). These arrested MDSCs were characterized by restricted maturation (low *CD14/FCGR3A*), impaired antigen presentation (*HLA-DRB1/CD74*), and a profound hypoxic signature (*ADM/HIF1A*) (**Figure 3I**). Cross-dataset benchmarking against external hypoxia-associated myeloid models—including Jackson et al.’s E-MDSCs^25^, Wang et al.’s Hypoxia-TAMs^26^, and Du et al.’s GPNMB+ MDMs^27^—confirmed precise transcriptional convergence (**Figure 3J**). The highest scores for all three external signatures mapped exclusively onto our hypoxic E-MDSC and M-MDSC populations. These results unify disparate nomenclatures in neuro-oncology, demonstrating that previously reported “Hypoxia-TAMs” and “GPNMB+ MDMs” are transcriptionally equivalent to the differentiation-arrested MDSCs driving ICB resistance.

### 1.4 Heterogeneous Lymphoid Responses: The Dual Roles of T Cells and NK Cells

Coarse-grained lymphoid profiling resolved seven major lineages, dominated by CD8+ T (42.0%), CD4+ T (26.7%), and NK cells (13.0%) (**Figure S4B**). Standard chemoradiotherapy left these baseline proportions unaltered. Conversely, ICB and combination ICB+Target interventions driven a significant expansion of both CD8+ T and NK cells within the immune macroenvironment, whereas B/Plasma cells selectively accumulated under combined ICB+Target pressure (**Figure S5C**). High-resolution subclustering partitioned these lineages into 36 distinct lymphoid subpopulations (**Figure S4C**). While cytotoxic transcripts (*IFNG/GZMB*) partially overlapped across clusters—reflecting a continuous evolutionary transition from effector CTLs to exhausted states—the atlas definitively captured terminally exhausted T (**Tex**) cells (**Figure S5E**).

Functional evaluation exposed a stark response-resistance dichotomy across these subsets (**Figure S4D-F**). Resistance to ICB+Target therapy was tightly coupled to the persistent accumulation of immunosuppressive Tregs, which uniquely hyperactivated hypoxic and inflammatory signaling cascades (*NFKB/TGFB*) to maintain themselves within immune-privileged niches. Paradoxically, B and plasma cell expansions preferentially tracked with Non-responders; our data suggest that this treatment-induced lymphoid aggregation manifests as immature, non-functional tertiary lymphoid structures (TLS) that accumulate suppressive myeloid cells rather than coordinating productive anti-tumor immunity^28^.

Effector dynamics further clarified clinical outcomes. Effector CTL subsets consistently enriched in Responders (**Figure S4E**). Crucially, **Tex** subpopulations exhibited divergent trajectories: the hypoxic Tex.03.ALDOA.MIF subset was exclusively enriched in Non-responders, representing a terminally dysfunctional, non-reinvigoratable pool arrested by microenvironmental hypoxia. In contrast, terminal Tex.02 cells enmeshed with Responders, suggesting these checkpoint-expressing populations represent plastic, “reinvigoratable” precursors responsive to immune blockade. Within the innate arm, NK cells exhibited the highest baseline cytotoxic program expression across the entire lymphoid compartment (**Figure S4H**). These NK subsets displayed intense response-dependent subset skewing driven by an NKG2A^high^CD16^low^ progenitor pool (**Figure S4G**), highlighting an essential, non-redundant innate effector axis required for driving immunotherapy efficacy.

### 1.5 Treatment-Driven Plasticity: Identifying Resistance-Associated Endothelial and Stromal Subpopulations

Coarse-grained annotation partitioned the non-immune microenvironment into EC and stromal compartments, subclassifying ECs into arterial, venous, capillary, and tip-like variants, and stromal components into smooth muscle cells (SMCs), pericytes (PCs), and CAFs (**Figure 4A, B**). Proportional quantification across cohorts revealed that despite active anti-angiogenic pressure, the CAF lineage was uniquely and significantly elevated in the rGBM-ICB+Target group compared to both pGBM and rGBM-R+C baselines (**Figure S6C**), an adaptive fibrotic response validated in external datasets (**Figure S6D**). High-resolution subclustering resolved these lineages into 10 EC and 8 Stromal subsets (**Figure 4C**), distinguishing three myofibroblastic CAF (myCAF) subsets—dominated by matrix-remodeling myCAF.01, matrix-stiffening myCAF.02.COL6, and collagen-producing myCAF.03—alongside antigen-presenting apCAF.CD74 and proliferative prolCAF.TOP2A.

**Figure 4.**
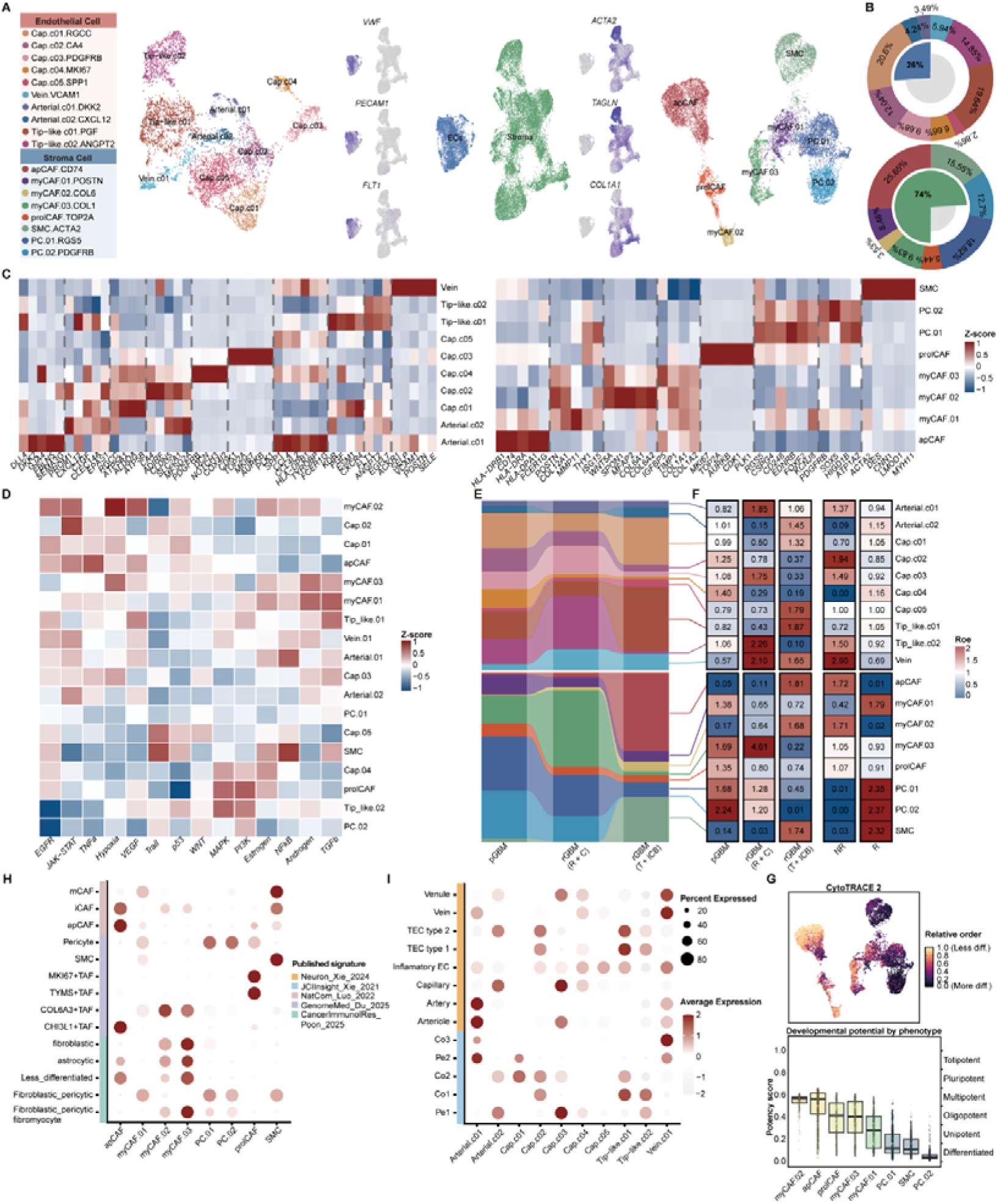
Remodeling of the Perivascular and Stromal Niche Under Therapeutic Pressure. **(A)** Landscape of the non-immune stromal compartment showing integrated UMAP clustering (center), feature plots of canonical lineage markers (inner flanks), and independent sub-clustering UMAPs for Endothelial (far left) and Stromal (far right) cells. **(B)** Nested donut charts showing major lineage (inner) and fine-grained subtype (outer) distributions for endothelial (top) and stromal (bottom) populations. **(C)** Heatmaps of top differentially expressed genes defining endothelial (left) and stromal (right) sub-clusters. **(D)** DecoupleR pathway activation scores highlighting functional heterogeneity across perivascular and stromal subsets. **(E, F)** Structural remodeling of the niche across treatment stages tracked via an alluvial plot **(E)** and an aligned Ro/e enrichment heatmap **(F)**. **(G)** CytoTRACE2 stemness scores and developmental potential quantification across stromal sub-clusters. **(H, I)** Cross-study validation matching identified Stromal **(H)** and Endothelial **(I)** sub-clusters with reference signatures using AUCell scores. See also Figure S6.

Functional profiling unmasked a paradoxical, highly heterogeneous neovascularization axis under combination therapy (**Figure 4D-F**). Despite anti-angiogenic targeted pressure, two neovascular EC subsets—Vein.VCAM1 and Arterial.01.DKK2—were paradoxically enriched in Non-responders, driven by hyperactivated EGFR, NFKB, and VEGF signaling. Benchmarking against external datasets confirmed that these populations match tumor-core inflammatory and peripheral immune-evasive vascular phenotypes, acting as critical nodes of therapeutic failure (**Figure 4G**). Furthermore, we identified two functionally divergent tip-like EC subsets: Tip_like.01.PGF.CXCR4 hyperactivated angiogenic machinery and surged following combination therapy, representing adaptive neovascular repair; conversely, Tip_like.02.ANGPT2 tracked with standard-treatment residual vessels and dominated in combination Non-responders, indicating inherent vascular resistance.

Stromal compartment dissection identified apCAF.CD74 and myCAF.02.COL6 as the twin drivers of microenvironmental therapeutic failure, with both significantly accumulating in combination Non-responders (**Figure 4H**). While apCAF.CD74 displayed immune-interactive characteristics, the Type VI collagen-secreting myCAF.02.COL6 subset exhibited profound hypoxic and TGF-beta signaling, orchestrating extensive vascular fibrosis and matrix stiffening. Trajectory mapping via CytoTRACE 2 revealed that myCAF.02.COL6 possessed the highest developmental and differentiation potential within the entire stromal compartment, outstripping even proliferative subsets (**Figure 4I**). This indicates that myCAF.02.COL6 acts as a highly plastic, treatment-selected stromal progenitor. Rather than a passive architectural bystander, this population is selectively maintained under therapeutic pressure to build the refractory fibrotic scaffold that anchors GBM resistance.

### 1.6 Spatial Transcriptomics Reveals a cNMF7-Centric Niche within Hallmark Hypoxic Regions of Glioblastoma

To characterize the spatial architecture anchoring the aggressive malignant cNMF7 subpopulation, we integrated spatial transcriptomic data from 48 glioblastoma sections. Following cell-type deconvolution, spatial relationship modeling identified three highly co-localized cellular modules tightly synchronized with cNMF7 distribution (**Figure 5A, Figure S7A, Table S5**). Given the profound hypoxic profile of cNMF7 cells, we tracked their positioning relative to two classic histopathological pillars of severe tissue hypoxia: MVP and PAN.

**Figure 5.**
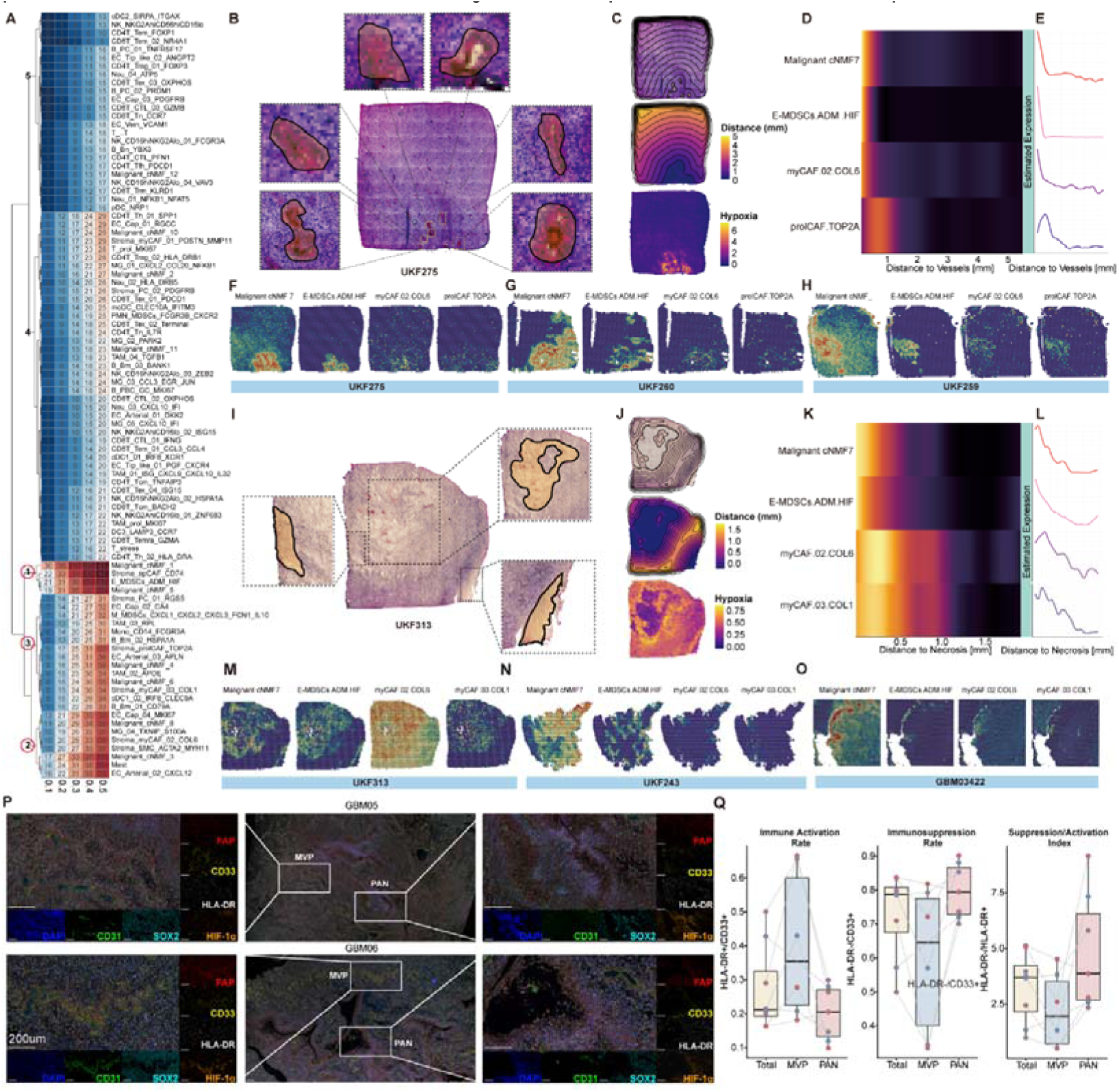
Spatial Phylogeography Defines a Recurrent “Spatial Resistance Triad” Within Hypoxic Niches. **(A)** Meta-analysis heatmap of spatial co-localization recurrence between cNMF7 and other cell types across 48 Visium sections, with cell types hierarchically clustered into 5 distinct spatial modules based on MISTy importance scores. **(B)** Representative H&E staining of section UKF275 with high-magnification views of annotated microvascular proliferation (MVP) zones. **(C)** SPATA2 trajectory analysis modeling the MVP niche: surface plot of annotated regions (top), spatial distance gradient map (middle), and spatial projection of hypoxia signature scores along the gradient (bottom). **(D, E)** Distance-dependent abundance profiling from the MVP core visualized via a gradient heatmap **(D)** and trajectory line plots **(E)** for cNMF7, E-MDSC.ADM.HIF1A, myCAF.02.COL6, and prolCAF.MKI67 cells. **(F–H)** RCTD-deconvoluted spatial feature plots of the triad components across three independent MVP-containing Visium samples. **(I)** Representative H&E staining of section UKF313 highlighting perinecrotic pseudopalisading (PAN) zones. **(J)** SPATA2 trajectory analysis modeling the PAN niche relative to the necrotic core. **(K, L)** Distance-dependent abundance profiling from the necrotic edge visualized via a gradient heatmap **(K)** and trajectory line plots **(L)** for the core triad members. **(M–O)** RCTD spatial feature plots validating triad component distribution across three independent PAN-containing Visium sections. **(P)** Multiplex IHC scans validating the physical co-localization of FAP, CD33, HLA-DR, CD31, SOX2, and HIF-1α within MVP and PAN regions. Scale bar = 200 um. **(Q)** Quantitative evaluation of regional myeloid phenotypes across global tissue areas, MVP, and PAN mapped via mIHC, showing box plots of the Myeloid Immune Activation Rate (HLA-DR positive cells divided by CD33 positive cells), Myeloid Immunosuppression Rate (HLA-DR negative cells divided by CD33 positive cells), and Myeloid Suppression / Activation Index (HLA-DR negative cells divided by HLA-DR positive cells). Individual paired lines track distinct patient specimens. See also Figures S7, S8, and S9.

Investigation of the MVP niche within representative section UKF275 confirmed these histopathological zones as the most severely hypoxic regions in the tissue (**Figure 5B, C**). Spatial gradient analysis across annotated MVP zones demonstrated that the highest proportions of cNMF7 cells, E-MDSC.ADM.HIF1A, myCAF.02.COL6, and prolCAF.MKI67 precisely anchored within the MVP core, with their presence dropping sharply as a function of distance (**Figure 5D-F, Figure S7B**). This localized spatial triad was independently validated across multiple additional MVP-containing patient sections (**Figure 5G, H**). Parallel assessment of the PAN feature in section UKF313 confirmed that the pseudopalisading boundaries enveloping necrotic cores represented the primary hypoxic epicenters (**Figure 5I, J**). Spatial gradient mapping showed that cNMF7 malignant cells, E-MDSC.ADM.HIF1A, myCAF.02.COL6, and myCAF.02.COL1 concentrated heavily within the PAN zone, decaying progressively into the broader tumor parenchyma (**Figure 5K-M, Figure S7C**). This spatial recruitment pattern was highly reproducible in independent PAN validation cohorts (**Figure 5N, O**).

To overcome pixel-resolution constraints and validate this cellular architecture at single-cell protein resolution, we performed multiplex immunohistochemistry (mIHC) on an independent clinical glioblastoma cohort (**Figure 5P**). In situ fluorescence imaging confirmed the dense structural alignment of FAP+ CAFs, SOX2+CD44+ mesenchymal-like malignant cells, and CD33+HLA-DR- MDSCs. Functional mapping of niche-associated myeloid cells exposed a stark divergence in polarization between these microenvironments (**Figure 5Q**). Myeloid cells within the hypoxic PAN zones exhibited a profoundly immunosuppressive phenotype, defined by an elevated Myeloid Immunosuppression Rate (HLA-DR-/CD33+) and a skewed Myeloid Activation/Suppression Index. Conversely, the hypervascular MVP niches demonstrated a dominant Myeloid Immune Activation Rate (HLA-DR+/CD33+).

These multi-omic findings demonstrate that while both hypoxic MVP and PAN niches recruit the core components of this spatial resistance triad, they impose distinct functional polarization states on the infiltrative myeloid compartment to drive therapeutic failure.

### 1.7 Key Communication Axes: Collagen/Fibronectin-CD44 Drive myCAF Crosstalk with cNMF7 and E-MDSC

Integrated spatial transcriptomic analysis established ECM-secreting myCAFs as a foundational third compartment co-localizing with cNMF7 malignant cells and E-MDSCs within the histopathological structures of MVP and PAN (**Figure 5**). Intercellular communication modeling revealed that this tripartite niche is anchored by a dominant, myCAF-initiated network where myCAF-derived Collagen I, Collagen VI, and Fibronectin (FN1) preferentially engage the CD44 receptor expressed concurrently on cNMF7 cells and E-MDSCs (**Figure S8A, B**). Spatial mapping confirmed that these ligand-receptor pairs exhibit intense, overlapping transcript expression restricted within both MVP and PAN boundaries, establishing the myCAF-driven matrix as a primary signaling conduit rather than passive structural support (**Figure S8D, E**).

Beyond structural matrix wiring, paracrine and juxtacrine mapping showed that myCAFs deploy Macrophage Migration Inhibitory Factor (MIF) and surface APP toward E-MDSCs, while secreting Pleiotrophin (PTN) and utilizing homophilic CD99 contact to signal to cNMF7 cells (**Figure S7D, E, Figure S8C**). Intriguingly, while MIF transcripts were enriched in both MVP and PAN zones, its cognate receptors (CXCR4/CD74) lacked clear spatial co-localization (**Figure S7F**). This structural mismatch suggests a non-canonical or localized inefficiency for soluble MIF signaling, contrasting with the continuous, high-fidelity deployment of the core Collagen/FN-CD44 axis.

This multicellular crosstalk is highly reciprocal and reinforced by feedback loops from the microenvironment (**Figure S8F, G**). E-MDSCs signal back to myCAFs via Secreted Phosphoprotein 1 (SPP1), while cNMF7 cells leverage their mesenchymal state to secrete TNC, LAMB1, and FN1 toward myCAF integrins and CD44 (**Figure S7G, H**). Crucially, high-resolution spatial mapping unmasked distinct regulatory environments between the two hypoxic domains: the SPP1, TNC, and PTN feedback loops and their cognate receptors demonstrated profound, specific spatial co-localization within hypervascular MVP zones, but faded significantly within perinecrotic PAN regions (**Figure S8H, I, Figure S7I**). This heterogeneity indicates that while the foundational Collagen/FN-CD44 matrix axis forms a conserved microenvironmental shield, the accessory paracrine regulatory mechanisms are niche-specific.

### 1.8 Clinical Relevance and Pan-Cancer Validity of the myCAF-ECM-CD44 Axis

Single-cell baseline profiling in the GRIT-Atlas confirmed that collagen expression is strictly stromal-restricted and *FN1* spans both stromal and endothelial compartments, while their cognate receptor *CD44* is ubiquitously expressed across the microenvironment (**Figure S8A**). Interrogation of our bulk RNA-seq cohort confirmed that tissue expression of these six core ligands is significantly elevated in glioblastoma compared to normal brain and directly correlates with advancing tumor grade (**Figure S8B**). Survival stratification across seven independent patient cohorts revealed that hyperactivation of this ECM module consistently predicts poor prognosis and tracks with a reduced objective response rate to immunotherapy (**Figure S8C, D**).

Spatial mapping using the Ivy Glioblastoma Atlas Project confirmed that all six ligands peak within peri-vascular regions with secondary enrichment in peri-necrotic zones, providing independent histopathological validation for our MVP and PAN niche models (**Figure S8E**). Transcriptional correlation analysis demonstrated that this myCAF-driven ECM signature strongly synchronizes with hallmark microenvironmental programs—including hypoxia, EMT, and angiogenesis—while positively correlating with molecular readouts of M2-like TAMs, Tregs, MDSCs, and T-cell exhaustion (**Figure S8F**). Pan-cancer transcriptomic and proteomic evaluation via TCGA, CPTAC, and GTEx datasets established this axis as a highly conserved hallmark of human malignancy, correlating with poor overall survival in over two-thirds of cancer types (**Figure S8G**). Proteomic validation confirmed universal overexpression of these core matrix proteins in both clear cell renal cell carcinoma and glioblastoma. These clinical findings transition the myCAF-ECM-CD44 axis from a localized glioblastoma phenomenon into a generalized, clinically actionable paradigm of microenvironmental therapy resistance.

### 1.9 High-Plex Single-Cell Spatial Imaging Resolves the Sub-Class Architecture and Neighborhood Geography of Mesenchymal Malignant Niches

High-plex spatial molecular imaging of primary glioblastoma specimens mapped 406,689 single cells into 11 major cell lineages, capturing the global ecosystem at absolute single-cell resolution (**Figure 6A-C, Figure S10A-F**). To define how the localized microenvironment shapes malignant states, we performed spatial neighborhood analysis centered on the MES-like subpopulation (**Figure 6A**). Matrix factorization and hierarchical clustering resolved three distinct, reproducible “MES Niches” with highly specialized transcriptomic identities and divergent cellular architectures (**Figure 6D-H, Table S8**). Differential expression analysis of the imaging data revealed that these niches represent distinct functional states across the hypoxic and structural spectrum: The Ischemic/PAN Core (niche s1) is Driven by an intense anaerobic glycolytic shunt and terminal ischemic survival program, defined by severe hypoxia markers (*VEGFA, NDRG1, HILPDA, ADM*), glucose transporters (*SLC2A1, SLC2A3*), and rate-limiting glycolytic enzymes (*LDHA, PGK1, ENO1, PFKFB4, HK2*). The Immune-Infiltrated/Reactive Niche (niche s2): Exhibiting an active inflammatory and reactive astrocytic response network, marked by *FABP7, CHI3L1, CLU*, and *STAT3*, alongside crucial basement membrane structural components (*COL6A1, COL6A2*). The Hypervascular/Fibrotic MVP Niche (niche s3) is characterized by robust stromal-remodeling, immune-evasion, and matrix-receptor signaling, expressing top-tier desmoplastic elements (*TNC, SPP1, COL1A1/A2, COL3A1*), antigen-presentation/checkpoint partners (*CD74, HLA-DRA, CD274/PD-L1, NT5E/CD73*), and the master matrix receptor *CD44*. In situ single-cell coordinate mapping directly validated the physical authenticity of these environments (**Figure 6I, J**). Niche s1 anchored PAN structures, forming dense clusters of MES-like cells tightly interspersed with MDSCs (**Figure 6F, K**). Niche s2 manifested as an intermingled macrophage-microglia network undergoing a localized inflammatory response (**Figure 6F, K**). Conversely, niche s3 anchored MVP regions, tracing organized perivascular cuffs where CAFs wrapped tightly around the endothelial lining to drive a robust stromal-remodeling and ECM scaffolding program (**Figure 6F, K**).

**Figure 6.**
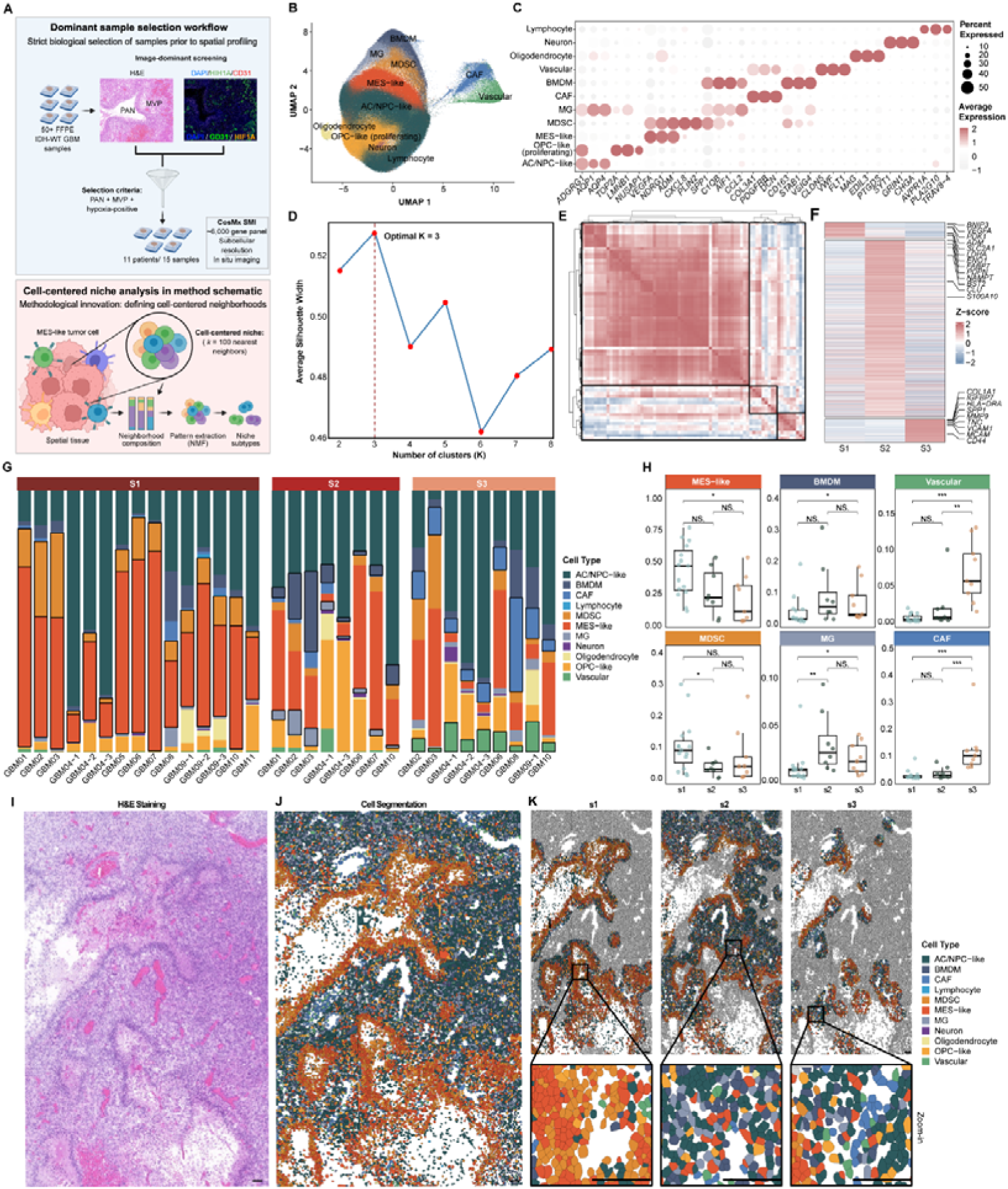
High-Plex Spatial Molecular Imaging and Microenvironmental Neighborhood Analysis of Mesenchymal Malignant Niches. **(A)** Schematic diagram of the CosMx 6k Spatial Molecular Imager (SMI) profiling pipeline (upper) and the unsupervised 100-nearest-neighbors spatial neighborhood analysis framework centered on MES-like cells (lower). **(B)** Global UMAP embedding of 406,689 single cells resolved into 11 major lineages. **(C)** Dot plot of characteristic marker genes across the identified single-cell lineages. **(D, E)** Determination of optimal MES niches shown by average silhouette width optimization **(D)** and a pairwise Pearson correlation matrix of neighbor cell compositions **(E)**. **(F)** Heatmap displaying differentially expressed genes across global MES niches s1, s2, and s3, with representative signature genes highlighted. **(G)** Stacked bar plots showing the compositional distribution of MES niches across individual patient samples. **(H)** Box plots of the proportional frequency of dominant neighborhood cell types defining niches s1, s2, and s3. **(I, J)** Representative H&E tissue overview of sample GBM02 **(I)** and its corresponding cell-type annotation map at absolute single-cell segmentation resolution **(J)**. Scale bars = 125 μm. **(K)** Spatially resolved cellular mapping of localized niches s1, s2, and s3. Top panels show global tissue distribution with background cells masked; bottom panels show high-magnification zoom-in views highlighting the unique multicellular architecture of each niche. Scale bars = 125 μm. See also Figure S10.

Crucially, physical distance modeling relative to macroscopic pathological landmarks revealed a rigid, continuous spatial gradient across the entire cohort: the hypervascular niche s3 was uniformly positioned closest to MVP cores, whereas niche s1 mapped to distant, perinecrotic domains (**Figure S10H**). Rather than representing static, isolated structures, this spatial continuum unmasks a spatiotemporal evolution of GBM pathology wherein PAN (niche s1) emerges as the direct pathophysiological consequence of dysfunctional MVP (niche s3). The chaotic and hyperpermeable vessel networks within the MVP-anchored niche s3 inevitably undergo structural failure, perfusion decay, and thrombosis. This vascular collapse precipitates acute tissue ischemia, a catastrophic transition captured transcriptomically by a metabolic shift from active HIF1A-driven neo-angiogenesis in s3/s2 to the chronic hypoxic coordinator *EPAS1 (HIF2A)* and downstream cell-death executioners (*BNIP3, BNIP3L, DDIT3*) in the perinecrotic niche s1 boundary. Importantly, this evolutionary trajectory uncovers the omnipresent and continuous nature of the COL6A1–CD44 resistance axis across the entire MVP-PAN continuum. While the mechanical ligand *COL6A1* is dynamically deposited within the reactive s2 infrastructure and its cognate receptor *CD44* is heavily enriched within the perivascular s3 cuffs at active MVP sites, their structural cross-talk does not vanish during vascular decay. Instead, as the microenvironment transitions into the ischemic PAN core (s1), the highly stable, degraded ECM tracks rich in COL6A1 persist within the scar tissue, continuously engaging with CD44-expressing migrating MES-like malignant cells flanking the necrotic borders. This reveals that the glioblastoma resistance triad is organized into anatomically precise niches that chronicle the inevitable evolution from vascular proliferation to tissue necrosis, utilizing the COL6A1–CD44 axis as a continuous mechanical sanctuary to sustain tumor stemness throughout the lifecycle of hypoxia.

### 1.10 Mechanistic Disruption of the COL6A1-CD44 Axis and Preclinical Efficacy of CNS-Penetrant Triad-Targeting Therapies

Exogenous ECM ligand screening in patient-derived glioma stem cells (GSCs) demonstrated that matrix components drive an overall trend toward upregulating core neural stemness markers (SOX2 and CD133), with recombinant COL6A1 (rCOL6A1) emerging as the most robust and consistent driver of this stemness cascade (**Figure S11A-C**). Furthermore, while all other tested matrix components successfully bypassed the CD44 blockade to trigger a significant surge in SOX2 expression, the stemness-inducing capacity of rCOL6A1 was completely and selectively abolished across both GSC01 and GSC06 lines (**Figure 7A-C, Figure S11D, E**). This selective vulnerability to receptor neutralization confirms that while other desmoplastic elements can engage alternative surface receptors, COL6A1 exhibits a strict functional dependency on the CD44 receptor to sustain the glioblastoma stemness program. Histological and immunofluorescence stratification of clinical glioblastoma cohorts confirmed that high-COL6A1 expression strongly tracks with severe tissue hypoxia and a profound enrichment of MVP and PAN niches (**Figure 7D, Figure S11F**). Multiplex in situ protein profiling validated the precise physical convergence and structural alignment of the FAP+/COL6A1+/CD44+ resistance triad surrounding these hallmark vascular and necrotic structures (**Figure 7E-G**).

**Figure 7.**
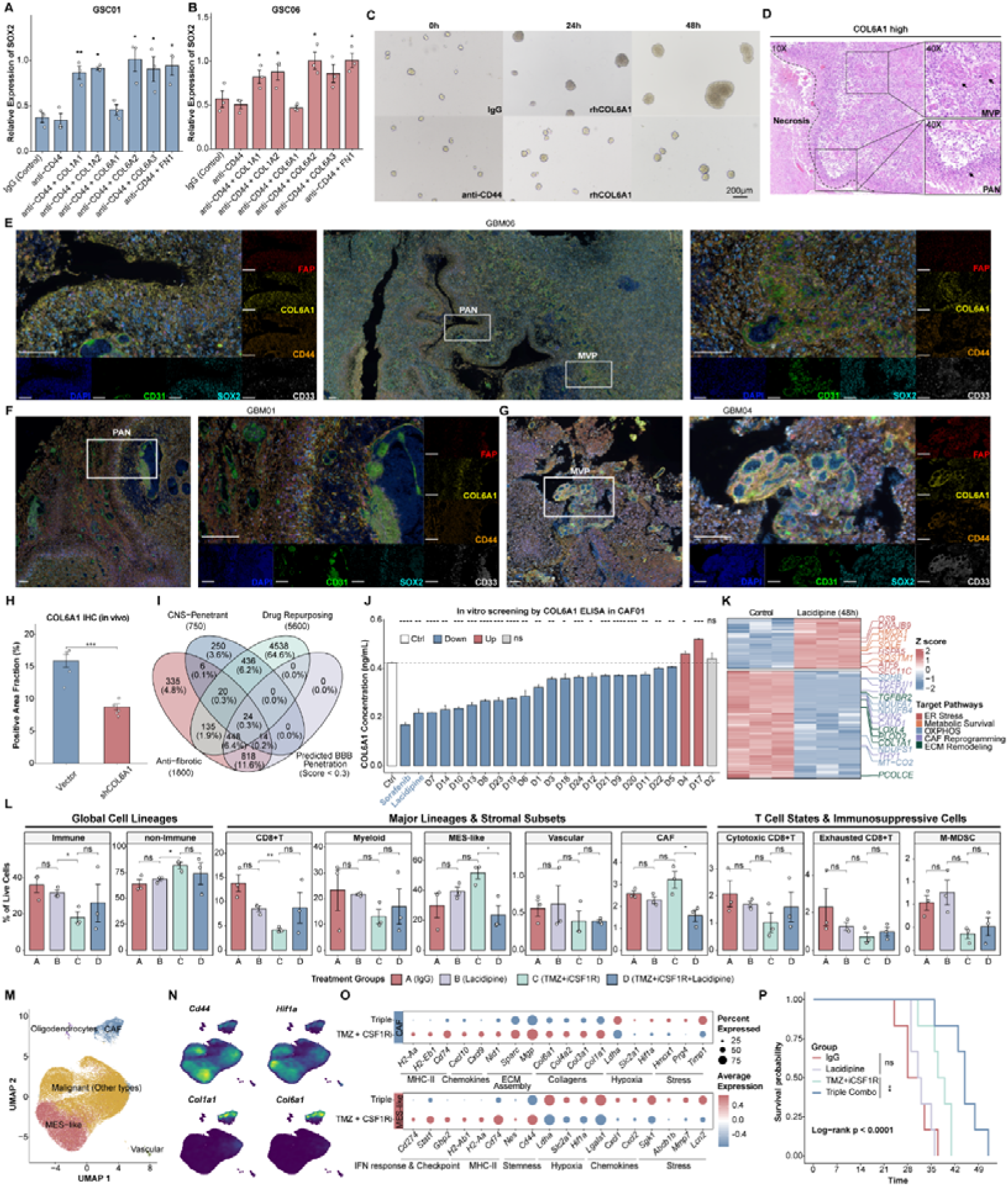
Mechanistic Disruption of the COL6A1-CD44 Axis and Preclinical Efficacy Evaluation of CNS-Penetrant Compounds in Orthotopic Glioblastoma Models. **(A, B)** Western blot quantification of SOX2 expression in primary GSC01 **(A)** and GSC06 **(B)** lines following 24-hour challenge with six candidate recombinant ECM ligands, with or without anti-CD44 neutralizing antibody pre-treatment. **(C)** Phase-contrast micrographs tracking longitudinal in vitro oncosphere formation kinetics under rhCOL6A1 stimulation combined with control IgG or anti-CD44 neutralization. **(D)** H&E overview of clinical specimens stratified by high COL6A1 expression, showing prominent PAN and MVP structures. **(E–G)** Multiplex IHC scans tracking FAP, COL6A1, CD44, CD31, SOX2, and CD33, confirming direct cell-to-cell co-localization of the triad within PAN and MVP niches across clinical specimens. **(H)** Bar plot quantifying intra-tumoral Col6a1 positive area fractions (%) in orthotopic models derived from shRNA-Col6a1 versus non-targeting control GL261 cells at day 14. **(I)** Four-way Venn diagram of the integrated compound screening workflow filtering the CNS-Penetrant, Drug Repurposing, and Anti-Fibrosis libraries via BBB penetration probabilities. **(J)** ELISA quantification of extracellular COL6A1 secretion in primary CAF01 cells across candidate screening compounds. **(K)** Hierarchical clustering heatmap of 4D-DIA quantitative proteomics profiles in CAF01 cells following Lacidipine versus vehicle treatment, annotated by core functional pathways. **(L)** Full-spectrum flow cytometry quantification of intra-tumoral immune and non-immune lineages across the four in vivo therapeutic arms. **(M, N)** Single-cell RNA-seq profiling of the isolated CD45- non-immune fraction from orthotopic tumors, shown by global UMAP embedding **(M)** and single-cell feature plots of *Cd44, Hif1a, Col1a1*, and *Col6a1* expression intensities **(N)**. **(O)** Comparative dot plot showing single-cell transcriptional shifts and pathway activation between the Doublet (TMZ + anti-CSF1R) and Triple-combination (Doublet + Lacidipine) cohorts within isolated CAF and MES-like sub-clusters. **(P)** Kaplan-Meier overall survival curves of orthotopic GL261-Luc tumor-bearing mice across the four therapeutic regimens. Log-rank test. See also Figures S11 and S12.

Comparative profiling of primary patient lines and tumor-autonomous *Col6a1* knockdown in orthotopic murine models revealed that stromal CAFs, rather than malignant cells, serve as the primary engine driving global microenvironmental COL6A1 deposition (**Figure 7H, Figure S11G-I**). To exploit this architectural dependency, we performed an ELISA screening across 24 candidate compounds to identify small-molecule disruptors of extracellular matrix secretion (**Figure 7I**). In CAF01 cells, Sorafenib and Lacidipine emerged among the top matrix-suppressing candidates (**Figure 7J**). Parallel screening in CAF06 consistently validated these results, ranking both Lacidipine and Sorafenib within the top five inhibitors of extracellular COL6A1 secretion (**Figure S11J**). Although Sorafenib demonstrated potent matrix suppression across both primary patient backgrounds, its clinical utility in glioblastoma has been historically limited by severe dose-limiting toxicities and a lack of sustained efficacy in patient trials^29,30^. Conversely, Lacidipine—a clinically approved, exceptionally well-tolerated antihypertensive agent with high lipophilicity—exhibited robust, reproducible COL6A1 suppression and an established capacity to mitigate tissue fibrosis^31^. Consequently, Sorafenib and Lacidipine were identified as the primary candidate choices, with Lacidipine selected as the optimal repurposing candidate to dismantle the tumor-shielding desmoplastic niche. Quantitative proteomic dissection of Lacidipine-treated CAFs unmasked a multi-modal stromal disruption program, characterized by the systematic downregulation of oxidative phosphorylation machinery, the silencing of CAF activation markers, and the comprehensive collapse of the extracellular matrix remodeling cascade (**Figure 7K, Table S10**).

In vivo functional validation in orthotopic murine models demonstrated that adding Lacidipine to standard chemo-immunotherapy (TMZ and anti-CSF1R) induces a sustained, long-term suppression of intracranial tumor burden, whereas the standard doublet backbone fails (**Figure S12A-C**). To dissect the cellular rearrangements driving this synergy, full-spectrum flow cytometry was performed across the treatment cohorts (**Figure 7L**). While the standard doublet (TMZ+iCSF1R) failed to eradicate protective niches—instead expanding the global non-immune compartment via the accumulation of therapy-resistant MES-like cells and CAFs alongside a reduction in CD8+ T cells—the triple combination incorporating Lacidipine selectively dismantled this resistance architecture. Compared directly to the doublet, Lacidipine precipitated a significant collapse of both the MES-like malignant subpopulation and the CAF compartment, effectively neutralizing the cellular engines of the desmoplastic niche (**Figure 7L**). Mechanistically, this therapeutic synergy forces the complete transcriptional silencing of core structural collagens and immunomodulatory chemokines within the residual CAF sub-cluster, while inducing severe microenvironmental stress in MES-like cells that suppresses canonical stemness determinants and immune checkpoint elements (**Figure 7M-O, Figure S12D-E, Table S11**). Ultimately, dismantling this biophysical fibrotic shield translated into a dramatic extension in overall survival (**Figure 7P**), establishing matrix-targeted disruption as a prerequisite to overcome glioblastoma resistance.

## Discussion

The catastrophic failure of ICB in phase III clinical trials for GBM underscores a critical structural blind spot: we have long conceptualized therapeutic resistance as a cell-intrinsic or spatially disordered phenomenon, rather than an evolutionarily engineered ecosystem. By construction of the GRIT-Atlas—encompassing nearly one million high-quality cells across 296 clinical specimens—we provide the definitive spatial and evolutionary blueprint of IDH-wildtype GBM under multi-modal treatment pressures. Beyond offering a static census of cell states, our study exposes a convergent evolutionary trajectory where diverse therapeutic stresses select for a highly aggressive MES-like malignant state. Crucially, by cross-linking macro-scale spatial transcriptomics with subcellular molecular imaging, we define a physical fortress protecting these cells—the “Spatial Resistance Triad”—comprising MES-like malignant cells, MDSCs, and CAFs. This desmoplastic sanctuary, physically anchored within the hypoxic niches of MVP and PAN, establishes an impermeable barrier to T cell infiltration, demonstrating that successful immunotherapy is fundamentally contingent upon the prior dismantling of this stromal architecture.

The remarkable plastic potential of malignant cells represents the primary engine of GBM recurrence. Our transcriptional network analysis demonstrates that while SOC chemoradiotherapy enriches for neuron-like states—likely hijacking synaptic integration to survive clearance^6—the addition of ICB and anti-angiogenic combinations forces a dramatic, non-responsive Darwinian selection toward the MES-like state. This cell state is defined by a potent, synergistic intersection of CD44-mediated stemness, profound tissue hypoxia, and hyperactivated NFκB/TNF inflammatory signaling. Our findings clarify that the “mesenchymal” subtype historically defined by bulk transcriptomics is not a monolithic clonal entity, but rather a spatially organized, stress-adaptive, drug-tolerant persister compartment. The exclusive hyper-enrichment of this specific MES-like state in targeted-and immuno-therapy non-responders implies that therapeutic paradigms must urgently pivot away from broad-spectrum cytostatic agents toward selective interventions targeting the upstream epigenetic or transcriptional master regulators that lock malignant cells into this lethal, immune-evasive phenotype.

A long-standing controversy in neuro-oncology concerns the exact identity and ontological origins of immunosuppressive myeloid cells within the GBM microenvironment: whether they represent classical M2-polarized tissue-resident macrophages or distinct bone marrow-derived developmental lineages. Our high-resolution single-cell mapping resolves this debate by clearly separating immunosuppressive MDSCs from canonical M2-like TAMs. We demonstrate that the highly hypoxic, immunosuppressive myeloid populations previously described under disparate nomenclatures—such as “Hypoxia-TAMs,” “GPNMB+ MDMs,” or “E-MDSCs”—are transcriptionally convergent and represent a single functional entity: differentiation-arrested, bone marrow-derived MDSCs. These data strongly argue against the simplistic, linear M1-to-M2 macrophage polarization model. Instead, we propose a “Maturation Arrest” model, wherein severe localized hypoxia within the recurrent niche acts as a developmental block, preventing monocyte-to-macrophage differentiation and locking these infiltrating myeloid cells into a permanent, highly potent immunosuppressive MDSC state. Consequently, myeloid-targeted therapies should abandon attempts to “re-polarize” mature TAMs, and instead focus on forcing the differentiation or targeted elimination of these arrested MDSCs by disrupting their localized survival signals within the hypoxic sanctuary.

The most profound conceptual advance of our study is the precise spatial definition and mechanistic dissection of this resistance niche, which effectively unifies and refines previously conflicting microenvironmental models. Prior studies have captured fragmented glimpses of these cellular relationships; for instance, Jain et al. demonstrated the co-localization of CAFs with mesenchymal GSCs, endothelial cells, and M2 macrophages in the representative UK275 section^32^, while Du et al. suggested a distinct mechanism where collagen-expressing CAFs, typically situated in the MVP area, secrete humoral factors to program macrophages into immunosuppressive phenotypes found in the PAN area^27^. Additionally, Poon et al. utilized digital spatial profiling of the perivascular region and identified vascular/perivascular stromal cells (PVSCs)—which our trajectory analysis links to a high-differentiation-potential CAF subset—co-localizing with HLA-DRA^low^ inhibitory monocytes and resident perivascular macrophages^15^. While these studies established a loose consensus around a stromal-myeloid-stem cell axis in MVP or PAN regions, they diverged significantly on cellular nomenclature and exact topological boundaries. Our data integrate these disparate observations into a unified framework. We confirm that a specialized subset of collagen-secreting CAFs acts as the primary architectural driver within these domains. Crucially, our study uncovers that this stromal subset constructs a dense, protective physical and signaling scaffold via the COL6A1-CD44 axis, directly securing the spatial trapping of MDSCs and the maintenance of MES-like malignant stemness, thereby erecting a definitive mechanical shield that excludes cytotoxic T cells.

Importantly, by leveraging high-plex spatial molecular imaging via the CosMx SMI to map over 400,000 single cells, we transcend the limitations of static spatial co-localization to unmask a dynamic pathobiological continuum. While previous low-resolution spatial studies treated MVP and PAN as distinct, isolated morphological landmarks, our sub-cellular gradient evidence reveals a spatiotemporal continuum wherein PAN emerges as the direct, inescapable functional consequence of progressive MVP advancement and subsequent localized microvascular collapse. The collagen-secreting CAFs within the MVP niche do not merely sit passively alongside vessels; they drive a fibrotic remodeling event—principally mediated by CAF-derived COL6A1—that remodels the ECM. This extensive desmoplastic deposition engages the CD44 receptors heavily upregulated on recipient cells in these regions^33–35^, accelerating a neural stemness cascade while simultaneously choking tissue perfusion. This mechanical and hypoxic stress culminates in vascular collapse, directly generating the peri-necrotic zones that define PAN, where COL6A1 expression is significantly enriched^36,37^. By identifying COL6A1 as the central structural lynchpin of this MVP-to-PAN transition, our work provides a mechanical rationale for the spatial transition of the tumor tissue during therapeutic resistance, wherein CAF-derived FN1 further coordinates malignant cell migration and invasion^38^.

Ultimately, the discovery of the Spatial Resistance Triad fundamentally reorganizes the translational paradigm for recurrent glioblastoma. Attempting to reinvigorate exhausted T cells via systemic immune checkpoint monotherapy is fundamentally futile when those effectors are mechanically and chemically barred from entering the core resistance niches. Overcoming this intractability demands a rational, multi-pronged combinatorial strategy designed to breach the desmoplastic fortress: (1) Dismantling the Structural Scaffold via selective inhibition of the CAF-mediated COL6A1-CD44 matrix axis; (2) Alleviating Niched Hypoxia through precise vascular normalization to starve the hypoxic drive of MDSCs; and (3) Unlocking Myeloid Differentiation Arrest to neutralize the immunosuppressive core. Our identification of the BBB-penetrant small molecule Lacidipine as a potent stromal disruptor provides a clinically actionable translation of this strategy. By combining Lacidipine with standard chemo-immunotherapy (TMZ + anti-CSF1R), we achieved the complete collapse of this desmoplastic barrier, converting an impermeable “cold” sanctuary into an immune-accessible microenvironment, and translating into a profound therapeutic eradication of tumor burden and extended survival in preclinical models. The GRIT-Atlas thus provides the essential spatial blueprint required to dismantle the structural foundations of glioblastoma resistance.

### Limitations

We acknowledge several methodological and biological limitations that contextualize our conclusions. First, while the cumulative cellular volume across our single-cell (GRIT-Atlas) and spatial imaging (CosMx SMI) cohorts is unprecedented, the clinical sample size for highly specialized, treated, or recurrent specimens remains intrinsically constrained by the rarity of these surgical tissues. Second, our multi-platform integration introduces inherent technological dependencies. While snRNA-seq exhibits nuclear isolation biases toward glial and neuronal lineages, the CosMx SMI platform relies on a targeted high-plex gene panel rather than an unbiased whole-transcriptome readout. Furthermore, the chaotic and dense cellular architecture characteristic of glioblastoma MVP and PAN niches introduces unavoidable cell-segmentation challenges typical of in situ spatial imaging. Third, our small-molecule screening and translational validation framework remains relatively preliminary. The in vitro matrix-suppression screen utilized only a single initial concentration for candidate selection, and our subsequent in vivo therapeutic evaluation relied strictly on a fixed dosage referenced from previous literature, resulting in a simplified pharmacological characterization of Lacidipine’s efficacy. Although these models successfully demonstrated the functional causality of the COL6A1–CD44 axis, a more comprehensive exploration of dose-response dynamics, optimal therapeutic windows, and long-term systemic effects is required before translating this desmoplastic disruption strategy into clinical trials.

## Methods

### Human subjects

Patients were prospectively enrolled into this study at First Affiliated Hospital of Soochow University. Written informed consent was provided by all participants according to approved protocol. Fresh tumor tissue was collected at the time of surgery after confirmation of primary glial neoplasm on frozen section. None of the patients was treated with chemotherapy or radiation before tumor resection. Tumors were categorized according to the 2021 WHO Classification of Central Nervous System Tumors based on the combination of relevant histopathologic and molecular features from IHC.

### Treatment-Based IDH-WT GBM Classification

The standard treatment paradigm for IDH-wildtype GBM, classified as primary GBM (pGBM), involves maximal safe resection followed by radiotherapy with concurrent and adjuvant temozolomide, yet disease recurrence inevitably occurs at a median of 7 months post-diagnosis^39,40^. In current study, recurrent cases segregated into three treatment-defined categories: (1) post-Stupp protocol recurrences (rGBM-R+C); (2) PD-1 inhibitor monotherapy-treated cases (rGBM-ICB), including pembrolizumab (200mg administered 14±5 days pre-resection; Lee et al.^12^) and nivolumab (single preoperative dose; Antunes et al.^14^); and (3) combination therapy cases (rGBM-ICB+Target) featuring PD-1 blockade plus anti-angiogenic agents like anlotinib (12mg daily for 1 week until 2 weeks pre-resection; Mei et al.^13^). Through serial MRI monitoring, we identified responders (demonstrating ≥3 months of tumor stability/regression; n=7) versus non-responders (n=5) in neoadjuvant cohorts, establishing a framework that enables precise molecular characterization of treatment-specific evolutionary patterns and resistance mechanisms in recurrent GBM.

### Single-cell RNA-seq data preprocessing

Newly generated single-cell sequencing data were aligned with the GRCh38 human reference genome and quantified using Cell Ranger (version 3.0, 10x Genomics). The preliminary filtered data generated from Cell Ranger were used for downstream filtering and analyses. The quality of cells was then assessed based on two metrics: (1) The number of detected genes per cell; (2) The proportion of mitochondrial gene (Genes starting with ‘MT-’) counts per cell; (3) The proportion of ribosomal gene (genes starting with ‘RPS’ or ‘RPL’) counts per cell; (4) The proportion of hemoglobin gene (genes starting with ‘HB’ but are not followed by ‘P’) counts per cell. Specifically, cells with detected genes fewer than 500 or more than 5000, mitochondrial unique molecular identifier (UMI) count percentage larger than 10%, lysosomal UMI count percentage larger than 50%, and hemoglobin UMI count percentage larger than 0.05% were filtered out. To remove the potential doublets, Scrublet^41^ was used for each sequencing library with the expected doublet rate set to be 0.05, and cells with the predicted doublet Score larger than 0.3 were further filtered out. Finally, we excluded samples with fewer than 500 retained cells. For scRNA-seq data from other publications, the same filtering steps were applied for the datasets with raw count matrix to obtain high-quality cells. For the specific datasets with no available count data^42–46^, the CPM or TPM matrix were downloaded and used directly. However, the subsequent integration process showed that integrating these datasets into the atlas would make it hard to fix the overall batch effect, so these data weren’t included in the final atlas. The detailed metadata (including tissue location, sample and patient identifier) were retrieved from the original studies (**Table S1**).

### Batch effect correction

We next normalized each strictly filtered count data using the *scanpy.normalize_total* function with parameter ‘*target_sum=1e4*’ in Scanpy^47^ (v1.10.1). All the normalized data were logarithmically transformed for downstream analyses. The unexpected effects of the total counts and the original identify were regressed out from the normalized expression matrix using the *scanpy.pp.regress_out* function with the parameter setting. Batch effects between samples were preliminarily corrected by simple linear regression. We then used *scanpy.concatenate* function to merge the dataset processed by the unified process together at the data structure level with the parameter ‘*join = ‘inner*’’. Batch effects between datasets were preliminarily corrected by merging standardized datasets rather than directly merging count matrices.

For the merged matrices, we scaled each gene to unit variance using sc.pp.scale function with the parameter ‘max_value=10’, and then highly variable genes (HVGs) were selected using the *scanpy.pp.highly_variable_genes* function with default parameters. Next, principal component analysis (PCA) was performed on the matrix of HVGs to reduce noise and reveal the main axes of variation using the *scanpy.tl.pca* function, and the top 30 components were retained for downstream analyses. The batch effects were corrected by the BBKNN^48^ algorithm (v1.6.0), which detected the top nearest neighbors of each cell from each batch respectively instead of the entire cell pool. The parameters of the *scanpy.external.pp.bbknn* function were set to ‘‘*batch_key=’dataset’, n_pcs=30*’’. As comparison of retaining batch effects, the neighborhood graph was computed using *scanpy.pp.neighbors* function with default parameters following PCA. Finally, Uniform Manifold Approximation and Projection (UMAP) was employed for visualization via the *scanpy.tl.umap* function with default parameters.

### Preliminary annotation

We performed the unsupervised clustering to unveil the structure of IDH-GBM TME population using the *scanpy.tl.leiden* function with the parameter ‘*resolution=2.0*’ and the first round annotation was performed based on the expression of a predefined dictionary of canonical markers, including myeloid cells (*CD14, CD68, FCGR3A, CSF1R, C1QA*), lymphocytes (*CD3D, CD3E, CD8A, NKG7, CD4, CD19*), endothelial cells (*PECAM1, FLT1, ENG, VWF, CLDN5*), stromal cells (*COL1A1, COL1A2, FAP, PDGFRB, ACTA2*), malignant cells (*EGFR, PTPRZ1, METRN, SOX2, NRCAM*), astrocytes/neurons (*GFAP, SLC1A2, NRCAM, TUBB2B, DCX*), and oligodendrocytes (*MBP, PLP1, CLDN11, MOBP, CNP*). The cluster-specific marker genes were identified using the *scanpy.tl.rank_genes_groups* function with the parameter of ‘*method=t-test*’. ‘Express Analysis’ module of Metascape^49^ (https://metascape.org/gp/index.html) was used to confirming the identification of myeloid cells, lymphocytes, endothelial cells and stroma cells.

Despite oligodendrocytes segregating as an independent cluster, malignant cells remained transcriptionally indistinguishable from neurons and astrocytes when assessed exclusively through canonical marker gene expression. Thus, using non-immune, non-endothelial, and non-stromal cells as input with oligodendrocytes designated as reference normal cells via *infercnvpy.tl.infercnv function with the parameter “reference_cat = “Oligodendrocyte” in Infercnvpy*^50^ *(v0.5.0)*, malignant cells were distinguished from both neurons and astrocytes. The hallmark genomic alterations of GBM^20^ - gain of chromosome 7 and loss of chromosome 10 - were utilized as primary classification criteria. However, this approach still failed to resolve transcriptional distinctions between astrocytes and neurons at single-cell resolution.

### Further annotation

For Malignant cells, we performed unsupervised clustering of malignant cells using consensus non-negative matrix factorization (cNMF) implemented in Omicverse^51^ (v1.6.10). The malignant cell count matrix was first preprocessed using *omicverse.pp.preprocess* with the parameter “mode=’shiftlog|pearson’, n_HVGs=2000” to select highly variable genes, followed by scaling (*omicverse.pp.scale*) and principal component analysis (*omicverse.pp.pca*). We then initialized cNMF decomposition using *omicverse.single.cNMF* function with the parameter ‘*components=np.arange(2,20), n_iter=20, num_highvar_genes=2000*’, performing factorization using *omicverse.factorize* function before combining results using *omicverse.combine* function, both with default parameter. To find the optimal number of programs, the *k_selection_plot* function with default parameter was used to calculate the stability and reconstruction error. Finally, consensus clustering was performed using the *omicverse.consensus* function with the parameter ‘k=15, density_threshold=0.2’. Programs showing marked overexpression of myeloid, lymphoid, endothelial, stromal, or oligodendrocyte markers and containing relatively few cells were identified as contamination and subsequently excluded from further analysis.

The characterization of non-malignant cells proceeded through sequential classification phases, beginning with lineage assignment using canonical markers: myeloid populations included microglia-derived macrophages (MDMs; *TMEM119, P2RY12, CX3CR1*), bone marrow-derived macrophages (BMDMs; *CD14, FCGR3A/CD16, CD163, MRC1/CD206*), neutrophils (*S100A8, S100A9, LYZ*), dendritic cells (DCs, *CCR7, CLEC9A, FCER1A*), mast cells (*CPA3, TPSAB1*), and proliferating subsets (*MKI67, TOP2A, PCNA*); lymphoid populations comprised CD4+ T cells (*CD4, IL7R, CD40LG*) including regulatory T cells (*FOXP3, CTLA4, TIGIT, IL2RA, TNFRSF18*), CD8+ T cells (*CD8A, CD8B, GZMK*), B cells (*CD79A, MS4A1/CD20*), plasma cells (*JCHAIN, XBP1, SDC1/CD138*), and natural killer cells (NKs, *NKG7, NCAM1/CD56, FCGR3A/CD16, KLRC1, FGFBP2*); endothelial cells (ECs) were subclassified as arterial (*SEMA3G, EFNB2, GJA4*), venous (*ACKR1, NR2F2, EPHB4*), capillary (*RGCC, CA4, CD36*), and tip-like (*KDR/VEGFR2, ESM1, ANGPT2, CXCR4, PGF*) variants; while stromal components included smooth muscle cells (SMCs, *ACTA2, DES, CNN1, MYH11*), pericytes (PCs, *PDGFRB, CSPG4/NG2, RGS4, ABCC9*), and cancer-associated fibroblasts (CAFs, *FAP, MMP2, COL1A1, PDGFRA, POSTN*).

Subsequent functional refinement primarily relied on differentially expressed genes identified by the *FindAllMarkers* function with default parameter in Seurat^52^ (v4.3.0) (**Table S2, Table S3**), with supplementary evidence drawn from one or more of the following information: (1) pathway enrichment, (2) gene regulatory network inference, or (3) CellChat-mediated cell communication patterns. For instance, the myCAF.01 subpopulation was definitively annotated as ‘myCAF.01.COL1’ based on its characteristic collagen I signature (COL1A1/COL1A2 upregulation), extracellular matrix pathway activation, and demonstrated COL1A1-CD44 interactions with both E-MDSC and Mal.cNMF7 populations.

### Quantitative Analysis of Cell Cluster Distribution Preferences

Building upon the STARTRAC-dist framework developed by Zhang et al.^53^, we implemented a robust quantitative approach to assess cell cluster distribution patterns across treatment groups. Our analytical pipeline calculated the ratio of observed-to-expected cell frequencies (*R*_o/e_) to precisely quantify cluster-level enrichment or depletion according the following formula:

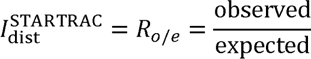

In which expected values were derived from chi-squared test contingency tables comparing cell clusters against treatment groups. This method provides distinct advantages over conventional statistical testing by: (1) generating continuous enrichment scores rather than binary significance thresholds. For example, if *R*_o/e_ > 1, it suggests that cells of the given cell cluster are more frequently observed than random expectations in the specific treatment group, that is, enriched. If *R*o/e < 1, it suggests that cells of the given cell cluster are less frequently observed than random expectations in the specific treatment group, that is, depleted; (2) maintaining directional information about cluster redistribution that is lost in traditional chi-squared values, which are defined as 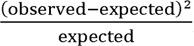; and (3) enabling cross-comparison of distribution preferences among all cell clusters within our annotated taxonomy.

### Cell Type Benchmarking with Published Signatures

Cell type identification and validation were performed through systematic benchmarking against established gene signatures from published studies (**Table S4**). For each cell subpopulation in our atlas, we first identified differentially expressed genes using the *FindMarkers* function in Seurat, followed by selection of the top 100 most significant genes based on statistical metrics. These gene signatures were then compared with published marker sets using Seurat’s *AddModuleScore* function to compute enrichment scores for specific cellular states.

For malignant cells, we benchmarked our classifications against four major established frameworks: (1) Ruiz-Moreno et al.’s^54^ astrocyte (AC)-like, neural precursor cell (NPC)-like, oligodendrocyte precursor cell (OPC)-like, and mesenchymal (MES)-like states; (2) Neftel et al.’s^43^ AC-like, OPC-like, and NPC-like meta-modules with NPC1 and NPC2 subdivisions; (3) Jackson et al.’s^25^ T1–T10 malignant cell classification system; and (4) Gavish et al.’s^55^ 41 consensus meta-programs representing transcriptional intratumoral heterogeneity patterns across 24 tumor types.

For myeloid populations, we specifically evaluated three functionally characterized subsets: (1) Jackson et al.’s^25^ early progenitor myeloid-derived suppressor cells (E-MDSCs) showing upregulation of metabolic and hypoxia pathways located in the pseudopalisading region; (2) Wang et al.’s^26^ hypoxia-responsive tumor-associated macrophages (Hypoxia-TAMs) localized in the peri-necrotic niches; and (3) Du et al.’s^27^ GPNMB+ MDMs localized in the peri-necrotic areas. (4) Miller et al.’s^56^ four consensus immunomodulatory programs, comprising the ‘microglial inflammatory’ and ‘systemic inflammatory’ programs, the ‘complement immunosuppressive’ program (associated with dexamethasone treatment), and the ‘scavenger immunosuppressive’ program (unique to primary CNS tumors and specifically enriched in hypoxic and vascular niches).

For endothelial cells, we integrated signatures from two high-resolution atlases by Xie et al.: (1) the 2024 atlas^57^ distinguishing GBM-EC type 1 (characterized by high vascular permeability and basement membrane remodeling genes like PLVAP and COL4A1) and GBM-EC type 2 (defined by Wnt signaling activation and partial blood-brain barrier retention); and (2) the 2021 atlas^58^ delineating distinct anatomical phenotypes, including the quiescent Peripheral EC type I (Pe1) enriched in BBB transporters and the angiogenic, tip-cell-like Tumor Core EC type I (Co1).

For stromal components, we synthesized three complementary classification schemes: (1) Poon et al.’s^15^ mesenchymal stromal cell system including astrocytic, fibroblastic, and fibropericytic subtypes; (2) Gao et al.’s^59^ cancer-associated fibroblast taxonomy comprising myofibroblastic (myCAF), inflammatory (iCAF), and antigen-presenting (apCAF) fibroblasts; and (3) Du et al.’s^27^ recent stromal atlas defining COL6A3+ tumor-associated fibroblasts (TAFs), a specialized matrix fibroblast subset characterized by robust extracellular matrix remodeling signatures and spatial localization within MVP niches.

### Pathway Enrichment analysis

Pathway enrichment was assessed through two complementary computational approaches to characterize biological processes across cell subpopulations. Pathway activity was inferred using the decoupleR package^60^ (v2.9.7) with the PROGENy model, which provides curated pathway-target interactions with regulatory weights for 14 major signaling pathways including Androgen, EGFR, Estrogen, Hypoxia, JAK-STAT, MAPK, NFkB, p53, PI3K, TGFb, TNFa, Trail, VEGF, and WNT. The analysis utilized the top 500 human pathway-responsive genes ranked by p-value, with normalized log-transformed count matrices processed using the Multivariate Linear Model (mlm) method via the *run_mlm()* function with the parameters “.source = ‘source’, .target = ‘target’, .mor = ‘weight’, minsize = 5”. Gene set enrichment analysis (GSEA) was performed on specific cell subpopulations. For each subpopulation, differentially expressed genes were first identified using the *FindMarkers* function in Seurat, followed by selection of the top 100 most significant genes based on statistical metrics. These gene sets were then analyzed using the *GSEA()* function from clusterProfiler^61^ (v4.14.6) package with two distinct collections from the Molecular Signatures Database (MSigDB): HALLMARK gene sets (c1.hallmark) and the legacy KEGG pathway collection (c2_kegg_legacy).

### Developmental Potential Quantification Using CytoTRACE 2

To assess cellular developmental potential across our single-cell atlas, we employed CytoTRACE 2^62^ (v1.1.0), an interpretable deep learning framework that predicts absolute developmental potential from scRNA-seq data. This validated algorithm classifies cells into discrete potency categories (totipotent, pluripotent, multipotent, oligopotent, unipotent, and differentiated states) while simultaneously generating continuous developmental potential scores ranging from 0 (terminally differentiated) to 1 (totipotent). We applied the *cytotrace2()* function with default parameters to leverage the algorithm’s deep neural network architecture trained on diverse developmental datasets to infer developmental potential based on gene expression patterns, where higher CytoTRACE scores (approaching 1) indicate greater developmental plasticity and lower scores reflect lineage commitment.

### Gene Regulatory Network (GRN) Inference and Transcription Factor Activity Analysis

GRN inference was performed using the pySCENIC pipeline^63^ (v0.12.1). To address data sparsity, we constructed metacells for each subpopulation. Specifically, we performed nearest-neighbor graph construction using the *FindNeighbors* function with parameter “k.param=10”, followed by high-resolution clustering using the *FindClusters* function with parameter “resolution=50” to identify metacell groupings. Metacell expression profiles were generated by summing raw counts across all cells within each cluster. Metacells containing fewer than 10 cells were filtered out. The aggregated metacell matrix was further filtered to retain genes expressed in at least 5 metacells and genes present in the hg38 cisTarget database. GRN inference was performed using GRNBoost2 (arboreto_with_multiprocessing.py) and a human TF list derived from motifs-v10nr_clust-nr.hgnc-m0.001-o0.0.tbl. Putative regulons were refined using cis-regulatory motif analysis (pyscenic ctx) against both promoter (hg38_500bp_up_100bp_down_full_tx_v10_clust.genes_vs_motifs.rankings.feather) and extended genomic regions (hg38_10kbp_up_10kbp_down_full_tx_v10_clust.genes_vs_motifs.rankings.feather) databases. Regulon activity scores were calculated for individual cells using the AUCell algorithm implemented via the *ComputeModuleScore* function in Seurat (v4.3.0), with results stored as a new assay for downstream analysis.

### Spatial Transcriptomic Data Collection

Spatial transcriptomic datasets were integrated from five previously published studies representing 48 patient samples profiled using the 10x Visium platform. The studies employed varying sample preservation methods and sequencing strategies: Ravi et al.^64^ (n = 16): Fresh frozen IDH-wildtype glioblastoma tissues embedded in OCT compound were sectioned at 10μm thickness. Libraries were prepared using the 10x Visium Spatial Gene Expression kit and sequenced on Illumina NextSeq 550. Mei et al.^13^ (n = 16): Fresh frozen GBM tissues were cryosectioned at 10μm thickness and processed using Visium Spatial Gene Expression kits. Libraries were sequenced on Illumina HiSeq X Ten. Ren et al.^65^ (n = 3): Fresh frozen GBM tissues were sectioned at 10μm thickness. Libraries were sequenced on both Illumina NovaSeq 6000 (short-read) and Oxford Nanopore PromethION/MinION platforms (long-read). Jackson et al.^25^ (n = 2): Formalin-fixed paraffin-embedded (FFPE) glioblastoma tissues were sectioned and processed using the Visium CytAssist Spatial Gene Expression protocol specifically designed for FFPE samples. Libraries were sequenced on Illumina NovaSeq 6000. Greenwald et al.^66^ (n = 13): Newly generated spatial transcriptomic data from fresh frozen GBM tissues. Libraries were prepared using standard Visium Spatial Gene Expression protocols and sequenced on Illumina platforms. All studies utilized the Visium platform but employed distinct sample preservation methods (fresh frozen vs. FFPE) and sequencing technologies, representing the technical diversity in spatial transcriptomic approaches for studying glioblastoma.

### Robust Cell Type Decomposition (RCTD) deconvolution

Cell type deconvolution was performed using RCTD as implemented in the spacexr^67^ package (v2.2.0). Comprehensively annotated GRIT-Atlas was employed as reference. For each spatial sample, we created an RCTD object using the *create.RCTD* function. Deconvolution was executed using *run.RCTD* with parameter “doublet_mode = ‘doublet’” to account for potential cell type mixtures within spatial spots. The resulting cell type proportions were incorporated into the spatial Seurat object metadata via the *AddMetaData* function. Cell type weight matrices were systematically extracted and synchronized with corresponding spatial spots, ensuring only common spots between the weight matrix and spatial object were retained for downstream analysis to maintain data integrity.

### MistyR Co-localization inference

Spatial inter-cellular relationships were analyzed using the MISTy framework in the mistyR^68^ package (v1.10.0) with a specific focus on identifying cell types that co-localize with the cNMF7 malignant subpopulation. Based on the RCTD deconvolution results, the MISTy framework defined two complementary views for analysis: (1) an *intraview* capturing cell type compositions within each spot, and (2) a *paraview* capturing cellular compositions in the surrounding tissue microenvironment with a radius determined by the mean distance to the nearest neighbor plus one standard deviation. For each of the cell subpopulations in the GRIT-Atlas, we selected the view (intra or para) that provided the strongest explanatory power for spatial relationships as determined by the highest importance values in the MISTy analysis.

The MISTy pipeline was executed independently for each of the 48 spatial samples using the *run_misty* function. For each sample, we extracted the importance values quantifying the spatial relationships between the cNMF7 malignant subpopulation (as predictor) and all other cell subtypes (as targets) from the aggregated results. Results across all samples were systematically compiled to identify consistent spatial relationship patterns between cNMF7 and other cell subtypes in the tumor microenvironment.

### Integration and Quantitative Analysis of Recurrent Patterns

To identify recurrent spatial relationship patterns between the cNMF7 malignant subpopulation and other cell types across all 48 samples, we implemented a quantitative thresholding approach. For each cell type, we calculated the frequency of significant spatial associations with cNMF7 across all 48 samples at multiple percentile thresholds (ranging from 10% to 50%). Importance values from MISTy analysis were binarized such that the top n% of values (where n = threshold × 100) were set to 1 and all others to 0, enabling computation of recurrence frequencies for each cell type across the cohort.

The resulting frequency matrices were analyzed using hierarchical clustering with Euclidean distance and complete linkage to identify modules of cell types exhibiting consistent spatial relationship patterns with cNMF7. Since MISTy’s importance values indicate the statistical significance of spatial relationships without distinguishing between co-localization (positive association) and mutual exclusion (negative association), we determined the directionality of these relationships through quantitative spatial gradient analysis relative to specific histopathological niches (MVP and PAN), as described below.

### Spatial Association Analysis with MVP and PAN

To quantitatively analyze spatial relationships between cellular compositions and key histopathological structures in glioblastoma, we employed SPATA2 (v3.1.0), a computational framework specifically designed for spatial transcriptomics data analysis that enables investigation of expression patterns within tissue architecture without prior data grouping. This approach implements the Spatial Gradient Screening algorithm (SAS) as described in Kueckelhaus et al.^69^, which facilitates supervised detection of histology-associated expression patterns.

We focused on two hallmark glioblastoma features: MVP and PAN^18^. Senior neuropathologists identified tissue sections enriched for these features (#UKF259, #UKF260, #UKF275 for MVP; #UKF243, #UKF313, GBM03422 for PAN). Using the *downloadSpataObject* function from the SPATAData package, we obtained pre-processed SPATA2 objects for two representative sections (#UKF275 for MVP, #UKF313 for PAN) following Kueckelhaus et al.’s methodology. Neuropathologists manually annotated the spatial extent of these regions on H&E-stained slides using *createImageAnnotations* function, identifying three distinct pseudopalisading regions surrounding necrotic areas in sample #UKF313 and six MVP regions in sample #UKF275. We then applied the Spatial Gradient Screening algorithm to quantify spatial relationships between cellular proportions (from RCTD deconvolution) and distance to these pathological structures. Distance to annotated regions was calculated using *getCoordsDfSA* function with parameters unit = ‘mm’ and binwidth = ‘200um’. Distance metrics were incorporated as meta-features via *addFeatures* function. Expression gradients of cell type proportions relative to pathological structures were quantified using SAS-specific functions including *plotSasLineplot* function (for linear trends) and *plotSasHeatmap* function (for spatial patterns across distance bins). These spatial gradients allowed us to confirm actual co-localization (increasing proportion with proximity to MVP/PAN) versus mutual exclusion (decreasing proportion with proximity) for the cell modules identified in the recurrent pattern analysis.

### Cell-Cell Communication Analysis

Cell-cell communication analysis was performed using CellChat^70^ (v1.6.1), a computational tool that infers intercellular communication networks by integrating single-cell transcriptomic data with a curated database of ligand-receptor interactions. We utilized CellChatDB.human, which contains manually curated literature-supported ligand-receptor interactions, including secreted signaling, ECM-receptor, and cell-cell contact interactions. To investigate specific communication patterns between malignant cells (cNMF7 subtype), cancer-associated fibroblasts (myCAF.02.COL6 and myCAF.03.COL1), and myeloid-derived suppressor cells (E_MDSCs.ADM.HIF), we integrated corresponding subsets from our single-cell atlas and the CellChat pipeline was executed as follows: (1) normalized expression data were input to *createCellChat()* function; (2) over-expressed genes and interactions were identified using *identifyOverExpressedGenes()* function and *identifyOverExpressedInteractions()* function; (3) communication probabilities were computed using *computeCommunProb()* function with the ‘triMean’ method, followed by pathway-level inference with *computeCommunProbPathway()* function; (4) aggregated networks were calculated using *aggregateNet()* function, and network centrality analysis was performed using *netAnalysis_computeCentrality()* function.

### Meta-Analysis for prognosis and response to ICB

Meta-analysis was conducted with the “meta” package^71^. To reduce inter-cohort heterogeneity, gene expression values were log_₂_-transformed and standardized into z-scores across patients. Response outcomes were evaluated using risk ratios (RR) by the *metagen* function with the parameter “sm = ‘RR’”, while survival associations were assessed using Benjamini–Hochberg adjusted hazard ratios (HR) by the *metabin* function with the parameter “sm = ‘HR’”, of which the input was derived from univariate Cox regression. Both measures are reported with 95% confidence intervals (CI) based on a random-effects model, and effects were considered statistically significant at p < 0.05. Heterogeneity across studies was quantified with Chi-squared tests and I² statistics; a p-value below 0.05 accompanied by an I² value exceeding 50% was considered indicative of substantial heterogeneity.

### CosMx Spatial Transcriptomic Profiling and Cell Segmentation

Spatial transcriptomic profiling was performed on 5μm FFPE sections from 15 samples (obtained from 11 patients, **Table S6**) using the CosMx Spatial Molecular Imager (SMI) (NanoString Technologies) equipped with the 6000-plex Human Universal Cell Characterization Panel. The experimental workflow followed a rigorous multi-day protocol. Briefly, FFPE sections were dried at 65°C, deparaffinized in xylene, and rehydrated through a graded ethanol series. Heat-induced antigen retrieval was executed at 100°C for 15 min using a pressure cooker (BioSB), followed by stabilization in DEPC-treated water. Tissues were then permeabilized with a specialized digestion buffer at 40°C for 30 min. To facilitate precise image registration, fiducial markers (0.001%) were applied and incubated for 5 min. Following post-fixation in 10% neutral buffered formalin (NBF), sections were treated with Sulfo-NHS-acetate for 15 min to reduce background autofluorescence. For in situ hybridization, the CosMx RNA probe mixture was denatured at 95°C for 2 min and hybridized to the sections at 37°C overnight.

On the second day, stringency washes were performed twice for 25 min each using a pre-warmed 50% formamide/4×SSC solution at 37°C. To ensure high-fidelity cell boundary detection within the complex brain microenvironment, sections were incubated for 1 hour with the CosMx Human Neuroscience Cell Segmentation Kit. This specialized cocktail enabled a four-channel morphology-based segmentation strategy: DAPI for nuclei (Ch1), Hs Neuro rRNA for neuronal cytoplasm (Ch2), Mm/Hs Neuro Histone for nuclear proteins (Ch3), and Mm/Hs GFAP for glial cytoplasm (Ch4). Finally, the slides were assembled into flow cells, and cyclic readout was initiated on the CosMx instrument. Raw data were processed via the AtoMx Spatial Informatics Platform for transcript decoding and cell-level assignment, yielding a single-cell spatial expression matrix. For quality control, cells with fewer than 100 detected transcripts (nFeature_RNA < 100) were excluded from downstream analysis.

### Niche Analysis of MES-like Cells via CosMx SMI

To define the localized cellular architecture surrounding Mesenchymal-like (MES-like) cells, we conducted spatial niche analysis mainly using Seurat package (v5.1.0), FNN package^72^ (v1.1.4.1) and RcppML^73^ package (v0.3.7). For each individual MES-like cell, its spatial neighborhood was captured by identifying its 100-nearest neighbors (k = 100) based on 2D spatial coordinates via the *get.knnx* function in FNN package. A localized composition matrix was constructed by calculating the proportional frequency of 11 major cell types within this k=100 window. To decompose these complex localized environments into recurrent spatial patterns, NMF was performed on the normalized niche matrix using *nmf* function with the parameters “k = 5” in RcppML package. The resulting NMF embeddings (W-matrix) were integrated into the Seurat object as a dimensional reduction and utilized for unsupervised sub-clustering of MES-like cells within each sample using *FindNeighbors* function with parameter “reduction = ‘nmf’, dims = 1:5” and *FindClusters* function with parameter “resolution = 0.1, algorithm = 1” in Seurat package.

To enable a robust comparison across the nine samples, we integrated the niche sub-clusters while implementing a systematic background correction strategy. Recognizing that local cell-type enrichment can be confounded by tissue-wide abundance, we calculated the global cell-type fractions for each whole-tissue section. The average niche composition of each sub-cluster was normalized by its corresponding tissue background to generate a corrected enrichment score. These scores were then re-normalized to a sum of 1 to represent the relative enrichment of cell types within each niche. To ensure balanced representation across the cohort, a stratified downsampling strategy was applied, retaining a representative number of niche sub-clusters per sample to prevent specimens with high cell density from dominating the global landscape (**Table S7**).

Conserved spatial functional units, termed “Global MES Niches,” were identified by performing hierarchical clustering on the integrated and balanced niche correlation matrix. The optimal number of clusters (K) was objectively determined using the *silhouette* function in cluster package (v2.1.8.1), which maximized the average silhouette width across a range of K from 2 to 8. Global clustering was executed via the *hclust* function with parameter “method = ‘ward.D2’” parameter based on Pearson correlation coefficients (PCC) in stats package (v4.4.2). This hierarchical workflow partitioned the MES-like neighborhoods into four distinct global niches (S1–S3). Differential expression analysis was subsequently conducted using the *FindAllMarkers* function with default parameters to identify niche-specific molecular signatures.

### Multiplex Immunohistochemistry (mIHC)

mIHC was performed using a sequential staining strategy to characterize the GBM microenvironment. Briefly, 4-μm-thick FFPE sections were deparaffinized in Xylene, rehydrated through a graded series of ethanol (100%, 90%, and 70%), and rinsed in distilled water. Antigen retrieval was conducted via microwave heating in a dedicated retrieval solution at high power until boiling, followed by 15 min at low power. Endogenous peroxidase activity was quenched with 3% H_2_O for 10 min, and non-specific binding was blocked with normal goat serum for 30 min at room temperature. The sections were then subjected to multiple rounds of sequential staining. In each round, slides were incubated with primary antibodies for 1 h, followed by secondary antibody incubation (10 min) and signal amplification using tyramide signal amplification (TSA)-based fluorescent dyes (10 min). To enable multi-target labeling, the previous primary-secondary antibody complexes were stripped between each round using microwave-based heat treatment. To systematically characterize the spatial architecture, cellular identities, and functional interactions within distinct GBM microenvironmental niches, three tailored mIHC panels were deployed sequentially according to a hierarchical experimental strategy: Panel 1 (Screening Panel, 2 markers/3 colors) was employed as an initial screening gate to identify and select FFPE tissue blocks containing rich PAN or MVP niches, given the marked regional heterogeneity of these pathological structures across clinical specimens. This screening panel target-labeled cellular hypoxia and vascular structures using anti-HIF-1α (cat. no. ab51608, Abcam) and anti-CD31 (ab182981, Abcam). Only clinical specimens validated to possess these definitive architectural niches were approved for further high-order multiplexing. Panel 2 (Phenotypic/Identity Panel, 6 markers/7 colors) was subsequently utilized to define the precise spatial co-localization, structural coordinates, and microenvironmental composition of the “spatial resistance triad” within the pre-screened PAN and MVP niches. This lineage-tracing panel consisted of anti-SOX2 (cat. no. ab92494, Abcam) for malignant cell states, anti-FAP (cat. no. ab53066, Abcam) for myCAFs, anti-CD33 (cat. no. ab269456, Abcam) and anti-HLA-DR (cat. no. ab92511, Abcam) for E-MDSCs, and anti-CD31 (cat. no. ab182981, Abcam) for endothelial structures. Panel 3 (Functional Panel, 6 markers/7 colors) was engineered to elucidate the mechanistical crosstalk within this immunosuppressive sanctuary, focusing specifically on the spatial engagement of the COL6A1–CD44 signaling axis within the triad infrastructure. This functional panel retained the niche-defining backbone of anti-SOX2, anti-FAP, anti-CD33, and anti-CD31, while incorporating anti-COL6A1 (cat. no. ab182744, Abcam) and anti-CD44 (cat. no. 15675-1-AP, Proteintech) to capture ligand-receptor interactions *in situ*. Following the final sequential staining round of each respective panel, nuclei were counterstained with DAPI for 10 min. Slides were subsequently mounted with fluorescence mounting medium and scanned using a high-resolution slide scanner to generate multiplexed multichannel images.

### In Vitro

#### Primary Tissue Dissociation and Patient-Matched Cell Isolation Preparation

Fresh tumor tissues were obtained from six patients diagnosed with IDH-WT GBM at the First Affiliated Hospital of Soochow University. All experimental procedures involving clinical human specimens were strictly reviewed and approved by the Ethics Committee of Soochow University. To ensure precise patient-matched cellular modeling, a single-cell suspension was prepared from each patient’s specimen to simultaneously isolate paired GSCs and CAFs. Briefly, fresh tumor tissues were finely minced with sterile scalpels and enzymatically digested in papain (cat. no. M3826, AbMole Bioscience) at 37 °C for 30 min with intermittent vortexing. The resulting cell mixture was passed through a 50-μm cell strainer, rinsed with culture medium, and centrifuged at 500 × *g* for 5 min. To eliminate erythrocytes, the cell pellet was incubated with Red Blood Cell Lysis Buffer (cat. no. C3702, Beyotime Biotechnology) for 2 min, which was subsequently neutralized by adding Phosphate-Buffered Saline (PBS). Following centrifugation at 500 × *g* for 5 min, the fully dissociated single cells were resuspended in fresh PBS, counted, and immediately allocated into GSC or CAF purification pipelines.

#### Primary GSC Enrichment and Culture

For GSC enrichment, the designated single-cell suspension from each of the six patients was subjected to magnetic-activated cell sorting (MACS) using CD133 MicroBeads (cat. no. 130-097-049, Miltenyi Biotec) according to the manufacturer’s instructions. The enriched CD133+ GSCs (designated as GSC01 to GSC06) were subsequently maintained in serum-free GSC culture medium, which consisted of DMEM/F12 (cat. no. 12634010, Thermo Fisher Scientific) supplemented with B27 supplement (cat. no. 17504044, Thermo Fisher Scientific), 20 ng/mL recombinant human epidermal growth factor (EGF; cat. no. 10605-HNAE, Sino Biological), and 20 ng/mL recombinant human basic fibroblast growth factor (bFGF; cat. no. 10014-HNAE, Sino Biological). Successful isolation and establishment of primary GSCs were validated by the formation of stable, non-adherent oncospheres under inverted phase-contrast microscopy. All GSC cultures were maintained in a humidified incubator at 37 °C with 5% CO2.

#### Primary CAF Purification and Characterization

Concurrently, the parallel portion of the single-cell suspension from each patient was utilized to establish patient-matched CAFs (designated as CAF01 to CAF06, fully paired with GSC01 to GSC06) via a modified serial trypsinization approach^32^. Dissociated cells were seeded and cultured in complete Fibroblast Medium (FM; cat. no. 2301, ScienCell Research Laboratories), composed of 500 mL of fibroblast base medium, 10 mL of fetal bovine serum (FBS; cat. no. 0010, ScienCell Research Laboratories), 5 mL of fibroblast growth supplement (FGS; cat. no. 2352, ScienCell Research Laboratories), and 5 mL of penicillin/streptomycin solution (P/S; cat. no. 0503, ScienCell Research Laboratories). During passages, cells were treated with Trypsin-EDTA (cat. no. 25200072, Thermo Fisher Scientific), and the detachment process was dynamically monitored under an inverted microscope. Once approximately 50% of the cells (predominantly the less-adherent primary GBM cells) had detached from the substratum, the reaction was immediately halted by adding PBS. The supernatant containing these weakly adherent cells was discarded. Subsequently, the remaining strongly adherent CAFs were detached by an extended trypsinization for 10 min and transferred to a fresh culture plate.

Homogeneous cell populations with distinct fibroblast morphology were successfully established within five weeks (approximately 5 passages). At passage 5 (P5), the identity and purity of the isolated CAFs were rigorously validated via a dual-dimensional strategy. For transcriptomic validation, total RNA was extracted using TRIzol Reagent (cat. no. R0016, Beyotime Biotechnology) for subsequent RNA sequencing (RNA-seq). For protein-level validation, Western blot analysis was performed to examine the expression of classic fibroblast markers, including FAP (cat. no. AF3715-SP, R&D Systems) and α-SMA (cat. no. MAB1420-SP, R&D Systems). Only successfully validated CAFs under 10 passages (P5 to P10) were utilized for subsequent functional experiments. All cell cultures were maintained in a humidified incubator at 37 °C with 5% CO2.

#### Screening of Pro-Stemness Microenvironmental Ligands and CD44 Receptor Blockade Assay

To identify the key microenvironmental components regulating glioblastoma stemness, a two-tier screening strategy was implemented using patient-derived GSCs. Initially, the baseline expression of CD44 was evaluated across the six established primary GSC lines via WB using antibodies against CD44 (cat. no. 15675-1-AP, Proteintech). The two GSC lines exhibiting the highest CD44 abundance were selected for subsequent functional assays. In the first-tier screening (ligand-induced stemness evaluation), GSCs in the logarithmic growth phase were treated with six candidate recombinant human matrix proteins: COL1A1 (cat. no. AP97587, SAB), COL1A2 (cat. no. AP90701, SAB), COL6A1 (cat. no. AP90909, SAB), COL6A2 (cat. no. AP90910, SAB), COL6A3 (cat. no. AP90911, SAB), and FN1 (cat. no. AP90493, SAB). Each recombinant protein was administered at a final concentration of 5 μg/mL. After 24 h of incubation, oncosphere formation was dynamically examined under an inverted microscope. Concurrently, total protein was extracted to determine the expression of core stemness markers via WB, utilizing antibodies against SOX2 (cat. no. GB15336, Servicebio; 1:500) and Nestin (cat. no. GB155683, Servicebio; 1:500). In the second-tier screening (receptor-dependence validation), a neutralization assay was conducted to ascertain whether the ligand-driven stemness enhancement was specifically mediated by the CD44 receptor. The selected GSC lines were pre-treated with an anti-CD44 neutralizing antibody (cat. no. BE0039, BioXCell) at a concentration of 10 μg/mL for 24 h to block the cell-surface CD44 binding sites. Subsequently, the cells were challenged with the aforementioned six recombinant proteins (5 μ/mL) for an additional 24 h. The impacts of this receptor blockade on GSC sphere-forming capacity and stemness marker expression were sequentially evaluated using the identical methodology described above.

#### Virtual Screening and Experimental Validation of CNS-Penetrant Anti-Fibrotic Compounds

To discover potent therapeutic agents capable of disrupting the tumor-shielding fibrotic niche in glioblastoma, a hybrid *in silico* and *in vitro* phenotypic screening workflow was established. Candidate compounds were derived from the intersections of three commercially available small-molecule libraries (TargetMol): (1) the Drug Repurposing Compound Library (cat. no. L9200), encompassing 5,745 approved or clinical-stage drugs; (2) the Anti-Fibrosis Compound Library (cat. no. L9810), containing 1,800 molecules targeting fibrotic pathways; and (3) the CNS-Penetrant Compound Library (cat. no. L5900), consisting of 750 validated BBB-permeable compounds. To further refine the clinical translatability of these candidates, the ADMETlab 3.0 computational platform was utilized to comprehensively evaluate their physicochemical parameters and pharmaceutical chemical friendliness^74^. Specifically, the Distribution/BBB Penetration module was implemented to predict the probability of molecules crossing the BBB. Based on the empirical decision threshold of the platform, only compounds exhibiting excellent BBB permeability scores within the predictive probability range of 0 to 0.3 were retained. Through the intersection of the aforementioned three physical compound libraries and the *in silico* ADMET filtering, a total of 24 candidate compounds were identified and purchased from TargetMol Chemicals (**Table S9**).

For *in vitro* experimental validation, the two primary, patient-matched CAF lines were seeded and cultured until optimal density. The CAFs were subsequently treated with each of the 24 candidate compounds at a final concentration of 20 μM for 48 h. Following incubation, the cell culture supernatants were harvested, clarified via centrifugation at 1,000 × *g* for 20 min to remove cellular debris, and immediately processed to quantify the extracellular secretion of COL6A1. The quantitative measurement of human COL6A1 in the biological fluids was performed using a commercial sandwich Enzyme-Linked Immunosorbent Assay (ELISA) kit (cat. no. EK13362, SAB) according to the manufacturer’s protocol. Optical densities were measured at 450 nm using a calibrated microplate reader to identify lead anti-fibrotic drugs capable of robustly downregulating CAF-mediated COL6A1 secretion.

#### High-Throughput 4D-DIA Quantitative Proteomics of Lead-Treated CAFs

Following the selection of Lacidipine, primary human CAFs were treated with Lacidipine (20 uM) or vehicle for 48 hours in triplicate to map global proteomic changes. Cells were harvested and lysed with SDS lysis buffer containing 1 mM PMSF, protease, and phosphatase inhibitors. Protein concentration was measured via a BCA kit (cat. no. 23225, Thermo Scientific) and validated by 4%-12% SDS-PAGE. Equal protein amounts were captured on hydrophilic and hydrophobic Sera-Mag SpeedBeads carboxyl magnetic beads for SP3 processing, reduced with DTT, and digested with Trypsin and LysC overnight. Peptides were desalted, dried, and mixed with iRT standard peptides (cat. no. Ki-3002-2, Biognosys) at a 1:20 ratio. High-resolution LC-MS/MS was executed on a Vanquish Neo LC system coupled to a timsTOF HT mass spectrometer (Bruker). Peptides were separated over a 10-minute linear gradient (7.5% to 35% buffer B) on a C18 column at a flow rate of 0.8 uL/min, using mobile phases prepared with mass spectrometry-grade water and acetonitrile. Data were acquired in dia-PASEF mode across a 300-1500 m/z mass range and 0.7-1.3 ion mobility window. Raw files were quantified using DIA-NN software against the UniProt Homo sapiens database with peptide and protein FDR thresholds under 0.01. Differentially expressed proteins were filtered via an unpaired two-sided t-test using thresholds of fold-change greater than or equal to 2 or less than or equal to 0.5, and a p-value less than 0.05.

### In Vivo

#### Orthotopic Murine Glioblastoma Model and In Vivo Bioluminescence Imaging

All animal experimental protocols were strictly conducted in accordance with the guidelines for animal research institutionalized by Soochow University and were officially approved by the Ethics Committee of Soochow University. The murine glioma cell line GL261 was acquired from the American Type Culture Collection and cultured in DMEM supplemented with 10% fetal bovine serum (FBS; cat. no. A5669701, Thermo Fisher Scientific). Five-week-old male C57BL/6J mice were purchased from the SLAC Laboratory Animal Center (Shanghai, China) and allowed to acclimatize before procedures. Prior to tumor inoculation, the mice were randomly allocated into distinct experimental groups. To facilitate non-invasive longitudinal monitoring, GL261 cells were stably labeled with firefly luciferase (GL261-Luc). Intracranial tumors were established via stereotactic injection, wherein a total of 1 x 10^5 GL261-Luc cells suspended in serum-free medium were stereotactically inoculated into the right striatum of each anesthetized mouse. To track intracranial tumor progression *in vivo*, bioluminescence imaging (BLI) was regularly performed on days 7, 14, 21, 28, 35, and 42 post-injection using an imaging system following the intraperitoneal administration of D-luciferin. Upon termination of the study or when mice met the predefined humane endpoints, animals were euthanized, and Kaplan–Meier survival curves were constructed to statistically evaluate the overall survival duration across groups.

#### Generation of Stable Col6a1-Knockdown GL261 Glioma Cells via Lentiviral Transduction

To systemically evaluate the role of tumor-derived collagen VI in shaping the TME, a stable Col6a1-knockdown GL261 cell line was established using a lentiviral vector system. The recombinant lentiviral particles expressing short hairpin RNA (shRNA) targeting mouse Col6a1 (sh-Col6a1) and a non-targeting negative control lentivirus (LV3-NC) were commercially engineered and synthesized by GenePharma (Shanghai, China). The specific genomic core target sequence utilized for sh-Col6a1 was 5’-CCAAGCGCTTCATCGACAA-3’. The functional viral constructs were packaged with the vesicular stomatitis virus G (VSV-G) envelope protein, yielding a stock viral titer of 3 x 10^8 TU/mL.

For lentiviral transduction, GL261 cells were seeded in 6-well plates at a density of 2 x 10^5 cells per well in DMEM supplemented with 10% FBS and allowed to adhere overnight to reach approximately 50% confluence. Cells were subsequently transduced with either sh-Col6a1 or LV3-NC lentivirus at an optimized multiplicity of infection (MOI) of 50 in fresh complete medium. To enhance endocytic viral uptake, Polybrene (GenePharma) was co-administered into each well at a final working concentration of 5 ug/mL (10 ug per well). At 24 h post-transduction, the virus-containing supernatants were thoroughly replaced with fresh complete culture medium. At 72 h post-transduction, green fluorescent protein (GFP) expression was verified under an inverted fluorescence microscope to confirm successful viral integration. To eliminate non-transduced parental cells and isolate a stable population, cells were subjected to selection pressure in complete medium supplemented with 10 ug/mL of puromycin hydrochloride (GenePharma). The puromycin selection was maintained continuously for at least 4 days with fresh medium replenished every 48 h until 100% cell death was achieved in the non-transduced control wells. To rigorously confirm the knockdown efficiency of the target protein prior to in vivo experiments, total protein was extracted from the surviving resistant cells and subjected to Western blot analysis utilizing a anti-Col6a1 antibody (cat. no. ab182744, Abcam). The surviving cells, representing a validated stable mixed-clone population with robust Col6a1 silencing, were expanded and utilized within three passages for downstream in vivo orthotopic experiments.

#### Orthotopic Inoculation and In Vivo Bioluminescence Imaging

C57BL/6J mice were orthotopically inoculated with either the control or shRNA-mediated Col6a1-knockdown GL261 murine glioma cells to allow subsequent evaluation of microenvironmental perturbation. Successful intracranial tumor engraftment and real-time longitudinal growth kinetics were confirmed via bioluminescence imaging on day 7 post-injection. Following successful confirmation of the intracranial niches, mice were maintained under standard housing conditions for an additional week to facilitate matrix remodeling and stromal architecture adaptation. On day 14 post-inoculation, fresh tumor-bearing brain tissues were harvested, fixed in 4% neutral-buffered formalin for 24 h, and embedded in paraffin.

#### Immunohistochemical (IHC) Analysis of Microenvironmental Col6a1 Expression

FFPE brain sections (4-um thick) were subjected to IHC staining to delineate the precise cellular source of intra-tumoral collagen and map the alterations in microenvironmental Col6a1 deposition following tumor-specific silencing. Briefly, sections were deparaffinized in xylene and sequentially rehydrated through a graded series of ethanol. Antigen retrieval was performed in a microwave-heated citrate buffer (pH 6.0) at high power until boiling, followed by sustained maintenance at low heat for 15 min. To eliminate endogenous peroxidase activity, sections were treated with 3% hydrogen peroxide for 10 min at room temperature in the dark. After blocking non-specific binding with normal goat serum at room temperature for 30 min, the slides were incubated with a primary rabbit anti-Col6a1 antibody (cat. no. ab182744, Abcam) for 1 h in a humidified chamber. Subsequently, the sections were washed and incubated with a horseradish peroxidase (HRP)-conjugated secondary antibody for 10 min, followed by colorimetric visualization using a 3,3’-diaminobenzidine (DAB) substrate kit. The sections were then counterstained with hematoxylin, differentiated with 1% acid ethanol, and blued under running water. Finally, the slides were dehydrated, cleared in xylene, and mounted with neutral resin. All stained sections were digitized using a high-resolution slide scanner, and the histopathological scoring or positive area percentage of TME-associated Col6a1 was quantified to assess the relative contribution of glioma cells versus host-derived stromal components to the fibrotic niche.

#### In Vivo Therapeutic Regimen and Drug Administration

Upon bioluminescence imaging confirmation of tumor engraftment, C57BL/6J mice were stratified and assigned into four balanced experimental groups based on their initial tumor burdens: (1) Control IgG group, (2) Lacidipine monotherapy group, (3) TMZ + Pexidartinib combotherapy group, and (4) Triple-combination group (TMZ + Pexidartinib + Lacidipine). All small-molecule inhibitors were purchased from TargetMol Chemicals, including Temozolomide (TMZ; cat. no. T1178), Pexidartinib (PLX3397, anti-CSF1R; cat. no. T2115), and Lacidipine (cat. no. T1439). Treatment was initiated immediately after grouping on day 7. The specific pharmacological dosages, administration routes, and treatment frequencies were customized as follows: Lacidipine was administered at a dose of 10 mg/kg via daily oral gavage, and Pexidartinib was administered at a dose of 40 mg/kg via daily oral gavage. For cytotoxic chemotherapy, TMZ was administered at a dose of 25 mg/kg via intraperitoneal (i.p.) injection. To mitigate systemic hematological toxicity and better recapitulate clinical scenarios, the TMZ dosing schedule was administered as a cyclic regimen consisting of 5 consecutive days of dosing followed by a 2-day drug holiday (5 days on, 2 days off) each week. Lacidipine and Pexidartinib treatments were maintained on a continuous daily schedule throughout the study until the defined experimental or humane endpoints for survival analysis.

#### In Vivo Flow Cytometry Analysis of the Tumor and Immune Microenvironment

To systematically interrogate the cellular composition and functional states within the tumor microenvironment across the four therapeutic arms, multi-color full-spectrum flow cytometry was performed on fresh intracranial tumor tissues harvested from three representative mice per group.

##### Tumor Dissociation and Single-Cell Preparation

Freshly isolated mouse glioblastoma tissues were finely minced with sterile scalpels and digested in RPMI-1640 medium supplemented with 50 uL of FREEDOM universal tissue dissociation enzyme on a horizontal shaker at 37 degrees Celsius for 50 min. The resulting cell suspension was mechanically disrupted by gentle pipetting and passed through a 70-um cell strainer to remove tissue debris. Following centrifugation at 500 x g for 5 min, the single cells were washed with PBS and quantified. To achieve non-interfering phenotypic profiling and functional evaluation, the single-cell suspension from each sample was allocated into two parallel aliquots for baseline surface characterization and ex vivo stimulation assays, respectively.

##### Baseline Cell-Surface Staining

For the direct identification of immune lineages, tumor cells, fibroblasts, and endothelial compartments, the first aliquot of single cells (6 x 10^6 cells in a 50 uL staining volume) was initial-incubated with the Zombie Aqua Fixable Viability Kit (cat. no. 423102, BioLegend; 1:1,000) on ice for 10 min in the dark to exclude dead cells. Non-specific Fc receptor binding was subsequently blocked using a TruStain FcX anti-mouse Cd16/32 antibody (cat. no. 101320, BioLegend; 2 uL per 50 uL system) for 15 min at room temperature in the dark. Cells were then labeled with an optimized cell-surface fluorochrome-conjugated antibody cocktail at 4 degrees Celsius for 30 min in the dark. The precise capturing volumes per 50 uL reaction system were utilized as follows: Pacific Blue anti-Cd45 (cat. no. 157212, BioLegend; 0.25 uL), FITC anti-Cd3epsilon (cat. no. 103126, BioLegend; 1 uL), APC/Cyanine7 anti-Cd8a (cat. no. 100204, BioLegend; 0.125 uL), Brilliant Violet 421 anti-Cd279/Pd-1 (cat. no. 47-0081-82, Invitrogen; 1 uL), Brilliant Violet 605 anti-Cd366/Tim-3 (cat. no. 135231, BioLegend; 1 uL), PerCP/Cyanine5.5 anti-Cd11b (cat. no. 119721, BioLegend; 0.125 uL), Brilliant Violet 650 anti-Ly-6C (cat. no. 101228, BioLegend; 1 uL), Brilliant Violet 711 anti-Ly-6G (cat. no. 128049, BioLegend; 0.5 uL), APC anti-Cd31 (cat. no. 127643, BioLegend; 1 uL), CoralLite Plus 647 anti-Fap (cat. no. FAB3715R, R&D Systems; 1.25 uL), and PE anti-Cd44 (cat. no. 55413, BD Biosciences; 0.5 uL).

##### Ex Vivo Stimulation and Intracellular Cytokine Staining

Concurrently, to evaluate the cytotoxic capacity of tumor-infiltrating lymphocytes without compromising baseline membrane marker readouts, the parallel cell aliquot was subjected to ex vivo activation. Cells were cultured in RPMI-1640 medium containing 10% FBS and supplemented with a 1x working concentration of eBioscience Cell Stimulation Cocktail containing protein transport inhibitors (cat. no. 00-4970-03, Invitrogen) at 37 degrees Celsius with 5% CO2 for 4 h. Following stimulation, cells were harvested, washed, and processed through dead cell exclusion and Fc blockade matching the protocols described above. To evaluate intracellular functional molecules, cells were subjected to fixed-intracellular compartmentalization using the eBioscience Foxp3/Transcription Factor Staining Buffer Set (cat. no. 00-5523-00, Invitrogen). Cells were incubated in 200 uL of Fixation/Permeabilization working solution for 1 h at room temperature in the dark. After washing with 1x Permeabilization Buffer, the cells were resuspended in approximately 50 uL of 1x Permeabilization Buffer and stained with intracellular functional antibodies for 60 min at room temperature in the dark. The precise volumes per 50 uL reaction system were utilized as follows: PE/Cyanine7 anti-Granzyme B (cat. no. 12-0441-81, Invitrogen; 1 uL) and PE-Dazzle 594 anti-Ifn-gamma (cat. no. 372214, BioLegend; 1 uL). Finally, cells were washed twice and resuspended in 200 uL of staining buffer for spectral acquisition.

##### Data Acquisition and Immunophenotypic Gating

Multicolor data acquisition was executed on a 3-laser Cytek Aurora full-spectrum flow cytometer (Cytek Biosciences), and multi-parametric analysis was conducted via FlowJo software (version 10.10, FlowJo LLC). After sequentially excluding doublets and dead cells (Zombie Aqua-), specific intra-tumoral cellular lineages were precisely gated as follows: CTLs were defined as Cd45+ Cd3+ Cd8a+ Granzyme B+ Ifn-γ +; Tex were defined as Cd45+ Cd3+ Cd8a+ Pd-1+ Tim-3+; M-MDSCs were identified as Cd45+ Cd3- Cd11b+ Ly-6G- Ly-6Chigh; MES-like glioma cells were characterized as Cd45- Fap- Cd44+; CAFs were distinguished as Cd45- Fap+; and the tumor endothelial vasculature was mapped as Cd45- Cd31+.

### Single-Cell RNA Sequencing and Microenvironmental Transcriptomic Profiling

To further dissect the non-immune microenvironmental remodeling induced by Lacidipine, single-cell RNA sequencing was performed on tumors from the TMZ + anti-CSF1R doublet group and the Triple-combination group on day 7 post-treatment. Fresh intracranial tumor tissues from two representative mice per group were pooled, enzymatically dissociated into single-cell suspensions, and depleted of immune cells via magnetic-activated cell sorting (MACS) using CD45 MicroBeads (cat. no. 130-052-301, Miltenyi Biotec). The isolated viable CD45-negative single-cell fraction was immediately loaded onto the MobiNova-100 microfluidic platform using the MobiCube High-throughput Single Cell 3 Prime Transcriptome Kit V2.1 (MobiDrop Co., Ltd.) for droplet encapsulation, barcoding, and library construction. Following sequencing on an Illumina platform, raw reads were aligned to the mm10 genome and quantified via the MobiCube pipeline. High-quality cells were retained, normalized, and integrated using the Seurat package (v4.3.0). Major non-immune lineages were projected via UMAP and annotated using canonical marker expression: CAFs (Col1a1), MES-like Cells (Cd44, Hif1a), Vascular Cells (Pecam1), and Oligodendrocytes (Plp1). Differential expression analysis between two groups within the CAF and MES-like clusters was conducted using the *FindMarkers* function in Seurat with the default parameters.

## QUANTIFICATION AND STATISTICAL ANALYSIS

All data were analyzed and processed using R (v4.3.2) and Python (v3.10.12), with details of specific functions and libraries provided in the Methods sections. Statistical significance was determined by Mann-Whitney U test (Wilcoxon rank-sum test), Fisher’s exact test, Student’s t test, Proportion test, Chi-squared test, Pearson Correlation Coefficient, and Spearman Correlation Coefficient, with a p-value < 0.05 considered statistically significant. Survival comparisons were quantified using log rank Mantel-Cox test and Cox proportional hazards model.

## Supplementary Figure Legends

**Figure S1.**
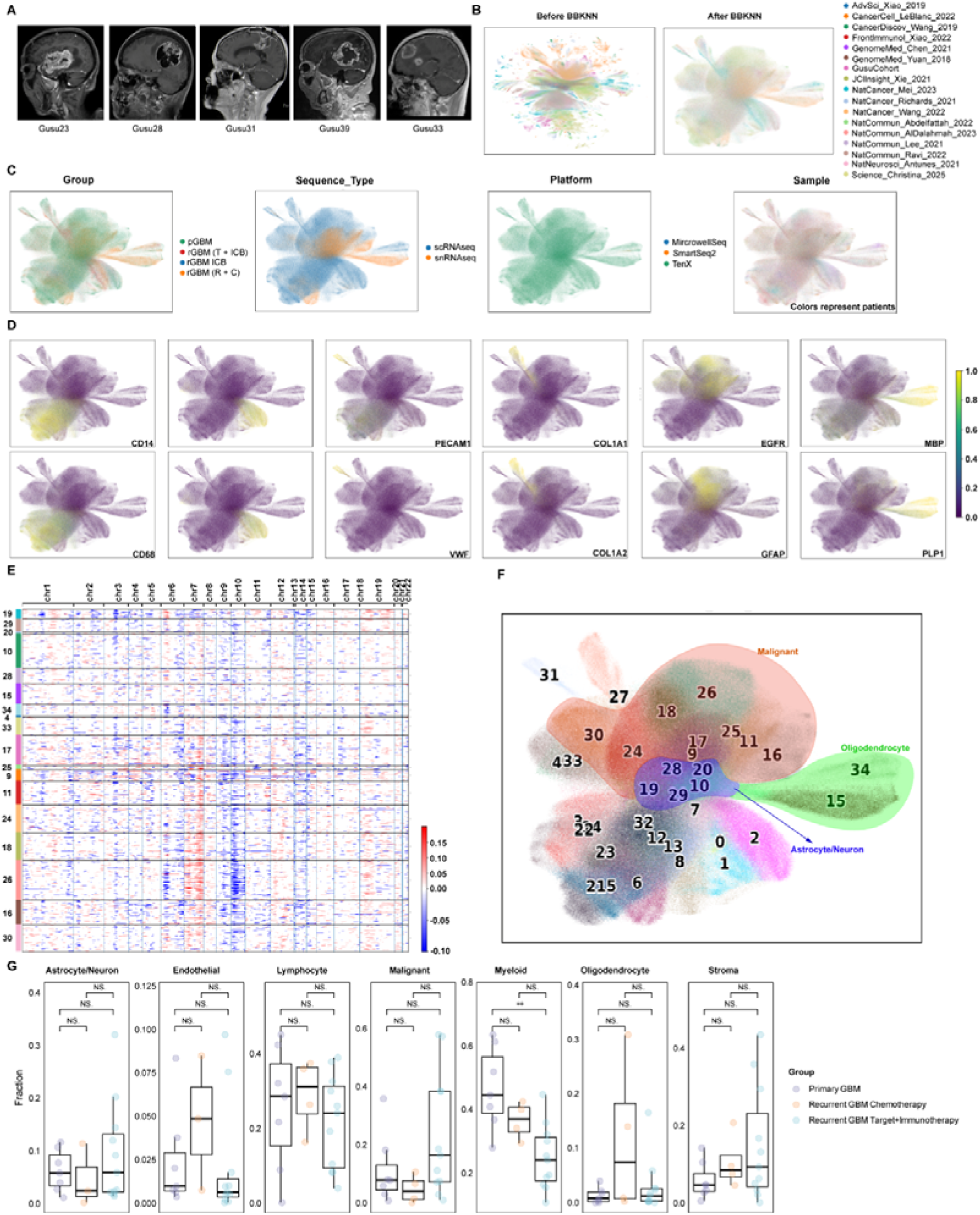
Quality Control, Batch Correction, and Cell Type Annotation of the GRIT-Atlas. **(A)** Representative sagittal contrast-enhanced MRI scans of five patients from the in-house Gusu cohort. **(B)** UMAP visualization of the aggregate dataset before (left) and after (right) batch effect correction using BBKNN. **(C)** Global UMAP embeddings colored by clinical treatment group, sequencing modality (snRNA-seq vs. scRNA-seq), sequencing platform, and individual sample origin. **(D)** Feature plots displaying the expression of representative marker genes for the seven major integrated cell types. **(E)** Heatmap of inferred Copy Number Variations (InferCNV) used to separate malignant cells from normal cell types, utilizing oligodendrocytes as the diploid reference. **(F)** UMAP projection distinguishing malignant cells (defined by CNV) from normal astrocytes/center neurons and oligodendrocytes. **(G)** Bar plots comparing cell-type proportions among the three therapeutic stratification groups specifically within the independent Mei et al. validation dataset. See also Figure 1.

**Figure S2.**
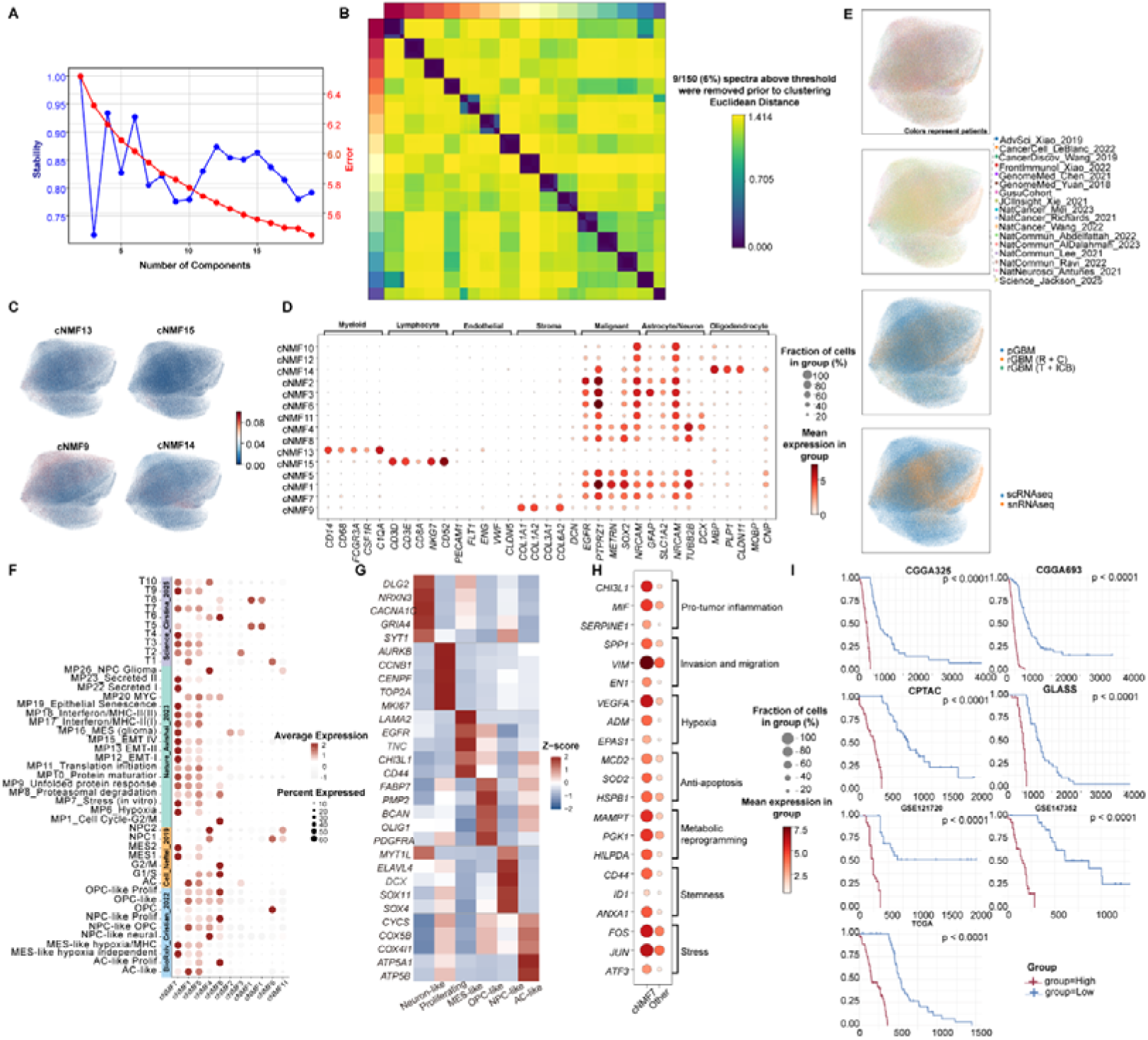
Robustness of cNMF Factorization and Clinical Validation of the cNMF7 Signature. **(A)** Line plots tracking model stability and reconstruction error across a range of factorization ranks ($K$), guiding the selection of the optimal decomposition value. **(B)** Consensus clustering heatmap for the selected rank, demonstrating the reproducibility of cluster assignments across independent factorization replicates. **(C, D)** Technical noise filter profiling, showing UMAP feature plots **(C)** and marker gene dot plots **(D)** characterizing the four excluded cNMF components associated with background artifacts. **(E)** UMAP embeddings colored by sample origin, batch, treatment cohort, and sequencing modality to confirm the absence of technical batch effects in the final alignment. **(F)** Dot plot correlating the identified cNMF meta-programs with previously established glioblastoma cellular states based on AUCell enrichment scores. **(G)** Transcriptional classification heatmap grouping individual cNMF programs into broader cellular phenotypes based on signature gene expression. **(H)** Dot plot comparing the expression of core functional networks—spanning inflammation, invasion, hypoxia, anti-apoptosis, metabolism, stemness, and stress—between the cNMF7 state and remaining malignant populations. **(I)** Kaplan-Meier overall survival curves validating the clinical prognostic significance of the cNMF7 program across seven independent patient cohorts, stratified by ssGSEA scores. Log-rank test. See also Figure 2.

**Figure S3.**
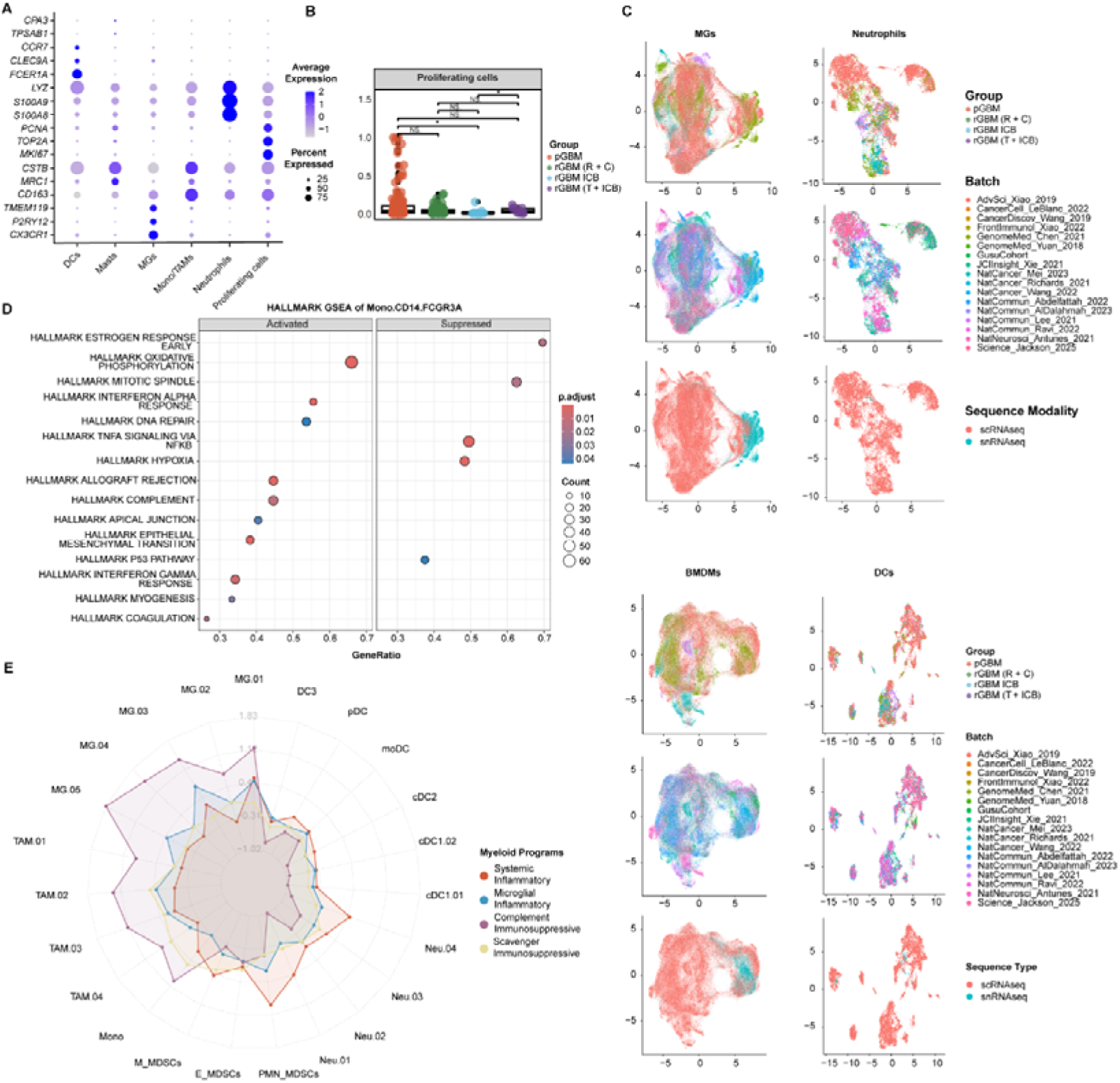
Quality Control and Functional Validation of Myeloid Sub-Classifications. **(A)** Dot plot visualizing the expression of canonical lineage-defining markers used to resolve the major myeloid categories. **(B)** Box plot comparing the proportional abundance of proliferating myeloid cells (normalized to total CD45+ cells) across treatment cohorts. Wilcoxon rank-sum test. **(C)** Myeloid UMAP embeddings colored by sample group, batch, and sequencing modality, confirming effective integration. **(D)** Gene Set Enrichment Analysis (GSEA) bubble plot highlighting the functional signaling pathways enriched within the Mono.CD16.FCGR3A monocyte subset. **(E)** Radar plot mapping the functional orientation of all fine-grained myeloid subsets onto four reference microenvironmental programs (microglial inflammatory, scavenger immunosuppressive, systemic inflammatory, and complement immunosuppressive). See also Figure 3.

**Figure S4. Divergent Functional States and Treatment-Associated Dynamics of T and NK Lymphocytes (A)** Fine-grained UMAP visualization showing the sub-clustering of four major lymphoid lineages: CD4+ T cells, CD8+ Memory/Naïve T cells, CD8+ Cytotoxic/Exhausted T cells, and Natural Killer (NK) cells.

**(B)** Nested donut charts displaying the hierarchical composition of each lymphoid lineage relative to the total lymphoid compartment.

**(C)** Heatmaps illustrating the expression of distinct marker genes defining the sub-clusters for each fine-grained lineage.

**(D)** Signaling pathway activity heatmap displaying DecoupleR-inferred scores for 14 pathways across the identified lymphoid subsets.

**(E, F)** Lymphoid state transitions across treatment stages tracked via an alluvial plot **(E)** and an aligned Ro/e enrichment heatmap **(F)**.

**(G)** CytoTRACE2 differentiation potential of NK cell subsets projected onto the NK UMAP space.

**(H)** Heatmap comparing the functional strength of Cytotoxicity and NK cell signature modules between CD8+ CTL/Tex and NK cell populations using AUCell scores.

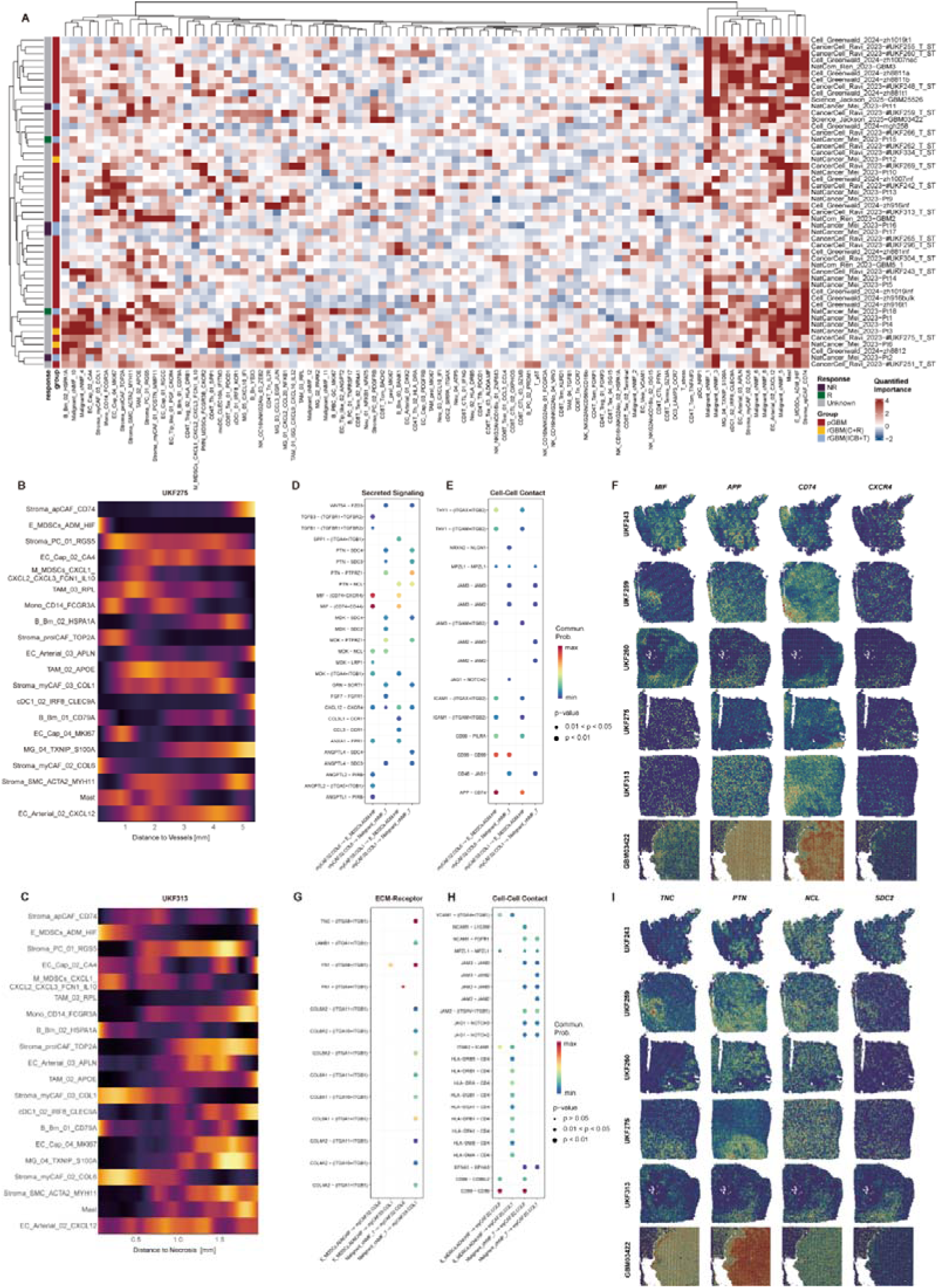

**Figure S5.**
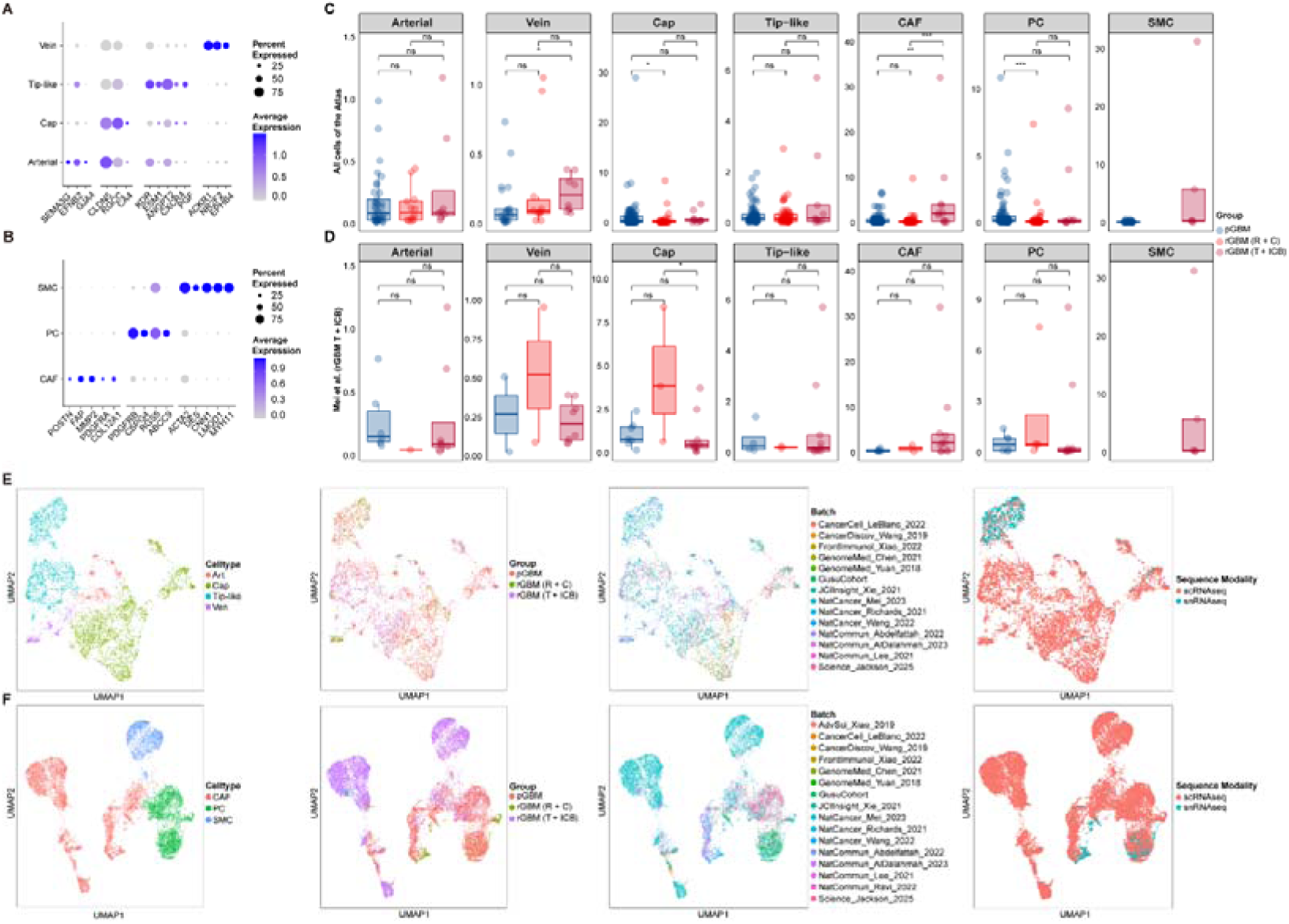
Global Characterization and Quality Assessment of the Lymphoid Compartment. **(A)** Global UMAP embedding of the aggregate lymphoid compartment, annotated by major lineages. **(B)** Dot plot showing the expression of canonical marker genes used to define broad lymphoid categories. **(C)** Box plots comparing the relative abundance of broad lymphoid lineages (normalized to CD45+ cells) across the four treatment groups. Wilcoxon rank-sum test. **(D)** Lymphoid integration assessment UMAPs colored by Sample Group, Batch, Sequencing Modality, and medium-resolution cell annotations. **(E)** Feature plots validating the molecular basis of NK cell sub-classification, highlighting the expression of *KLRC1*, *NCAM1*, and *FCGR3A*. **(F)** Box plots quantifying the proportional distribution of medium-resolution lymphoid subsets within the CD45+ compartment across treatment arms. Wilcoxon rank-sum test. See also Figure S4.

**Figure S6.**
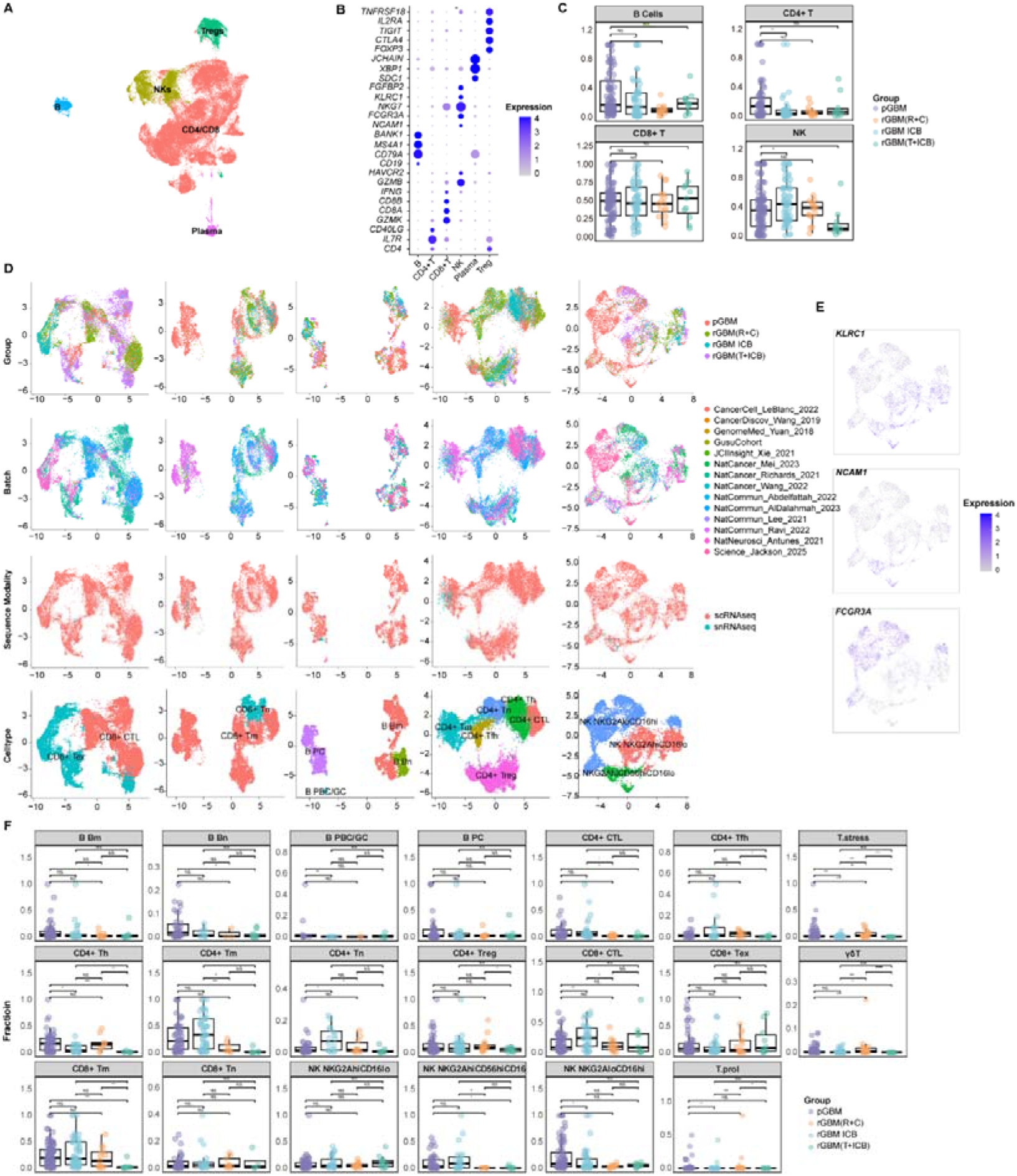
Annotation Resolution and Quality Assessment of Vascular and Stromal Cells. **(A, B)** Dot plots displaying the expression of lineage-defining markers for medium-resolution annotations of **(A)** Endothelial cell subtypes and **(B)** Stromal cell lineages. **(C)** Box plots comparing the proportional abundance of Stromal cells across treatment groups, utilizing non-sorted datasets to exclude technical collection bias. Wilcoxon rank-sum test. **(D)** Verification of cell type proportional trends across treatment groups specifically within the independent Mei et al. dataset. **(E, F)** Quality control and batch integration UMAP embeddings for the **(E)** Endothelial and **(F)** Stromal compartments, colored by annotation, sample group, batch, and sequencing modality. See also Figure 4.

**Figure S7.**
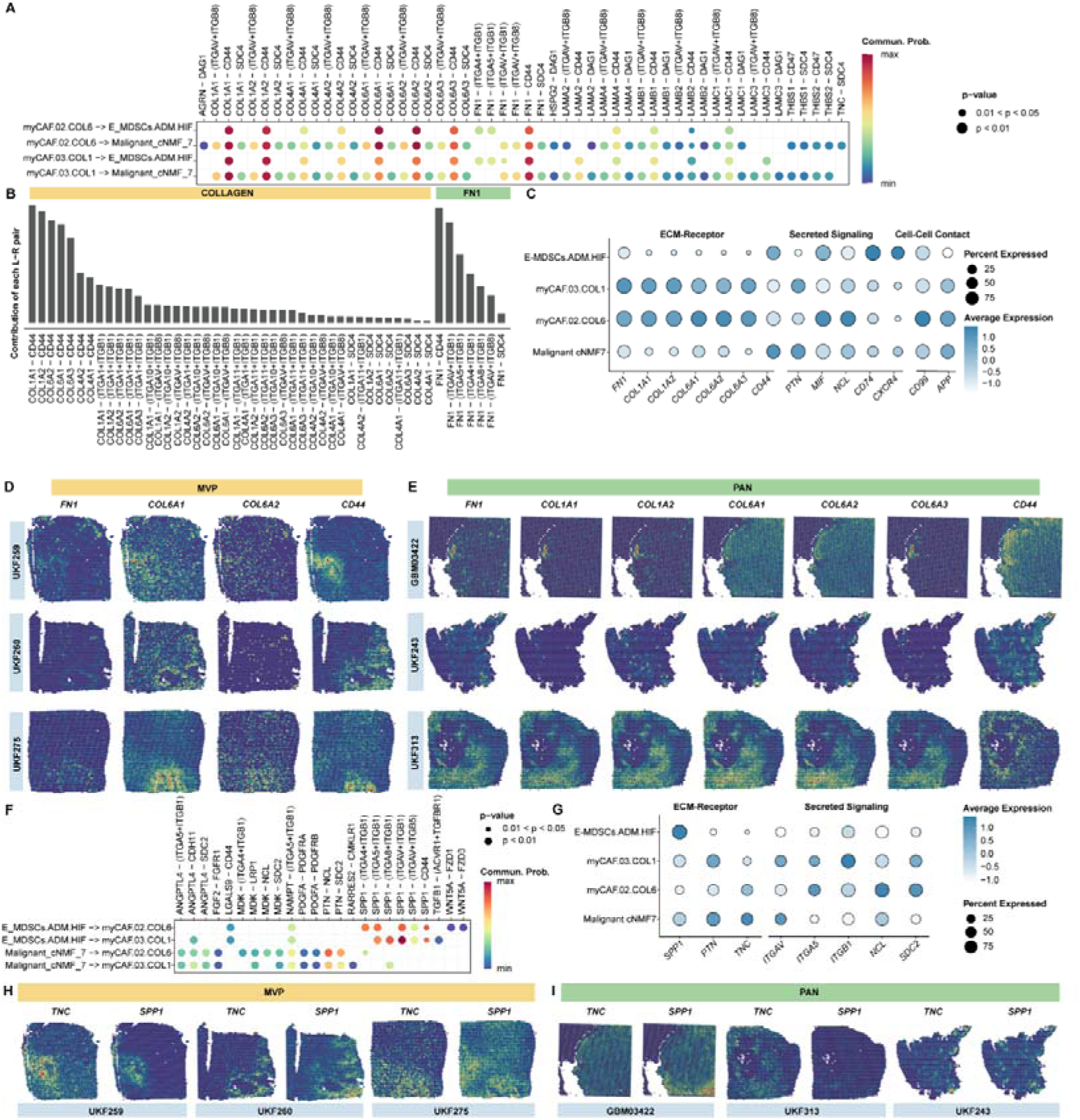
Robustness of Spatial Co-Localization Inference and Niche-Specific Enrichment. **(A)** Comprehensive heatmap of MISTy importance scores across all 48 spatial transcriptomic samples, reporting the predictive spatial importance of individual cell types for the distribution of the cNMF7 subpopulation. **(B, C)** Extended spatial trajectory heatmaps **(B)** and distance-dependent line plots **(C)** modeling the abundance trends relative to MVP (left) and PAN (right) cores for all cell types within the identified spatial modules. **(D, E)** CellChat dot plots displaying significant **(D)** Secreted Signaling and **(E)** Cell-Cell Contact interactions originating from myCAFs to the microenvironment. **(F)** Spatial feature plots visualizing the transcript co-expression and localization of validated signaling molecules (*MIF, APP, CD74, CXCR4*) involved in myCAF-myeloid crosstalk. **(G, H)** CellChat dot plots displaying significant incoming **(G)** ECM-Receptor and **(H)** Cell-Cell Contact interactions received by myCAFs. **(I)** Spatial feature plots of additional paracrine mediators (*TNC, PTN, NCL, SDC2*) validating their enrichment across histopathological niches. See also Figure 5.

**Figure S8.**
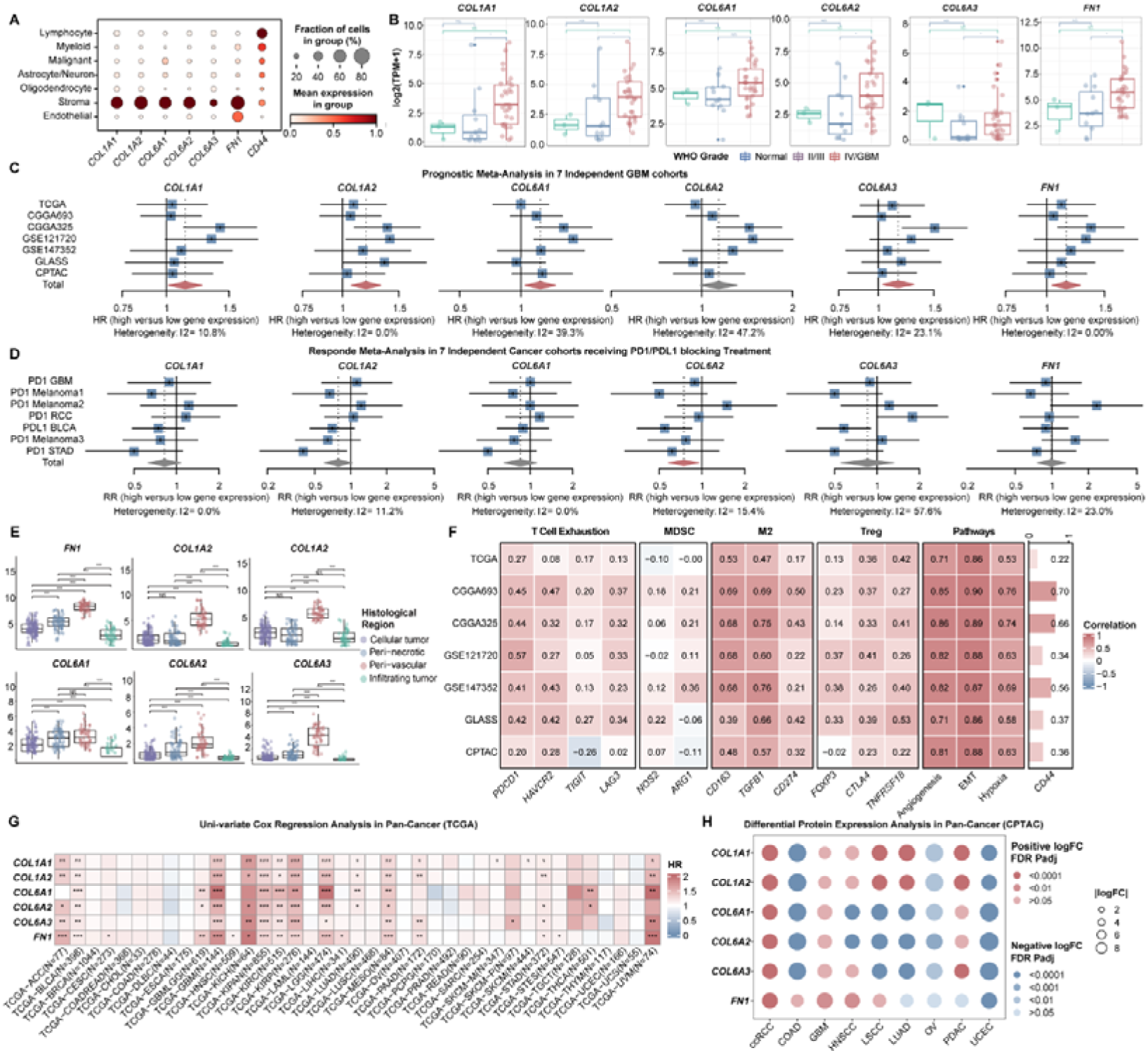
The ECM-CD44 Axis and Reciprocal Paracrine Signaling Sustain the Spatial Resistance Triad. **(A)** CellChat dot plot visualizing ECM-mediated communication probabilities from myCAFs (sender) to cNMF7 malignant cells and MDSCs (receivers). **(B)** Bar plots quantifying the relative contribution of individual ligand-receptor pairs to the dominant COLLAGEN (left) and FN1 (right) signaling pathways. **(C)** Expression dot plot of key ligand-receptor pairs grouped by interaction category across the triad components. **(D, E)** Spatial feature plots displaying the expression of core matrix ligands (*FN1, COL6A1, COL6A2, COL1A1, COL1A2, COL6A3*) and their cognate receptor (*CD44*) across independent Visium sections containing **(D)** MVP and **(E)** PAN structures. **(F)** Dot plot summarizing incoming Secreted Signaling interactions received by myCAFs from other TME components. **(G)** Expression profile of ligand-receptor pairs mediating incoming signals to myCAFs across the triad cell types. **(H)** Spatial feature plots of paracrine mediators (*TNC, SPP1*) confirming specific regional enrichment and co-localization with the resistance triad in MVP (left) versus PAN (right) niches. See also Figure 5.

**Figure S9.**
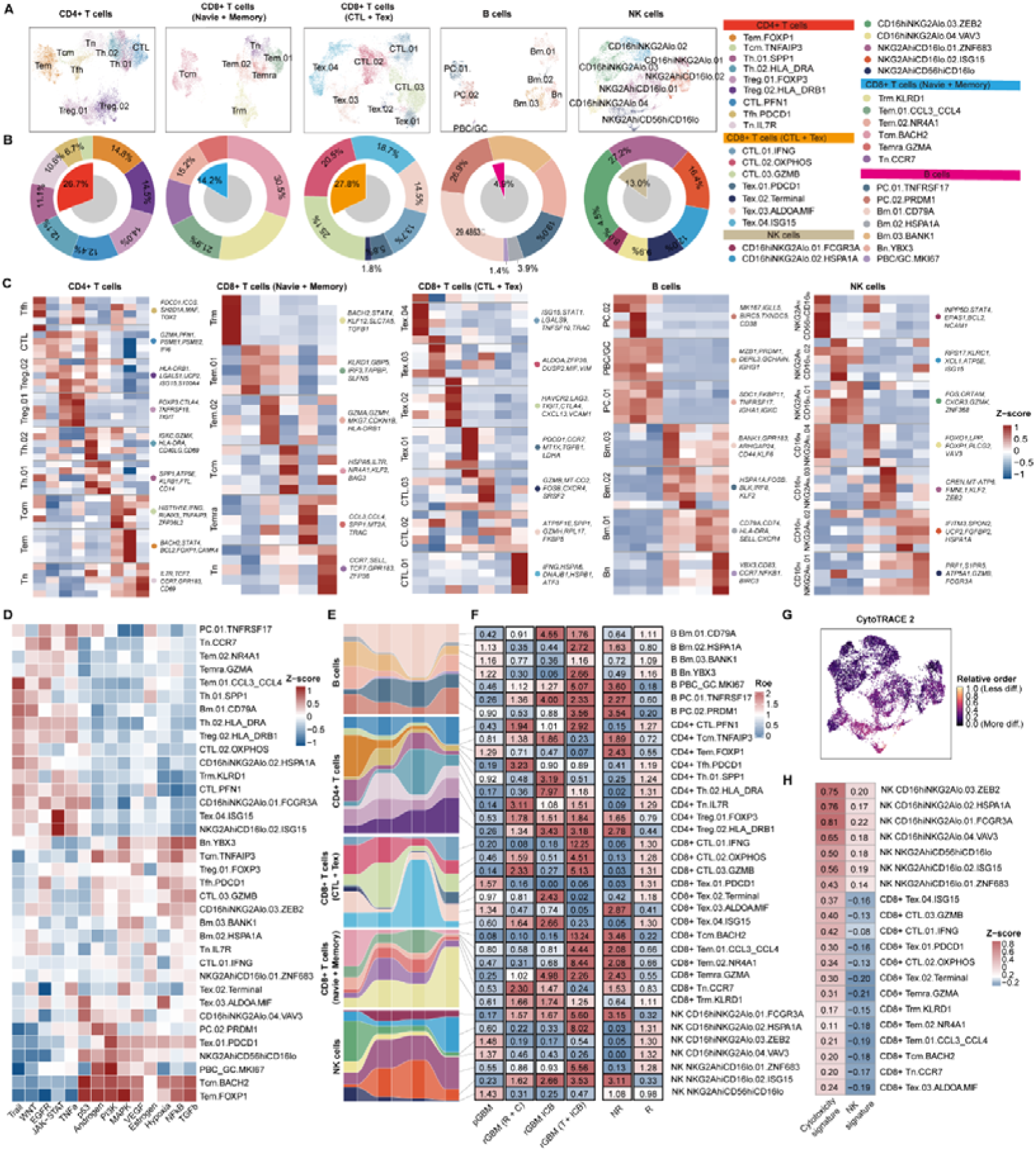
Clinical Relevance, Prognostic Value, and Pan-Cancer Validation of the Resistance-Associated ECM Signature. **(A)** Dot plot validating the cell-type specificity of the core ECM signature genes across the seven major lineages of the GRIT-Atlas. **(B)** Box plots comparing bulk RNA-seq expression levels of individual ECM components between normal brain tissue and gliomas of varying WHO grades within the in-house Gusu cohort. **(C)** Forest plots summarizing the meta-analysis of overall survival associations (Hazard Ratios and 95% confidence intervals) for each ECM gene across seven independent glioma cohorts using a random-effects model. **(D)** Forest plots summarizing the meta-analysis of immunotherapy response (Risk Ratios for objective response) evaluated across diverse clinical immunotherapy cohorts. **(E)** Box plots displaying the expression of ECM signature genes across micro-dissected anatomical regions from the Ivy Glioblastoma Atlas Project, confirming peak enrichment in peri-vascular and peri-necrotic zones. **(F)** Spearman correlation heatmap checking the association between individual ECM genes and key immune markers (T-cell exhaustion, MDSCs, M2-like TAMs, Tregs) and hallmark functional pathways across seven validation cohorts. **(G)** Pan-cancer survival heatmap displaying univariate Cox regression Hazard Ratios for the six ECM genes across 33 TCGA cancer types. **(H)** Pan-cancer proteomic validation dot plot visualizing the differential protein expression of the ECM signature genes across 10 distinct malignancies from the CPTAC dataset. See also Figure S7.

**Figure S10.**
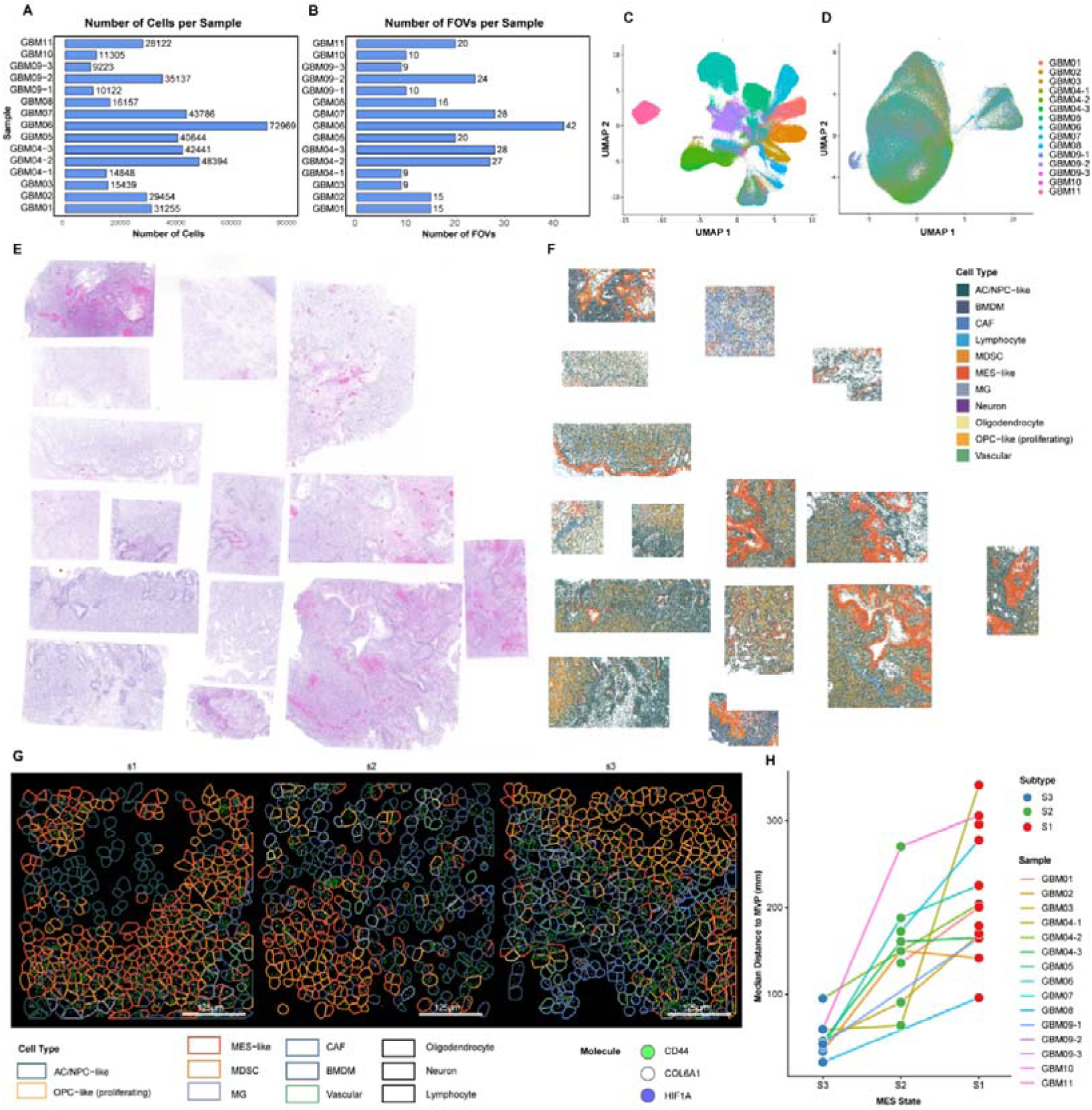
Quality Control, Cohort Harmonization, and Spatial Validation of MES-like Niches. **(A, B)** Bar plots reporting the total number of single cells recovered after quality filtering **(A)** and the total number of fields-of-view (FOVs) analyzed **(B)** per individual clinical sample on the CosMx platform. **(C, D)** Global UMAP visualization of the 406,689 single-cell spatial dataset prior to **(C)** and following **(D)** batch effect correction, colored by individual sample. **(E)** Serial histological sections stained with H&E capturing the global tissue architecture of the 15 spatial validation specimens. **(F)** Global spatial distribution maps of annotated cell lineages corresponding to the exact coordinates shown in (E). **(G)** High-resolution in situ transcript co-expression maps highlighting the localization of *HIF1A* (blue), *COL6A1* (white), and *CD44* (green) RNA transcripts within niches s1, s2, and s3. Scale bars = 125 μm. **(H)** Line plot displaying the median distance of individual cells within the three spatial MES-like states to the nearest MVP structure across individual clinical samples, mapping a consistent spatial gradient. See also Figure 6.

**Figure S11. Characterization of Patient-Matched Cellular Models, Matrix-Induced Stemness Cascades, and High-Throughput Anti-Fibrotic Small-Molecule Screening Metrics (A)** Contrast-enhanced T1-weighted MRI scans of the six glioblastoma patients selected for line derivation (left), alongside Western blot profiling and densitometric quantification of baseline CD44 protein expression across the newly established GSC lines.

**(B, C)** Western blot tracking of core neural stemness markers (*CD133* and *SOX2*) in primary lines GSC01 **(B)** and GSC06 **(C)** following 24-hour challenge with six candidate recombinant matrix proteins.

**(D, E)** Western blot analysis measuring downstream SOX2 expression changes in primary lines GSC01 **(D)** and GSC06 **(E)** subjected to pre-treatment with an anti-CD44 neutralizing antibody prior to ligand exposure.

**(F)** Parallel brightfield H&E overviews and high-plex immunofluorescence cross-sections comparing clinical specimens stratified into low versus high expression of COL6A1.

**(G)** Western blot analysis evaluating the protein expression of lineage markers (*COL6A1, FAP,* _α_*-SMA*) across the six primary CAF lines relative to their parallel patient-matched GSC lines.

**(H, I)** Quantitative Western blot bar plots **(H)** and matching colorimetric DAB immunohistochemical images **(I)** validating Col6a1 depletion levels in a stable shRNA knockdown model. Scale bar = 50 μm.

**(J)** ELISA screening concentration profiles of extracellular COL6A1 secretion across 24 candidate physical compounds utilizing primary CAF06 cells. See also Figure 7.

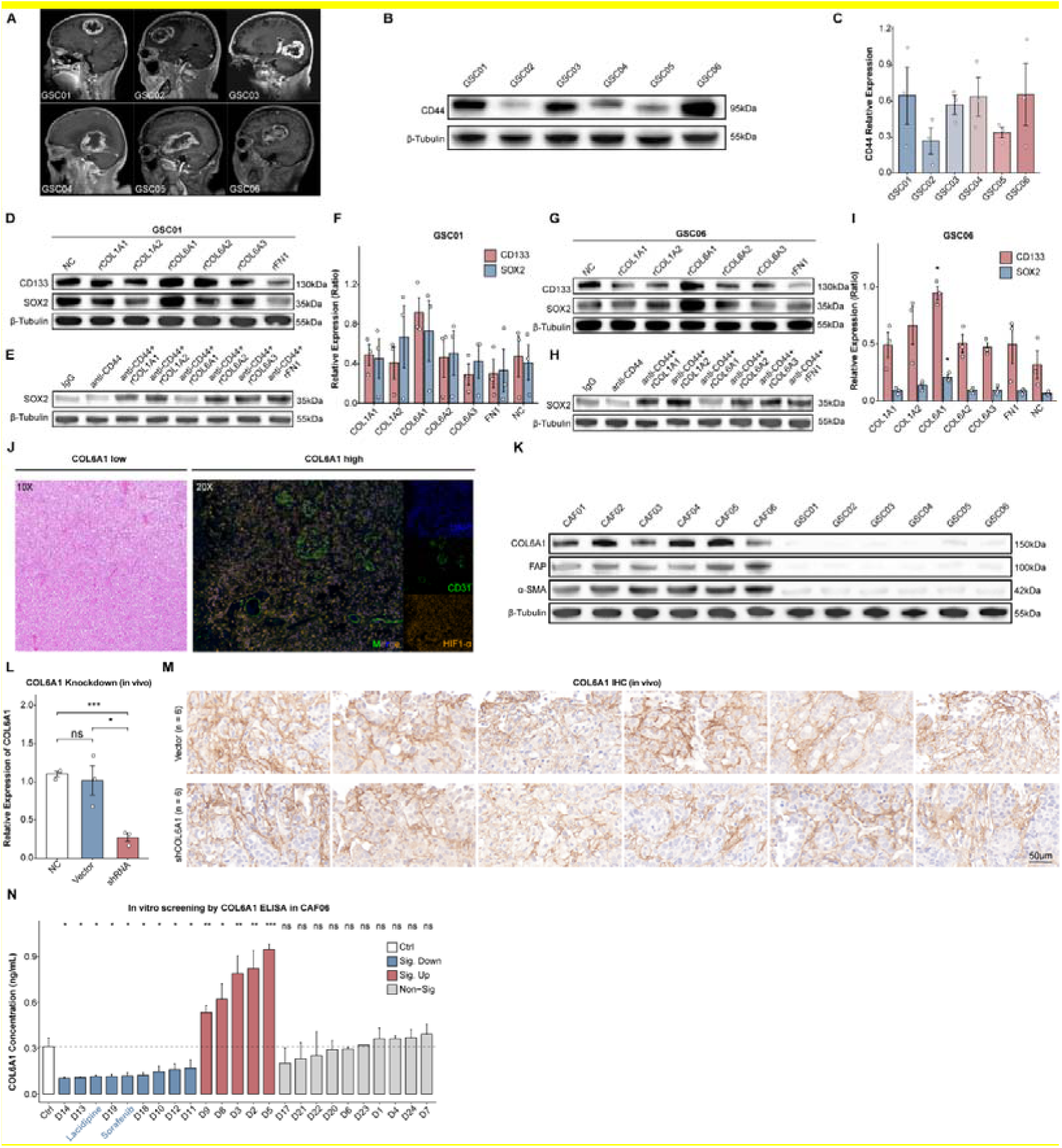

**Figure S12.**
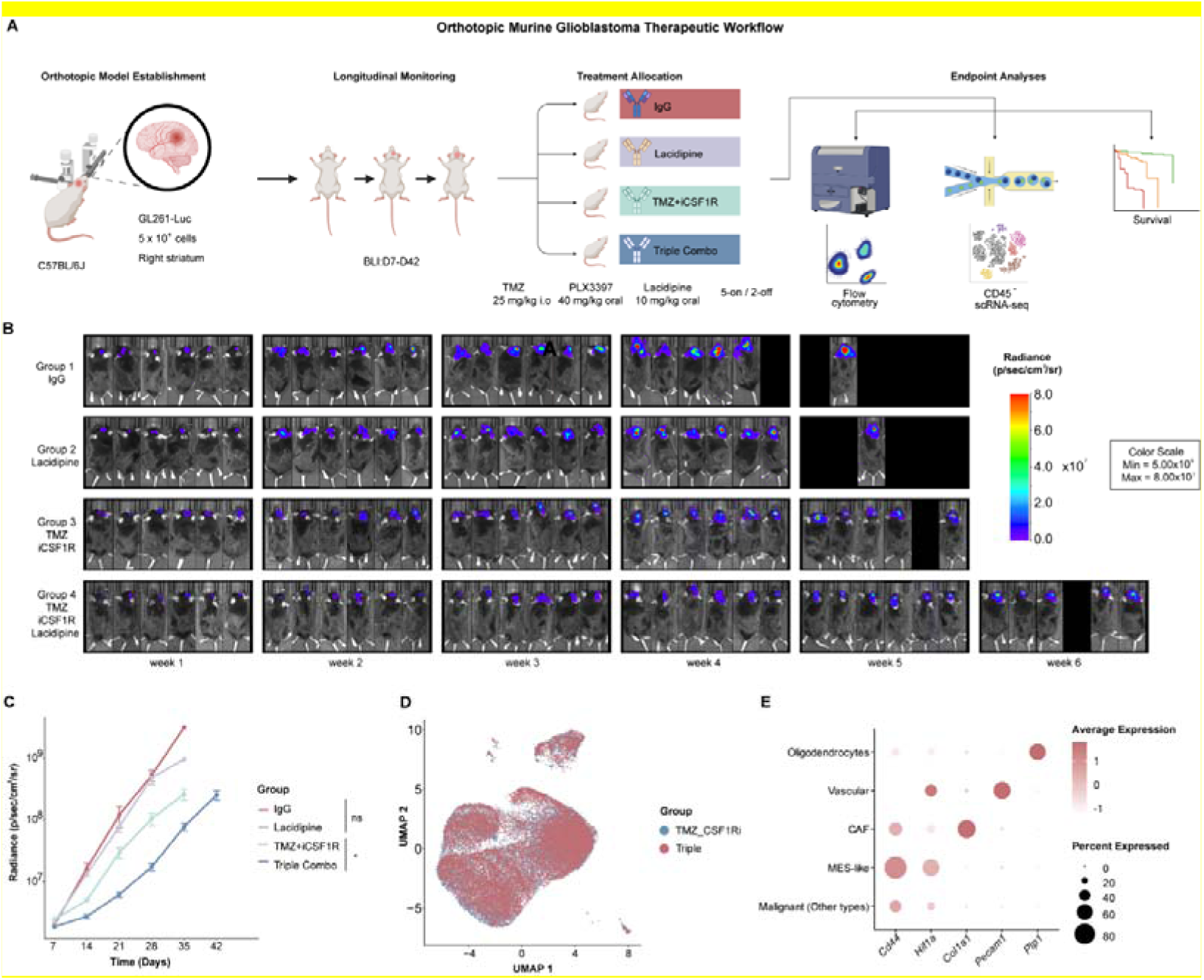
Preclinical Murine Optimization Pipeline, Longitudinal Bioluminescence Tumor Dynamics, and Single-Cell Transcriptomic Quality Controls of the Non-Immune Landscape. **(A)** Comprehensive experimental schematic outlining the orthotopic murine glioblastoma therapeutic workflow, detailing cell inoculation, longitudinal bioluminescence imaging (BLI), drug scheduling, and downstream multi-parametric endpoint processing. **(B)** Longitudinal in vivo BLI panels capturing consecutive intracranial tumor burden changes across a six-week timeline in representative mice belonging to the four experimental arms, presented on an absolute luminescence radiance scale. **(C)** In vivo growth kinetics curves tracking total luminescent radiance accumulation as a function of time across the therapeutic cohorts. **(D)** Seurat-integrated UMAP overlay tracking the overlapping distribution of isolated CD45- non-immune cells from the Doublet combo and Triple combo cohorts, demonstrating multi-sample batch alignment. **(E)** Single-cell dot plot characterization displaying the transcript expression intensity and population percentage of canonical lineage markers across the isolated non-immune clusters. See also Figure 7.

## Supplementary Table Legends

**Table S1. Summary of Datasets and Patient Clinicopathological Metadata.** Catalog of the integrated GRIT-Atlas cohort demographics, including sequencing modalities, sample counts, and treatment stratification across 17 single-cell/nucleus RNA-seq datasets, 48 spatial transcriptomics sections, and independent bulk RNA-seq cohorts.

**Table S2. Differentially Expressed Genes (DEGs) for All Cell Subpopulations.** Comprehensive list of cluster-specific marker genes, statistical significance metrics, and expression fractions resolved across all major malignant, myeloid, lymphoid, stromal, and endothelial cell lineages.

**Table S3. Curated Representative Marker Genes for Major Cell Lineages and Subpopulations.** Filtered selection of highly specific canonical marker genes utilized for high-fidelity phenotypic classification and characterization of individual fine-grained subclusters.

**Table S4. Summary of Published Gene Signatures for Cell Type Benchmarking and Validation.** Compendium of reference gene expression signatures extracted from landmark glioblastoma and pan-cancer studies, compiled to quantitatively validate the cell-state annotations of malignant, myeloid, endothelial, and stromal compartments.

**Table S5. Spatial Inter-Cellular Association Scores Calculated via MISTy Across 48 Patient Sections.** Matrix of localized spatial importance values modeling the recurrent geographical co-localization or exclusion patterns between the cNMF7 mesenchymal-like malignant subpopulation and neighboring microenvironmental cell types.

**Table S6. Patient Clinicopathological Metadata for the mIHC and CosMx SMI Validation Cohorts.** Comprehensive registry of the validation cohort, documenting patient demographics (Gender, Age), tumor characteristics (WHO Grade, Histology, IDH Status), treatment history, alongside molecular (HIF1A/CD31) and histopathological validation of MVP and PAN architectures, and tissue allocation for mIHC and CosMx SMI 6k profiling.

**Table S7. Downsampled and Balanced Spatial Niche Assignment Matrix from CosMx High-Plex Imaging.** Cellular neighborhood intensity and composition matrix, optimized via downsampling and balancing, used to construct spatial architecture domains across integrated clinical tissue specimens.

**Table S8. Differential Expression Analysis Across the Three Spatial Niches.** Statistical outputs and differential transcriptomic profiles characterizing the microenvironmental features of the three identified spatial niches (s1, s2, and s3) within the mesenchymal-like domains.

**Table S9. Profile of the 24 Candidate Small-Molecule Compounds for Anti-Fibrotic Screening.** Chemical documentation, compound classification, identifiers, and targeted treatment metrics for the 24 candidate physical small molecules evaluated in the high-throughput extracellular matrix secretion screen.

**Table S10. Quantitative Proteomic Matrix and Differential Abundance Analysis of Lacidipine-Treated CAFs.** High-throughput 4D-DIA quantitative proteomics dataset profiling patient-derived primary cancer-associated fibroblasts (CAF01) across a three-versus-three replication design of Lacidipine treatment (L1–L3) versus vehicle control (N1–N3).

**Table S11. In Vivo Single-Cell Transcriptomic DEGs of Murine CAF and Malignant MES Populations.** Differential expression signatures mapping treatment-responsive transcriptional shifts within the CAFs and MES-like malignant cell compartments isolated from orthotopic murine models.

## Data and Code Availability

### Data availability

The Glioblastoma Resistance Insights from Treatment Atlas (GRIT-Atlas) represents a comprehensive single-cell and spatially resolved transcriptomic resource for IDH-wildtype glioblastoma, with all processed data objects deposited at Zenodo and publicly available as of the date of publication (https://doi.org/10.5281/zenodo.18009414/20686853). This repository encompasses the master Seurat/AnnData object capturing 978,065 high-quality cells, high-resolution sub-lineage objects for the malignant, myeloid, lymphoid, stromal, and endothelial compartments, as well as the RCTD deconvolution results and MistyR spatial importance matrices for 48 Visium sections. Furthermore, it contains sample-specific Seurat v5 objects for the CosMx SMI high-plex spatial transcriptomics data fully annotated with global cell-type identities, specialized Seurat v5 objects focused exclusively on the MES-like malignant domains with descriptions of the four spatial architectural niches (S1–S3), and the standard expression files for the in vivo mouse glioblastoma scRNA-seq cohort validating preclinical therapeutic responses. Primary raw sequencing data utilized for this atlas were sourced from public repositories including GEO, EGA, and SRA, with specific accession numbers and metadata for the 296 integrated samples provided in Table S1.

### Code availability

All original code and computational pipelines developed for the construction and analysis of the GRIT-Atlas have been deposited at GitHub and are publicly available as of the date of publication (https://github.com/Dr-fly/GRIT-Atlas). The repository contains detailed, reproducible workflows implemented in Python (v3.10.12) and R (v4.3.2 and v5.3.0) environments, covering standardized single-cell quality control and preprocessing, BBKNN-based multi-batch integration, malignant functional state characterization via consensus NMF, spatial transcriptomic deconvolution using RCTD, ntercellular communication modeling via CellChat and high-plex spatial neighborhood composition and architectural niche modeling for the CosMx SMI dataset. A stable, version-controlled release of the code specifically used to generate the final results and figures reported in this paper is also archived at Zenodo alongside the processed data objects.

## Acknowledgements

Not applicable.

## Author contributions

**Conceptualization:** Fei Wang, Zhouqing Chen, Zhong Wang. **Data curation:** Fei Wang, Xin Wu, Guozheng Zhao, Run Huang. **Formal analysis:** Fei Wang, Xin Wu, Guozheng Zhao, Run Huang, Chen Yang, Yuhan Bai. **Funding acquisition:** Zhouqing Chen, Zhong Wang, Fei Wang. **Investigation:** Fei Wang, Xin Wu, Guozheng Zhao, Run Huang, Chen Yang, Yuhan Bai, Wenqian Cao, Yue Lu, Guangling Xu, Haohao Qiu, Hongyi Ling, Dengfeng Lu, Youjia Qiu, Juyi Zhang, Bixi Gao. **Methodology:** Fei Wang, Xin Wu, Guozheng Zhao, Run Huang, Yanbo Yang, Ting Sun. **Project administration:** Zhouqing Chen, Zhong Wang. **Resources:** Zhouqing Chen, Zhong Wang, Ting Sun. **Software:** Fei Wang, Xin Wu, Run Huang, Chen Yang. **Supervision:** Zhouqing Chen, Zhong Wang. **Validation:** Fei Wang, Xin Wu, Guozheng Zhao, Run Huang, Wenqian Cao, Yue Lu. **Visualization:** Fei Wang, Xin Wu, Guozheng Zhao, Run Huang. **Writing – original draft:** Fei Wang, Xin Wu, Guozheng Zhao, Run Huang. **Writing – review & editing:** Zhouqing Chen, Zhong Wang, Fei Wang, Yanbo Yang, Ting Sun.

## Funding

This work was supported by the Innovative Technology Class A grant of the First Affiliated Hospital of Soochow University [grant number 0499980301001], National Natural Science Foundation of China [grant number 82571477] and National Natural Science Foundation of China [grant number 82201445].

## Notes

https://github.com/Dr-fly/GRIT-Atlas

https://zenodo.org/records/18009414

https://zenodo.org/records/20686853

### Competing Interest Statement

The authors have declared no competing interest.

### Summary of Updates

The latest version of the repository and dataset exhibits a profound upgrade in its multi-omic resolution, validation depth, and translational focus. Mechanistically, the narrative has shifted from a generalized description of the cellular "Spatial Resistance Triad" to a highly precise mapping of the spatiotemporal MVP-PAN (Microvascular Proliferation to Pseudopalisading Necrosis) evolutionary continuum, unmasking a deterministic desmoplastic dependency wherein stromal-derived COL6A1 constructs a mechanical structural sanctuary that engages CD44 on MES-like malignant cells to sustain tumor stemness during vascular collapse. To achieve true single-cell spatial resolution for this phenomenon, the previous 10x Visium spatial transcriptomics core has been seamlessly augmented with high-plex CosMx Spatial Molecular Imaging (SMI) data encompassing 406,689 individual cells, supported computationally by the newly added Module 5 script for advanced neighborhood composition extraction and consensus niche clustering (S1 to S3). Furthermore, the clinical discovery cohorts have been robustly reinforced with extensive preclinical validation; the archived data now hosts the standard expression triplet files for an orthotopic mouse glioblastoma single-cell sequencing cohort. This in vivo model serves to nominate and validate Lacidipine, an FDA-approved, blood-brain barrier-penetrant calcium channel blocker, as the lead translational compound capable of dismantling the desmoplastic matrix scar by blocking calcium-mediated CAF secretion. Finally, the entire pipeline has been upgraded to native Seurat v5 structural compatibility to handle these multi-modal, high-plex spatial arrays, and all author contributions have been rigorously overhauled and standardized according to the explicit CRediT guidelines.

## Reference

1. Miller, K.D., Ostrom, Q.T., Kruchko, C., Patil, N., Tihan, T., Cioffi, G., Fuchs, H.E., Waite, K.A., Jemal, A., Siegel, R.L., and Barnholtz-Sloan, J.S. (2021). Brain and other central nervous system tumor statistics, 2021. CA Cancer J Clin 71, 381–406. 10.3322/caac.21693.

2. Wen, P.Y., Weller, M., Lee, E.Q., Alexander, B.M., Barnholtz-Sloan, J.S., Barthel, F.P., Batchelor, T.T., Bindra, R.S., Chang, S.M., Chiocca, E.A., et al. (2020). Glioblastoma in adults: a Society for Neuro-Oncology (SNO) and European Society of Neuro-Oncology (EANO) consensus review on current management and future directions. Neuro Oncol 22, 1073–1113. 10.1093/neuonc/noaa106.

3. Schaff, L.R., and Mellinghoff, I.K. (2023). Glioblastoma and Other Primary Brain Malignancies in Adults: A Review. Jama 329, 574–587. 10.1001/jama.2023.0023.

4. Yabo, Y.A., Niclou, S.P., and Golebiewska, A. (2022). Cancer cell heterogeneity and plasticity: A paradigm shift in glioblastoma. Neuro Oncol 24, 669–682. 10.1093/neuonc/noab269.

5. Lin, H., Liu, C., Hu, A., Zhang, D., Yang, H., and Mao, Y. (2024). Understanding the immunosuppressive microenvironment of glioma: mechanistic insights and clinical perspectives. J Hematol Oncol 17, 31. 10.1186/s13045-024-01544-7.

6. Hendriksen, J.D., Locallo, A., Maarup, S., Debnath, O., Ishaque, N., Hasselbach, B., Skjøth-Rasmussen, J., Yde, C.W., Poulsen, H.S., Lassen, U., and Weischenfeldt, J. (2024). Immunotherapy drives mesenchymal tumor cell state shift and TME immune response in glioblastoma patients. Neuro Oncol 26, 1453–1466. 10.1093/neuonc/noae085.

7. Gupta, P., Dang, M., Oberai, S., Migliozzi, S., Trivedi, R., Kumar, G., Peshoff, M., Milam, N., Ahmed, A., Bojja, K., et al. (2024). Immune landscape of isocitrate dehydrogenase-stratified primary and recurrent human gliomas. Neuro Oncol 26, 2239–2255. 10.1093/neuonc/noae139.

8. Bunse, L., Bunse, T., Kilian, M., Quintana, F.J., and Platten, M. (2025). The immunology of brain tumors. Sci Immunol 10, eads0449. 10.1126/sciimmunol.ads0449.

9. Mahdi, J., Trivedi, V., and Monje, M. (2025). The promise of immunotherapy for central nervous system tumours. Nat Rev Immunol. 10.1038/s41577-025-01227-5.

10. Spitzer, A., Johnson, K.C., Nomura, M., Garofano, L., Nehar-Belaid, D., Darnell, N.G., Greenwald, A.C., Bussema, L., Oh, Y.T., Varn, F.S., et al. (2025). Deciphering the longitudinal trajectories of glioblastoma ecosystems by integrative single-cell genomics. Nat Genet 57, 1168–1178. 10.1038/s41588-025-02168-4.

11. Nomura, M., Spitzer, A., Johnson, K.C., Garofano, L., Nehar-Belaid, D., Galili Darnell, N., Greenwald, A.C., Bussema, L., Oh, Y.T., Varn, F.S., et al. (2025). The multilayered transcriptional architecture of glioblastoma ecosystems. Nat Genet 57, 1155–1167. 10.1038/s41588-025-02167-5.

12. Lee, A.H., Sun, L., Mochizuki, A.Y., Reynoso, J.G., Orpilla, J., Chow, F., Kienzler, J.C., Everson, R.G., Nathanson, D.A., Bensinger, S.J., et al. (2021). Neoadjuvant PD-1 blockade induces T cell and cDC1 activation but fails to overcome the immunosuppressive tumor associated macrophages in recurrent glioblastoma. Nat Commun 12, 6938. 10.1038/s41467-021-26940-2.

13. Mei, Y., Wang, X., Zhang, J., Liu, D., He, J., Huang, C., Liao, J., Wang, Y., Feng, Y., Li, H., et al. (2023). Siglec-9 acts as an immune-checkpoint molecule on macrophages in glioblastoma, restricting T-cell priming and immunotherapy response. Nat Cancer 4, 1273–1291. 10.1038/s43018-023-00598-9.

14. Pombo Antunes, A.R., Scheyltjens, I., Lodi, F., Messiaen, J., Antoranz, A., Duerinck, J., Kancheva, D., Martens, L., De Vlaminck, K., Van Hove, H., et al. (2021). Single-cell profiling of myeloid cells in glioblastoma across species and disease stage reveals macrophage competition and specialization. Nat Neurosci 24, 595–610. 10.1038/s41593-020-00789-y.

15. Poon, C.C., Herbrich, S.M., Chen, Y., Hossain, A., Fuller, G.N., Jindal, S., Basu, S., Ledbetter, D., Macaluso, M., Phillips, L.M., et al. (2025). Mesenchymal Stem Cells and Fibroblasts Contribute to Microvascular Proliferation in Glioblastoma and are Correlated with Immunosuppression and Poor Outcome. Cancer Immunol Res 13, 804–820. 10.1158/2326-6066.Cir-24-0743.

16. Migliozzi, S., Adabbo, B., Garofano, L., Wu, F., Davila, P., Komotar, R.J., Ivan, M.E., Shah, A.H., Currall, B.B., Williams, S., et al. (2025). Restraint of cancer cell plasticity by spatial homotypic clustering. Cancer Cell 43, 2206–2223.e2210. 10.1016/j.ccell.2025.08.009.

17. Artzi, S.B., Klausen, M.N., Harwood, D.S.L., Michaelsen, S.R., Maarup, S.B., Locallo, A., Fougner, V., Bager, N.S., Hammouda, N.M., Nørøxe, D.S., et al. (2025). Spatial transcriptomic analysis reveals lack of response to PD-1 blockade in recurrent glioblastoma. Acta Neuropathol 150, 29. 10.1007/s00401-025-02937-9.

18. Puchalski, R.B., Shah, N., Miller, J., Dalley, R., Nomura, S.R., Yoon, J.G., Smith, K.A., Lankerovich, M., Bertagnolli, D., Bickley, K., et al. (2018). An anatomic transcriptional atlas of human glioblastoma. Science 360, 660–663. 10.1126/science.aaf2666.

19. Huse, J.T., and Holland, E.C. (2010). Targeting brain cancer: advances in the molecular pathology of malignant glioma and medulloblastoma. Nat Rev Cancer 10, 319–331. 10.1038/nrc2818.

20. Brennan, C.W., Verhaak, R.G., McKenna, A., Campos, B., Noushmehr, H., Salama, S.R., Zheng, S., Chakravarty, D., Sanborn, J.Z., Berman, S.H., et al. (2013). The somatic genomic landscape of glioblastoma. Cell 155, 462–477. 10.1016/j.cell.2013.09.034.

21. Garofano, L., Migliozzi, S., Oh, Y.T., D’Angelo, F., Najac, R.D., Ko, A., Frangaj, B., Caruso, F.P., Yu, K., Yuan, J., et al. (2021). Pathway-based classification of glioblastoma uncovers a mitochondrial subtype with therapeutic vulnerabilities. Nat Cancer 2, 141–156. 10.1038/s43018-020-00159-4.

22. Kim, K.H., Migliozzi, S., Koo, H., Hong, J.H., Park, S.M., Kim, S., Kwon, H.J., Ha, S., Garofano, L., Oh, Y.T., et al. (2024). Integrated proteogenomic characterization of glioblastoma evolution. Cancer Cell 42, 358–377.e358. 10.1016/j.ccell.2023.12.015.

23. Varn, F.S., Johnson, K.C., Martinek, J., Huse, J.T., Nasrallah, M.P., Wesseling, P., Cooper, L.A.D., Malta, T.M., Wade, T.E., Sabedot, T.S., et al. (2022). Glioma progression is shaped by genetic evolution and microenvironment interactions. Cell 185, 2184–2199.e2116. 10.1016/j.cell.2022.04.038.

24. Ma, T., Chu, X., Wang, J., Li, X., Zhang, Y., Tong, D., Xu, W., Dang, G., Qi, L., Miao, Y., et al. (2025). Pan-Cancer Analyses Refine the Single-Cell Portrait of Tumor-Infiltrating Dendritic Cells. Cancer Res 85, 3596–3613. 10.1158/0008-5472.Can-24-3595.

25. Jackson, C., Cherry, C., Bom, S., Dykema, A.G., Wang, R., Thompson, E., Zhang, M., Li, R., Ji, Z., Hou, W., et al. (2025). Distinct myeloid-derived suppressor cell populations in human glioblastoma. Science 387, eabm5214. 10.1126/science.abm5214.

26. Wang, W., Li, T., Cheng, Y., Li, F., Qi, S., Mao, M., Wu, J., Liu, Q., Zhang, X., Li, X., et al. (2024). Identification of hypoxic macrophages in glioblastoma with therapeutic potential for vasculature normalization. Cancer Cell 42, 815–832.e812. 10.1016/j.ccell.2024.03.013.

27. Du, Y., Long, X., Li, X., Guan, F., Gao, W., Deng, K., Wang, S., Lin, X., Huang, M., She, X., et al. (2025). Spatial-reprogramming derived GPNMB(+) macrophages interact with COL6A3(+) fibroblasts to enhance vascular fibrosis in glioblastoma. Genome Med 17, 136 10.1186/s13073-025-01553-2.

28. Cakmak, P., Lun, J.H., Singh, A., Macas, J., Schupp, J., Schuck, J., Mahmoud, Z., Köhler, M., Starzetz, T., Burger, M.C., et al. (2025). Spatial immune profiling defines a subset of human gliomas with functional tertiary lymphoid structures. Immunity 58, 2847–2863.e2848. 10.1016/j.immuni.2025.09.018.

29. Nabors, L.B., Supko, J.G., Rosenfeld, M., Chamberlain, M., Phuphanich, S., Batchelor, T., Desideri, S., Ye, X., Wright, J., Gujar, S., and Grossman, S.A. (2011). Phase I trial of sorafenib in patients with recurrent or progressive malignant glioma. Neuro Oncol 13, 1324–1330. 10.1093/neuonc/nor145.

30. Galanis, E., Anderson, S.K., Lafky, J.M., Uhm, J.H., Giannini, C., Kumar, S.K., Kimlinger, T.K., Northfelt, D.W., Flynn, P.J., Jaeckle, K.A., et al. (2013). Phase II study of bevacizumab in combination with sorafenib in recurrent glioblastoma (N0776): a north central cancer treatment group trial. Clin Cancer Res 19, 4816–4823. 10.1158/1078-0432.Ccr-13-0708.

31. McCormack, P.L., and Wagstaff, A.J. (2003). Lacidipine: a review of its use in the management of hypertension. Drugs 63, 2327–2356. 10.2165/00003495-200363210-00008.

32. Jain, S., Rick, J.W., Joshi, R.S., Beniwal, A., Spatz, J., Gill, S., Chang, A.C., Choudhary, N., Nguyen, A.T., Sudhir, S., et al. (2023). Single-cell RNA sequencing and spatial transcriptomics reveal cancer-associated fibroblasts in glioblastoma with protumoral effects. J Clin Invest 133 10.1172/jci147087.

33. Chen, C., Zhao, S., Karnad, A., and Freeman, J.W. (2018). The biology and role of CD44 in cancer progression: therapeutic implications. J Hematol Oncol 11, 64. 10.1186/s13045-018-0605-5.

34. Johansson, E., Grassi, E.S., Pantazopoulou, V., Tong, B., Lindgren, D., Berg, T.J., Pietras, E.J., Axelson, H., and Pietras, A. (2017). CD44 Interacts with HIF-2α to Modulate the Hypoxic Phenotype of Perinecrotic and Perivascular Glioma Cells. Cell Rep 20, 1641–1653. 10.1016/j.celrep.2017.07.049.

35. Pietras, A., Katz, A.M., Ekström, E.J., Wee, B., Halliday, J.J., Pitter, K.L., Werbeck, J.L., Amankulor, N.M., Huse, J.T., and Holland, E.C. (2014). Osteopontin-CD44 signaling in the glioma perivascular niche enhances cancer stem cell phenotypes and promotes aggressive tumor growth. Cell Stem Cell 14, 357–369. 10.1016/j.stem.2014.01.005.

36. Turtoi, A., Blomme, A., Bianchi, E., Maris, P., Vannozzi, R., Naccarato, A.G., Delvenne, P., De Pauw, E., Bevilacqua, G., and Castronovo, V. (2014). Accessibilome of human glioblastoma: collagen-VI-alpha-1 is a new target and a marker of poor outcome. J Proteome Res 13, 5660–5669. 10.1021/pr500657w.

37. Cescon, M., Rampazzo, E., Bresolin, S., Da Ros, F., Manfreda, L., Cani, A., Della Puppa, A., Braghetta, P., Bonaldo, P., and Persano, L. (2023). Collagen VI sustains cell stemness and chemotherapy resistance in glioblastoma. Cell Mol Life Sci 80, 233. 10.1007/s00018-023-04887-5.

38. Galbo, P.M., Jr., Madsen, A.T., Liu, Y., Peng, M., Wei, Y., Ciesielski, M.J., Fenstermaker, R.A., Graff, S., Montagna, C., Segall, J.E., et al. (2024). Functional Contribution and Clinical Implication of Cancer-Associated Fibroblasts in Glioblastoma. Clin Cancer Res 30, 865–876. 10.1158/1078-0432.Ccr-23-0493.

39. Stupp, R., Mason, W.P., van den Bent, M.J., Weller, M., Fisher, B., Taphoorn, M.J., Belanger, K., Brandes, A.A., Marosi, C., Bogdahn, U., et al. (2005). Radiotherapy plus concomitant and adjuvant temozolomide for glioblastoma. N Engl J Med 352, 987–996. 10.1056/NEJMoa043330.

40. Nassiri, F., Patil, V., Yefet, L.S., Singh, O., Liu, J., Dang, R.M.A., Yamaguchi, T.N., Daras, M., Cloughesy, T.F., Colman, H., et al. (2023). Oncolytic DNX-2401 virotherapy plus pembrolizumab in recurrent glioblastoma: a phase 1/2 trial. Nat Med 29, 1370–1378. 10.1038/s41591-023-02347-y.

41. Wolock, S.L., Lopez, R., and Klein, A.M. (2019). Scrublet: Computational Identification of Cell Doublets in Single-Cell Transcriptomic Data. Cell Syst 8, 281–291.e289. 10.1016/j.cels.2018.11.005.

42. Mathewson, N.D., Ashenberg, O., Tirosh, I., Gritsch, S., Perez, E.M., Marx, S., Jerby-Arnon, L., Chanoch-Myers, R., Hara, T., Richman, A.R., et al. (2021). Inhibitory CD161 receptor identified in glioma-infiltrating T cells by single-cell analysis. Cell 184, 1281–1298.e1226. 10.1016/j.cell.2021.01.022.

43. Neftel, C., Laffy, J., Filbin, M.G., Hara, T., Shore, M.E., Rahme, G.J., Richman, A.R., Silverbush, D., Shaw, M.L., Hebert, C.M., et al. (2019). An Integrative Model of Cellular States, Plasticity, and Genetics for Glioblastoma. Cell 178, 835–849.e821. 10.1016/j.cell.2019.06.024.

44. Sun, H.F., Li, L.D., Lao, I.W., Li, X., Xu, B.J., Cao, Y.Q., and Jin, W. (2022). Single-cell RNA sequencing reveals cellular and molecular reprograming landscape of gliomas and lung cancer brain metastases. Clin Transl Med 12, e1101. 10.1002/ctm2.1101.

45. Schmassmann, P., Roux, J., Dettling, S., Hogan, S., Shekarian, T., Martins, T.A., Ritz, M.F., Herter, S., Bacac, M., and Hutter, G. (2023). Single-cell characterization of human GBM reveals regional differences in tumor-infiltrating leukocyte activation. Elife 12. 10.7554/eLife.92678.

46. Müller, S., Di Lullo, E., Bhaduri, A., Alvarado, B., Yagnik, G., Kohanbash, G., Aghi, M., and Diaz, A. (2018). A single-cell atlas of human glioblastoma reveals a single axis of phenotype in tumor-propagating cells. bioRxiv, 377606. 10.1101/377606.

47. Wolf, F.A., Angerer, P., and Theis, F.J. (2018). SCANPY: large-scale single-cell gene expression data analysis. Genome Biol 19, 15. 10.1186/s13059-017-1382-0.

48. Polański, K., Young, M.D., Miao, Z., Meyer, K.B., Teichmann, S.A., and Park, J.E. (2020). BBKNN: fast batch alignment of single cell transcriptomes. Bioinformatics 36, 964–965. 10.1093/bioinformatics/btz625.

49. Zhou, Y., Zhou, B., Pache, L., Chang, M., Khodabakhshi, A.H., Tanaseichuk, O., Benner, C., and Chanda, S.K. (2019). Metascape provides a biologist-oriented resource for the analysis of systems-level datasets. Nat Commun 10, 1523. 10.1038/s41467-019-09234-6.

50. Patel, A.P., Tirosh, I., Trombetta, J.J., Shalek, A.K., Gillespie, S.M., Wakimoto, H., Cahill, D.P., Nahed, B.V., Curry, W.T., Martuza, R.L., et al. (2014). Single-cell RNA-seq highlights intratumoral heterogeneity in primary glioblastoma. Science 344, 1396–1401. 10.1126/science.1254257.

51. Zeng, Z., Ma, Y., Hu, L., Tan, B., Liu, P., Wang, Y., Xing, C., Xiong, Y., and Du, H. (2024). OmicVerse: a framework for bridging and deepening insights across bulk and single-cell sequencing. Nat Commun 15, 5983. 10.1038/s41467-024-50194-3.

52. Hao, Y., Hao, S., Andersen-Nissen, E., Mauck, W.M., 3rd, Zheng, S., Butler, A., Lee, M.J., Wilk, A.J., Darby, C., Zager, M., et al. (2021). Integrated analysis of multimodal single-cell data. Cell 184, 3573–3587.e3529. 10.1016/j.cell.2021.04.048.

53. Zhang, L., Yu, X., Zheng, L., Zhang, Y., Li, Y., Fang, Q., Gao, R., Kang, B., Zhang, Q., Huang, J.Y., et al. (2018). Lineage tracking reveals dynamic relationships of T cells in colorectal cancer. Nature 564, 268–272. 10.1038/s41586-018-0694-x.

54. Ruiz-Moreno, C., Salas, S.M., Samuelsson, E., Minaeva, M., Ibarra, I., Grillo, M., Brandner, S., Roy, A., Forsberg-Nilsson, K., Kranendonk, M.E.G., et al. (2025). Charting the single-cell and spatial landscape of IDH-wild-type glioblastoma with GBmap. Neuro Oncol 27, 2281–2295. 10.1093/neuonc/noaf113.

55. Gavish, A., Tyler, M., Greenwald, A.C., Hoefflin, R., Simkin, D., Tschernichovsky, R., Galili Darnell, N., Somech, E., Barbolin, C., Antman, T., et al. (2023). Hallmarks of transcriptional intratumour heterogeneity across a thousand tumours. Nature 618, 598–606. 10.1038/s41586-023-06130-4.

56. Miller, T.E., El Farran, C.A., Couturier, C.P., Chen, Z., D’Antonio, J.P., Verga, J., Villanueva, M.A., Gonzalez Castro, L.N., Tong, Y.E., Saadi, T.A., et al. (2025). Programs, origins and immunomodulatory functions of myeloid cells in glioma. Nature 640, 1072–1082. 10.1038/s41586-025-08633-8.

57. Xie, Y., Yang, F., He, L., Huang, H., Chao, M., Cao, H., Hu, Y., Fan, Z., Zhai, Y., Zhao, W., et al. (2024). Single-cell dissection of the human blood-brain barrier and glioma blood-tumor barrier. Neuron 112, 3089–3105.e3087. 10.1016/j.neuron.2024.07.026.

58. Xie, Y., He, L., Lugano, R., Zhang, Y., Cao, H., He, Q., Chao, M., Liu, B., Cao, Q., Wang, J., et al. (2021). Key molecular alterations in endothelial cells in human glioblastoma uncovered through single-cell RNA sequencing. JCI Insight 6. 10.1172/jci.insight.150861.

59. Gao, Y., Li, J., Cheng, W., Diao, T., Liu, H., Bo, Y., Liu, C., Zhou, W., Chen, M., Zhang, Y., et al. (2024). Cross-tissue human fibroblast atlas reveals myofibroblast subtypes with distinct roles in immune modulation. Cancer Cell 42, 1764–1783.e1710. 10.1016/j.ccell.2024.08.020.

60. Badia, I.M.P., Vélez Santiago, J., Braunger, J., Geiss, C., Dimitrov, D., Müller-Dott, S., Taus, P., Dugourd, A., Holland, C.H., Ramirez Flores, R.O., and Saez-Rodriguez, J. (2022). decoupleR: ensemble of computational methods to infer biological activities from omics data. Bioinform Adv 2, vbac016. 10.1093/bioadv/vbac016.

61. Xu, S., Hu, E., Cai, Y., Xie, Z., Luo, X., Zhan, L., Tang, W., Wang, Q., Liu, B., Wang, R., et al. (2024). Using clusterProfiler to characterize multiomics data. Nat Protoc 19, 3292–3320. 10.1038/s41596-024-01020-z.

62. Kang, M., Gulati, G.S., Brown, E.L., Qi, Z., Avagyan, S., Armenteros, J.J.A., Gleyzer, R., Zhang, W., Steen, C.B., D’Silva, J.P., et al. (2025). Improved reconstruction of single-cell developmental potential with CytoTRACE 2. Nat Methods 22, 2258–2263. 10.1038/s41592-025-02857-2.

63. Van de Sande, B., Flerin, C., Davie, K., De Waegeneer, M., Hulselmans, G., Aibar, S., Seurinck, R., Saelens, W., Cannoodt, R., Rouchon, Q., et al. (2020). A scalable SCENIC workflow for single-cell gene regulatory network analysis. Nat Protoc 15, 2247–2276. 10.1038/s41596-020-0336-2.

64. Ravi, V.M., Will, P., Kueckelhaus, J., Sun, N., Joseph, K., Salié, H., Vollmer, L., Kuliesiute, U., von Ehr, J., Benotmane, J.K., et al. (2022). Spatially resolved multi-omics deciphers bidirectional tumor-host interdependence in glioblastoma. Cancer Cell 40, 639–655.e613. 10.1016/j.ccell.2022.05.009.

65. Ren, Y., Huang, Z., Zhou, L., Xiao, P., Song, J., He, P., Xie, C., Zhou, R., Li, M., Dong, X., et al. (2023). Spatial transcriptomics reveals niche-specific enrichment and vulnerabilities of radial glial stem-like cells in malignant gliomas. Nat Commun 14, 1028. 10.1038/s41467-023-36707-6.

66. Greenwald, A.C., Darnell, N.G., Hoefflin, R., Simkin, D., Mount, C.W., Gonzalez Castro, L.N., Harnik, Y., Dumont, S., Hirsch, D., Nomura, M., et al. (2024). Integrative spatial analysis reveals a multi-layered organization of glioblastoma. Cell 187, 2485–2501.e2426. 10.1016/j.cell.2024.03.029.

67. Cable, D.M., Murray, E., Zou, L.S., Goeva, A., Macosko, E.Z., Chen, F., and Irizarry, R.A. (2022). Robust decomposition of cell type mixtures in spatial transcriptomics. Nat Biotechnol 40, 517–526. 10.1038/s41587-021-00830-w.

68. Tanevski, J., Flores, R.O.R., Gabor, A., Schapiro, D., and Saez-Rodriguez, J. (2022). Explainable multiview framework for dissecting spatial relationships from highly multiplexed data. Genome Biol 23, 97. 10.1186/s13059-022-02663-5.

69. Kueckelhaus, J., Frerich, S., Kada-Benotmane, J., Koupourtidou, C., Ninkovic, J., Dichgans, M., Beck, J., Schnell, O., and Heiland, D.H. (2024). Inferring histology-associated gene expression gradients in spatial transcriptomic studies. Nat Commun 15, 7280. 10.1038/s41467-024-50904-x.

70. Jin, S., Guerrero-Juarez, C.F., Zhang, L., Chang, I., Ramos, R., Kuan, C.H., Myung, P., Plikus, M.V., and Nie, Q. (2021). Inference and analysis of cell-cell communication using CellChat. Nat Commun 12, 1088. 10.1038/s41467-021-21246-9.

71. Balduzzi, S., Rücker, G., and Schwarzer, G. (2019). How to perform a meta-analysis with R: a practical tutorial. Evid Based Ment Health 22, 153–160. 10.1136/ebmental-2019-300117.

72. Beygelzimer, A., Kakade, S., and Langford, J. (2006). Cover trees for nearest neighbor. Proceedings of the 23rd international conference on Machine learning. Association for Computing Machinery.

73. DeBruine, Z.J., Pospisilik, J.A., and Triche, T.J. (2024). Fast and interpretable non-negative matrix factorization for atlas-scale single cell data. bioRxiv, 2021.2009.2001.458620. 10.1101/2021.09.01.458620.

74. Fu, L., Shi, S., Yi, J., Wang, N., He, Y., Wu, Z., Peng, J., Deng, Y., Wang, W., Wu, C., et al. (2024). ADMETlab 3.0: an updated comprehensive online ADMET prediction platform enhanced with broader coverage, improved performance, API functionality and decision support. Nucleic Acids Res 52, W422–w431. 10.1093/nar/gkae236.

