## Supplementary figures and images for "A Single-Cell and Spatial Atlas of MVP-PAN Evolution Reveals a Desmoplastic Dependency to Overcome Glioblastoma Resistance"

### Figure1.tif

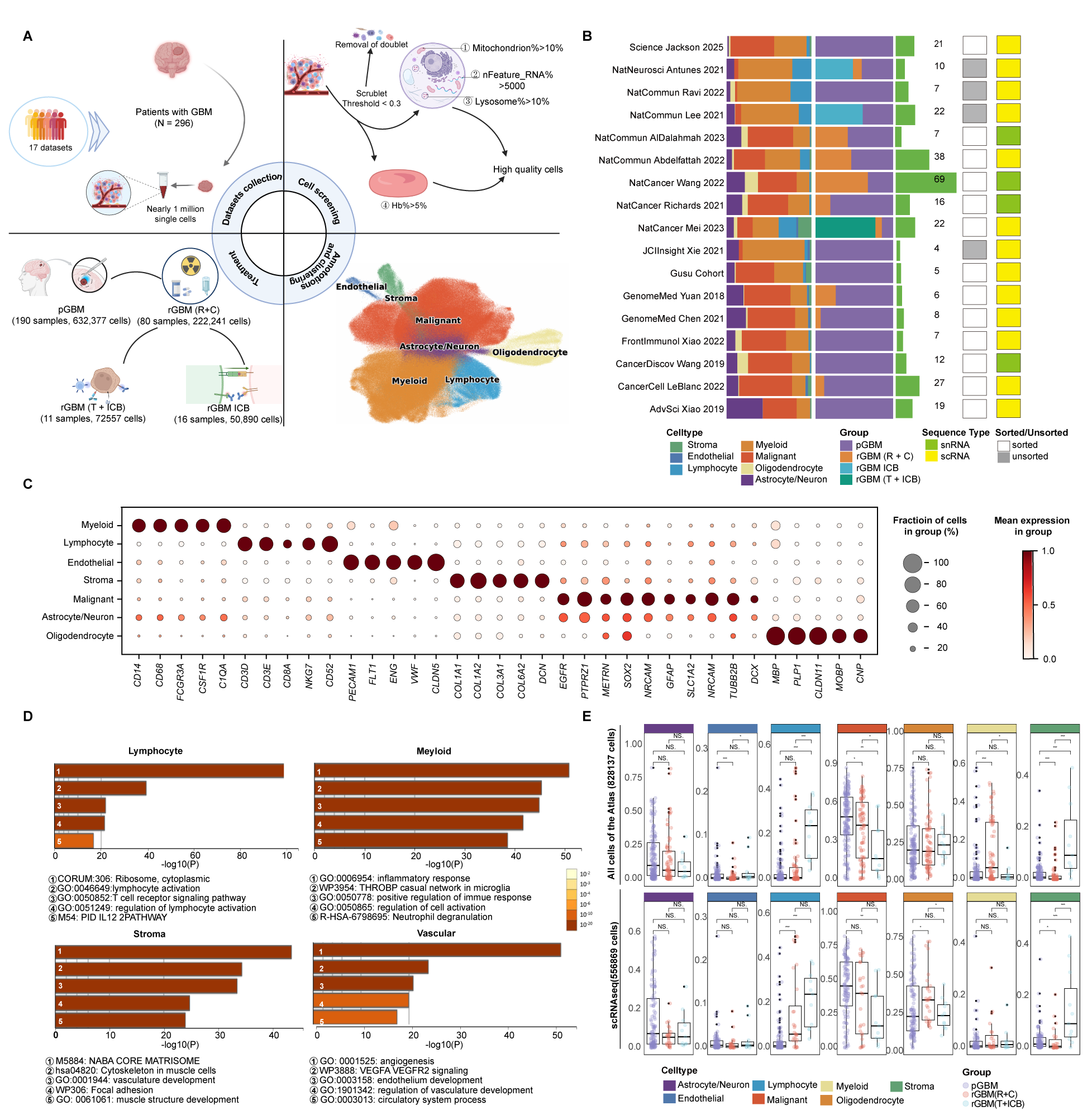

### Figure2_Malignant.tif

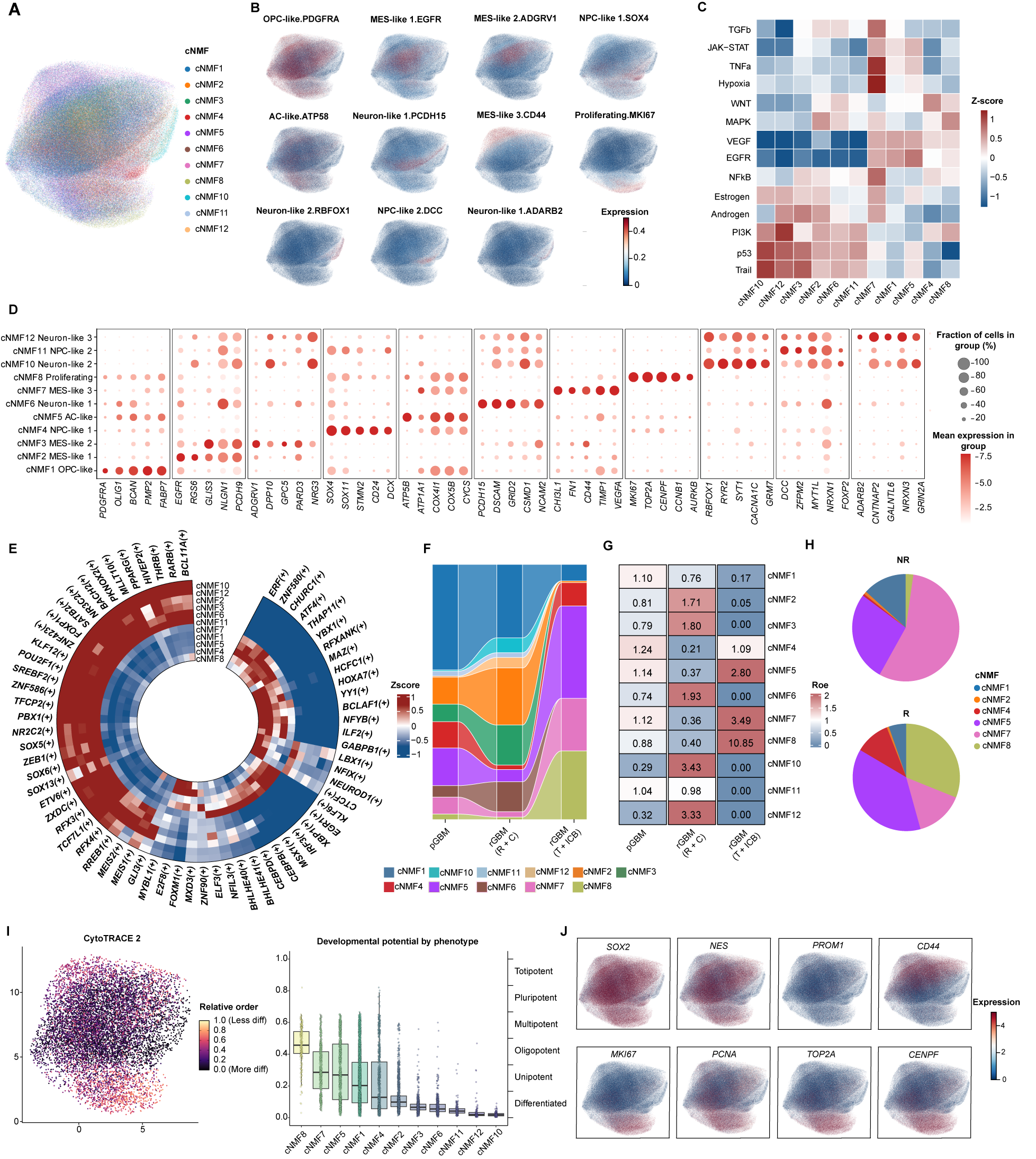

### Figure3_Myeloid.tif

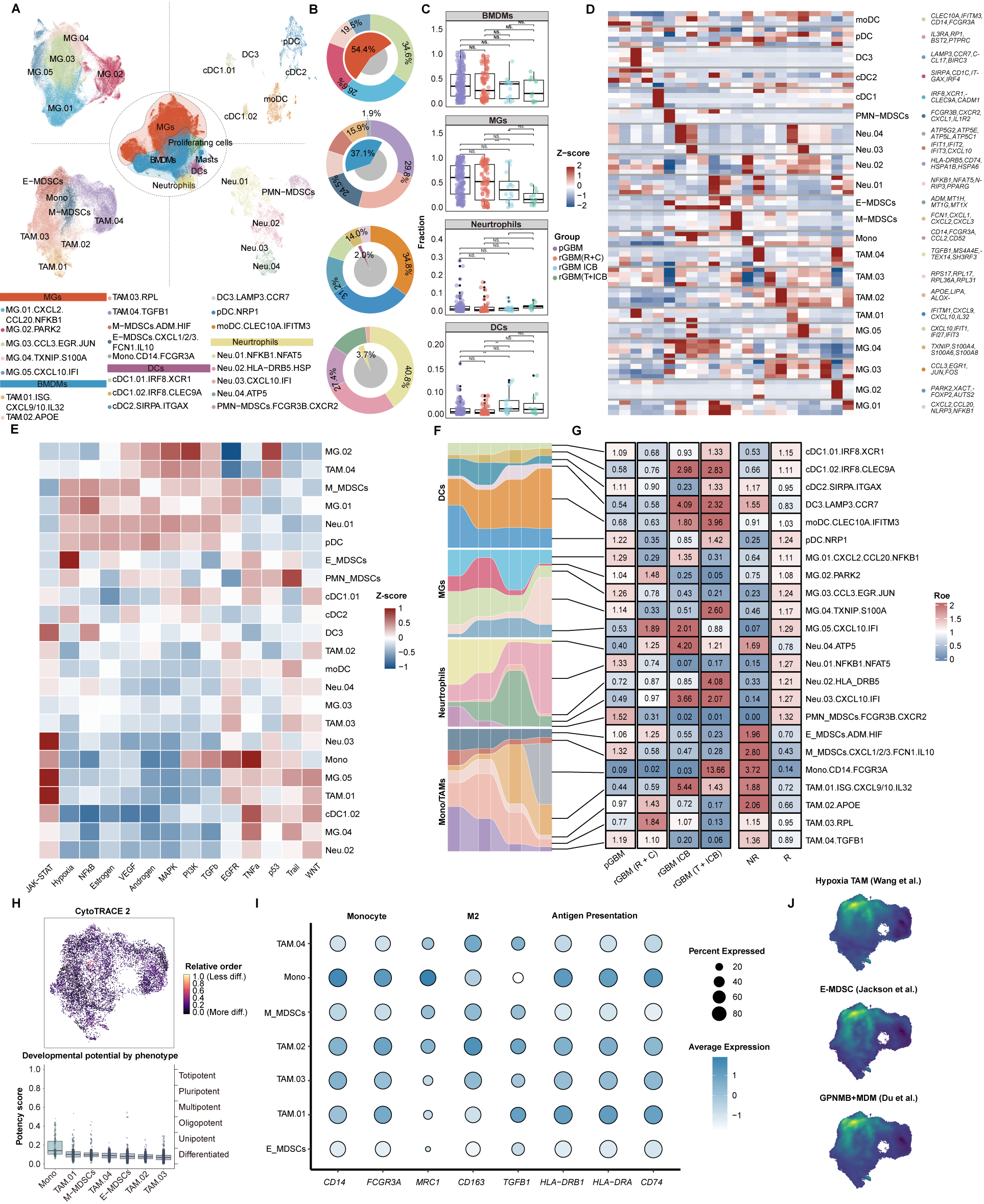

### Figure4_StromaVascular.tif

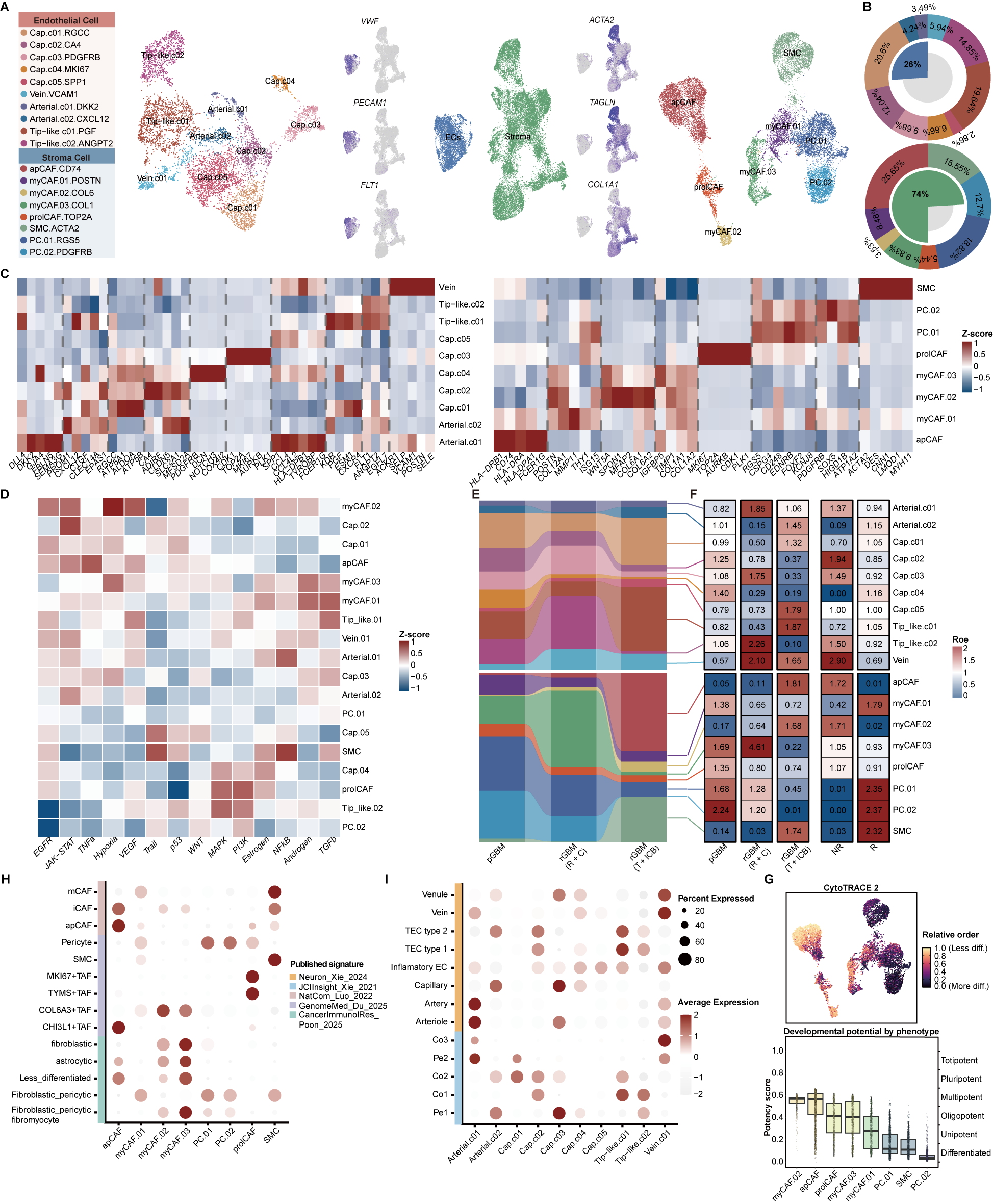

### Figure5_Spatial.tif

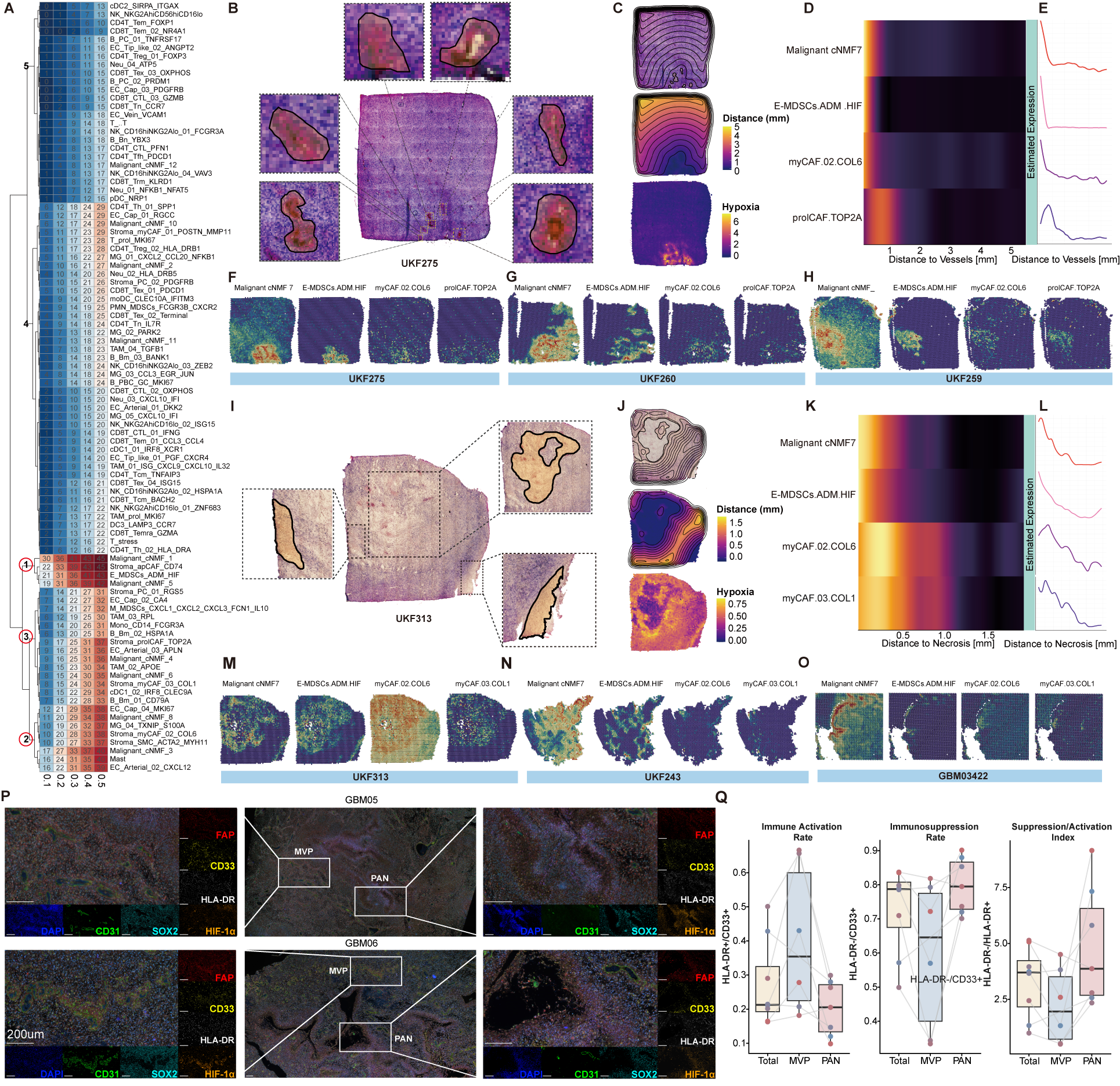

### Figure6_SMI.tif

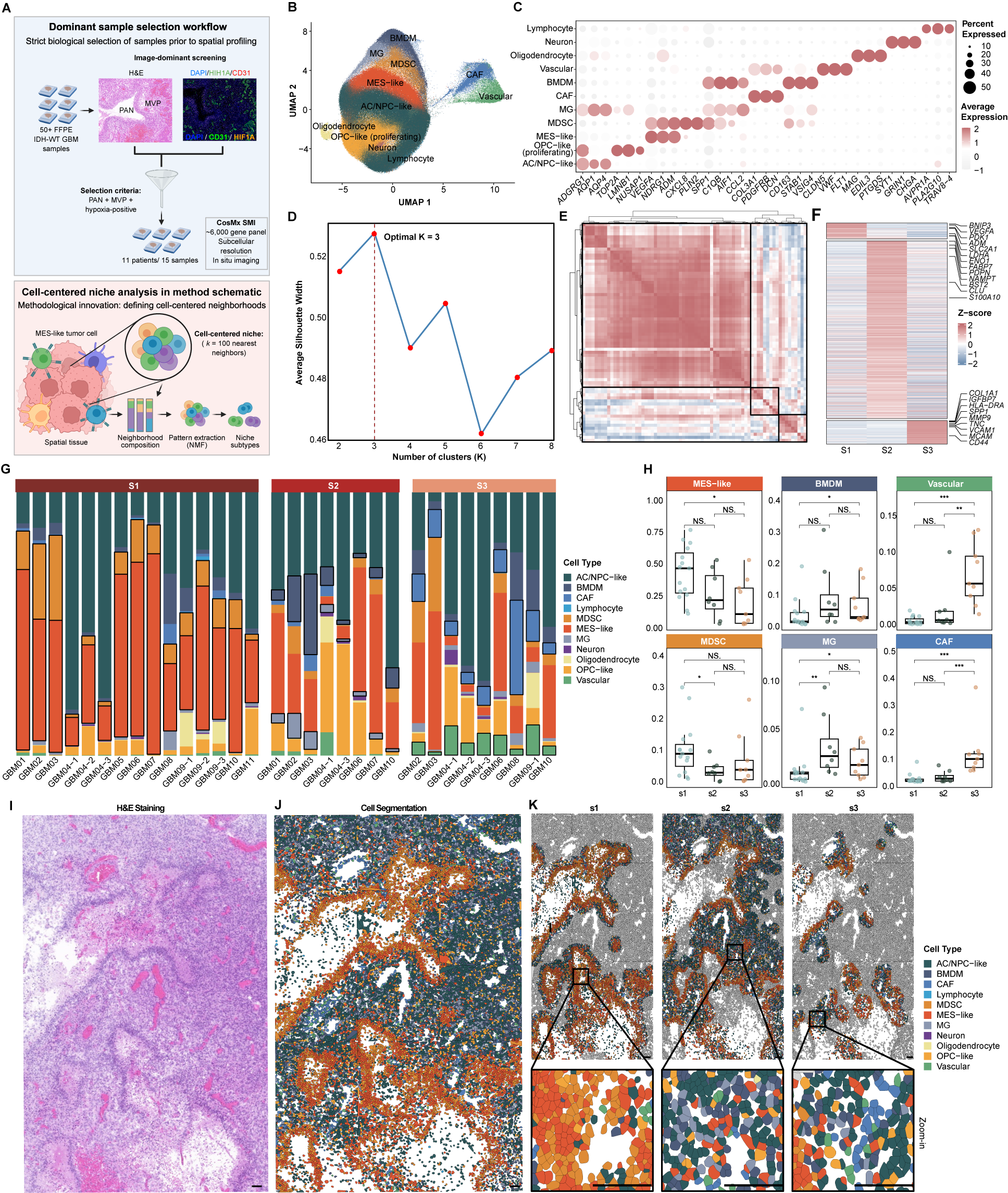

### Figure7_LAB.tif

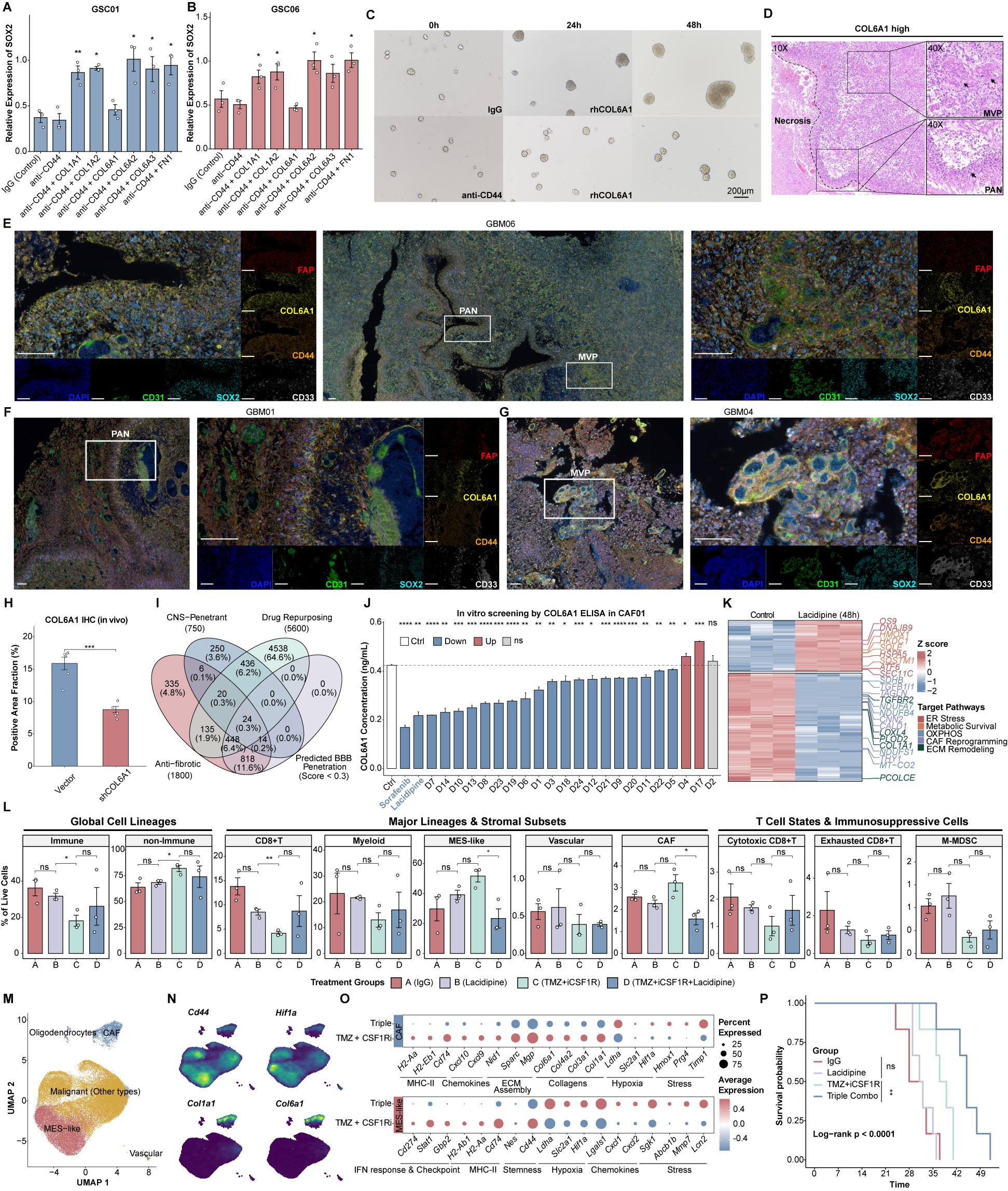

### FigureS1.tif

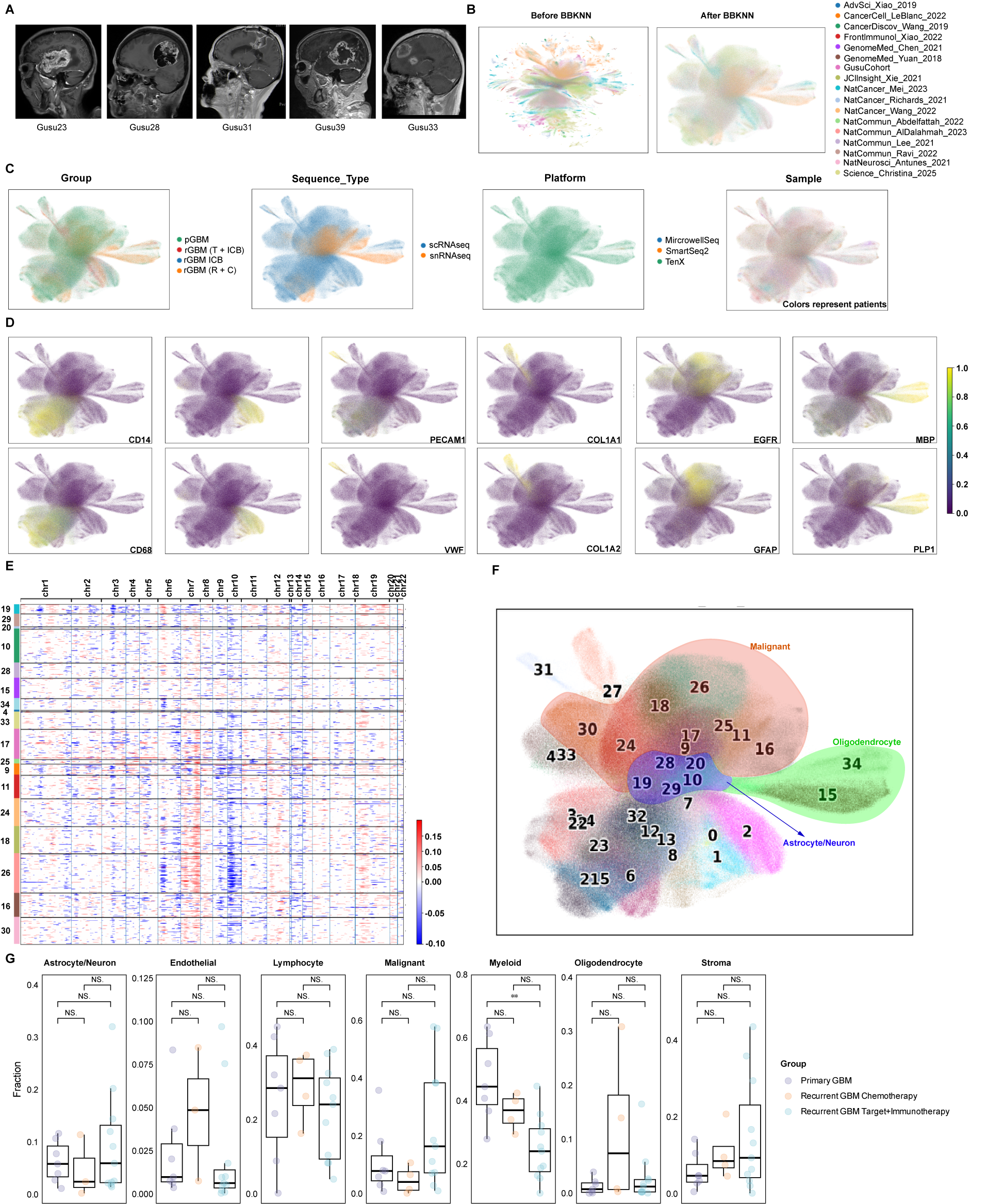

### FigureS2_Malignant.tif

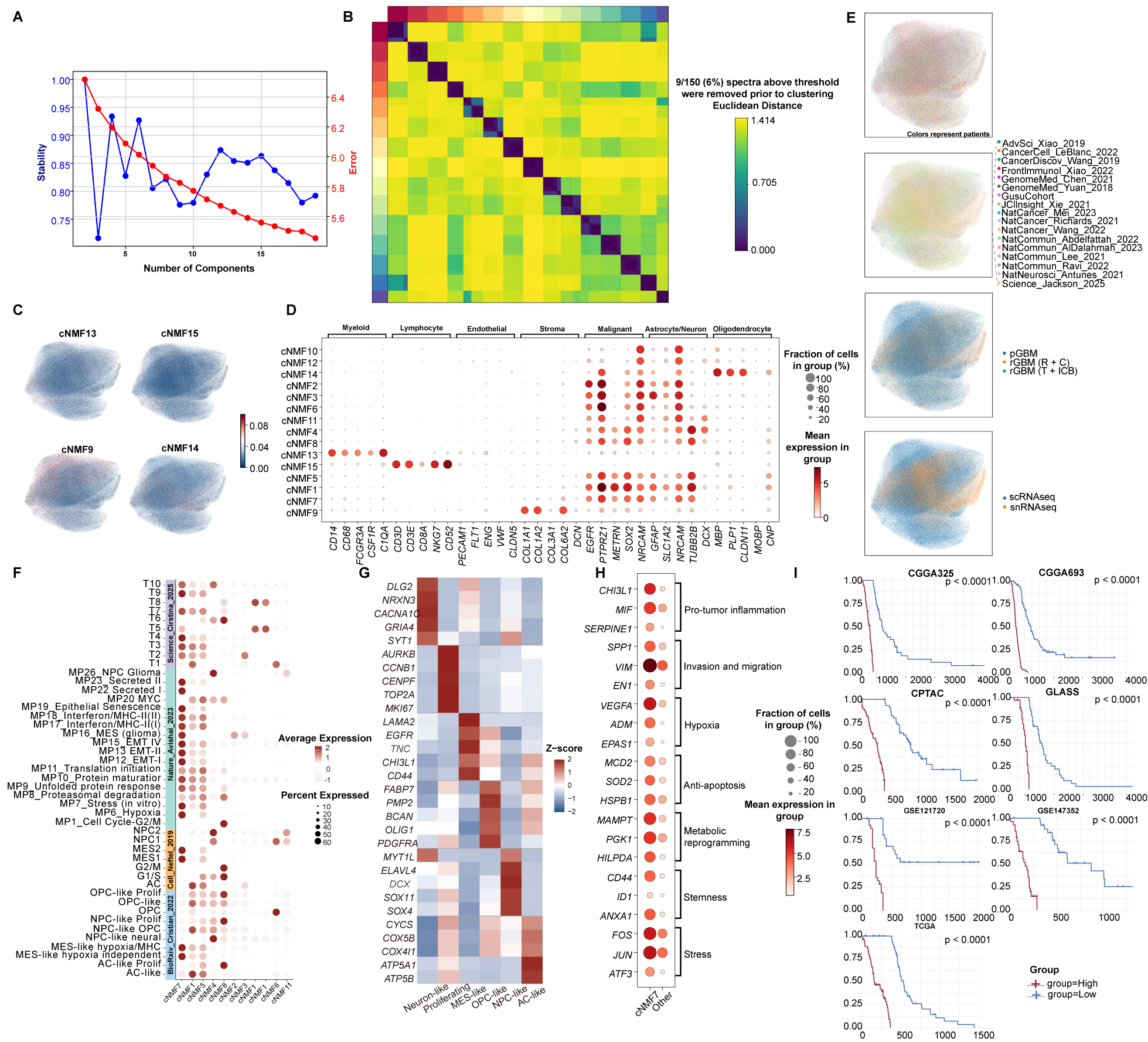

### FigureS3_Myeloid.tif

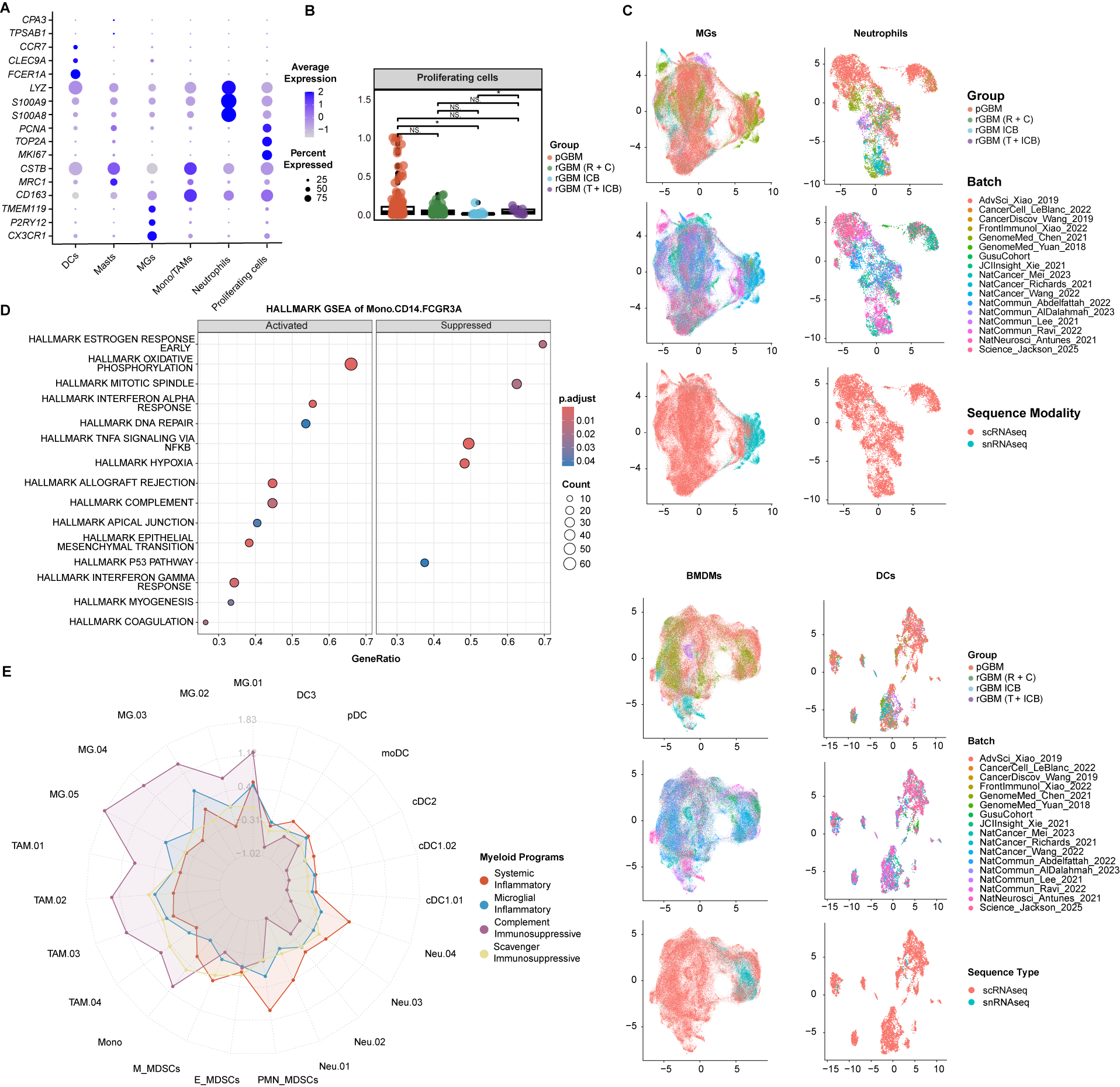

### FigureS4_Lymphocyte1.tif

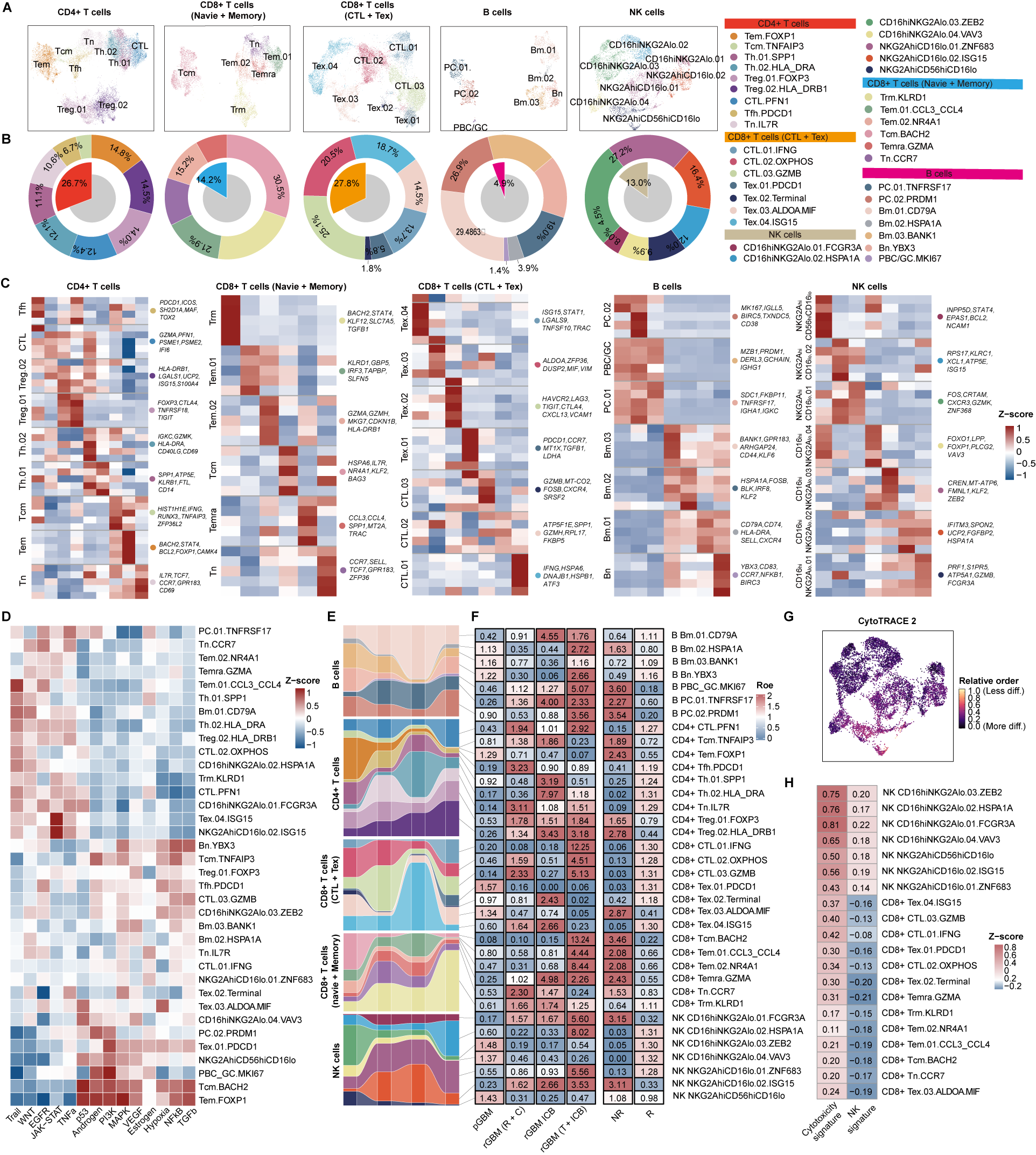

### FigureS5_Lymphocyte2.tif

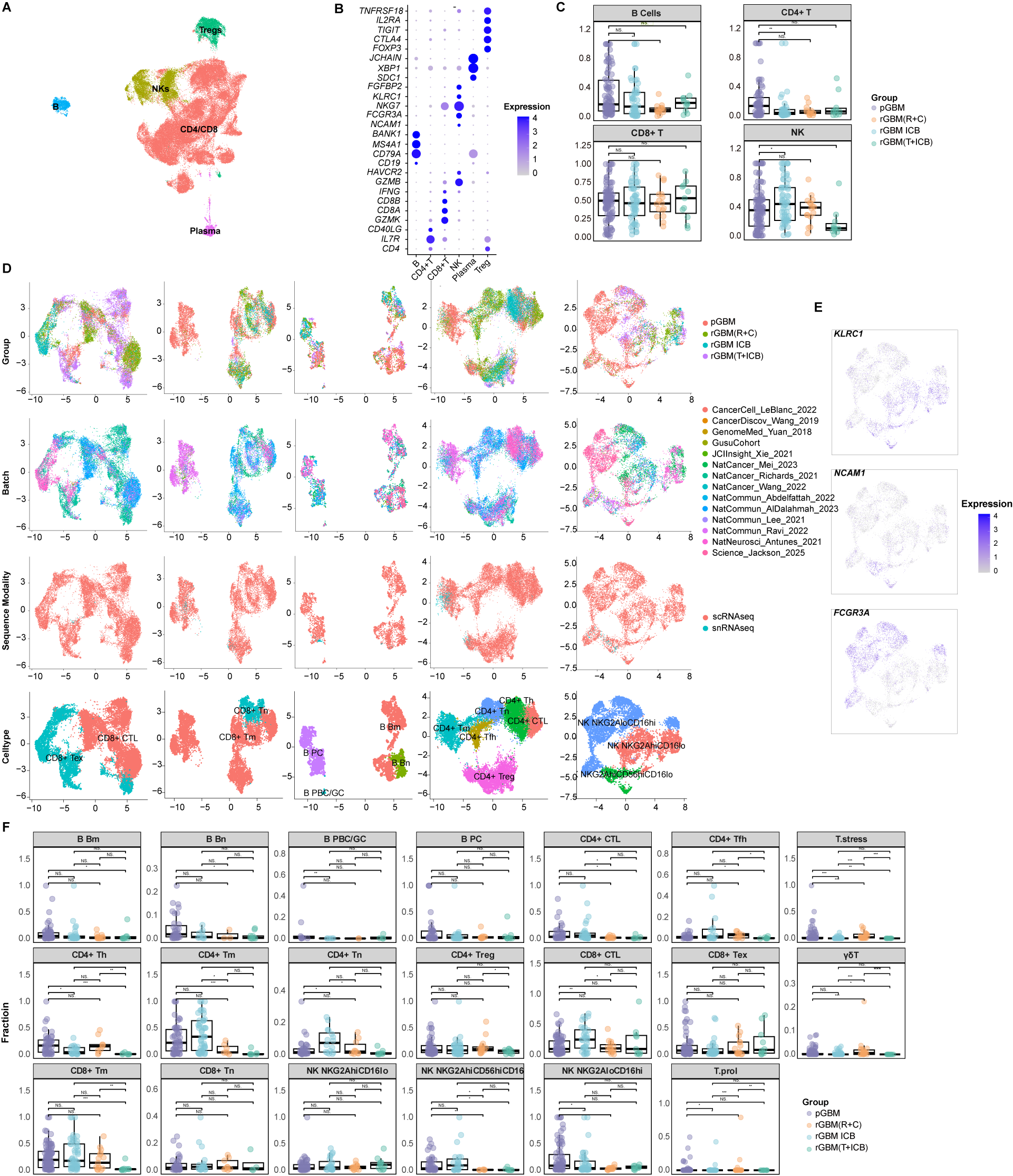

### FigureS7_Spatial.tif

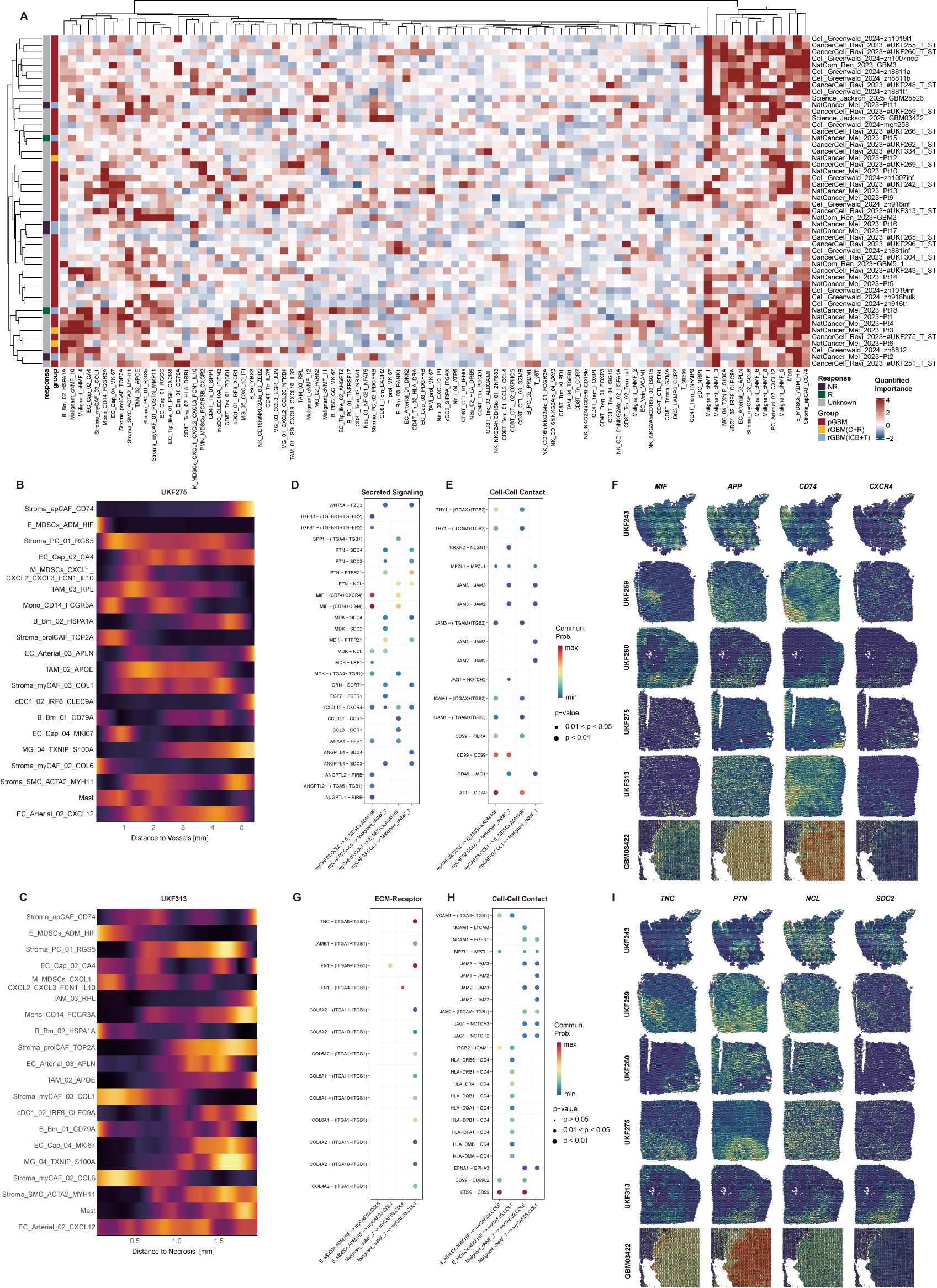

### FigureS8_Cellchat.tif

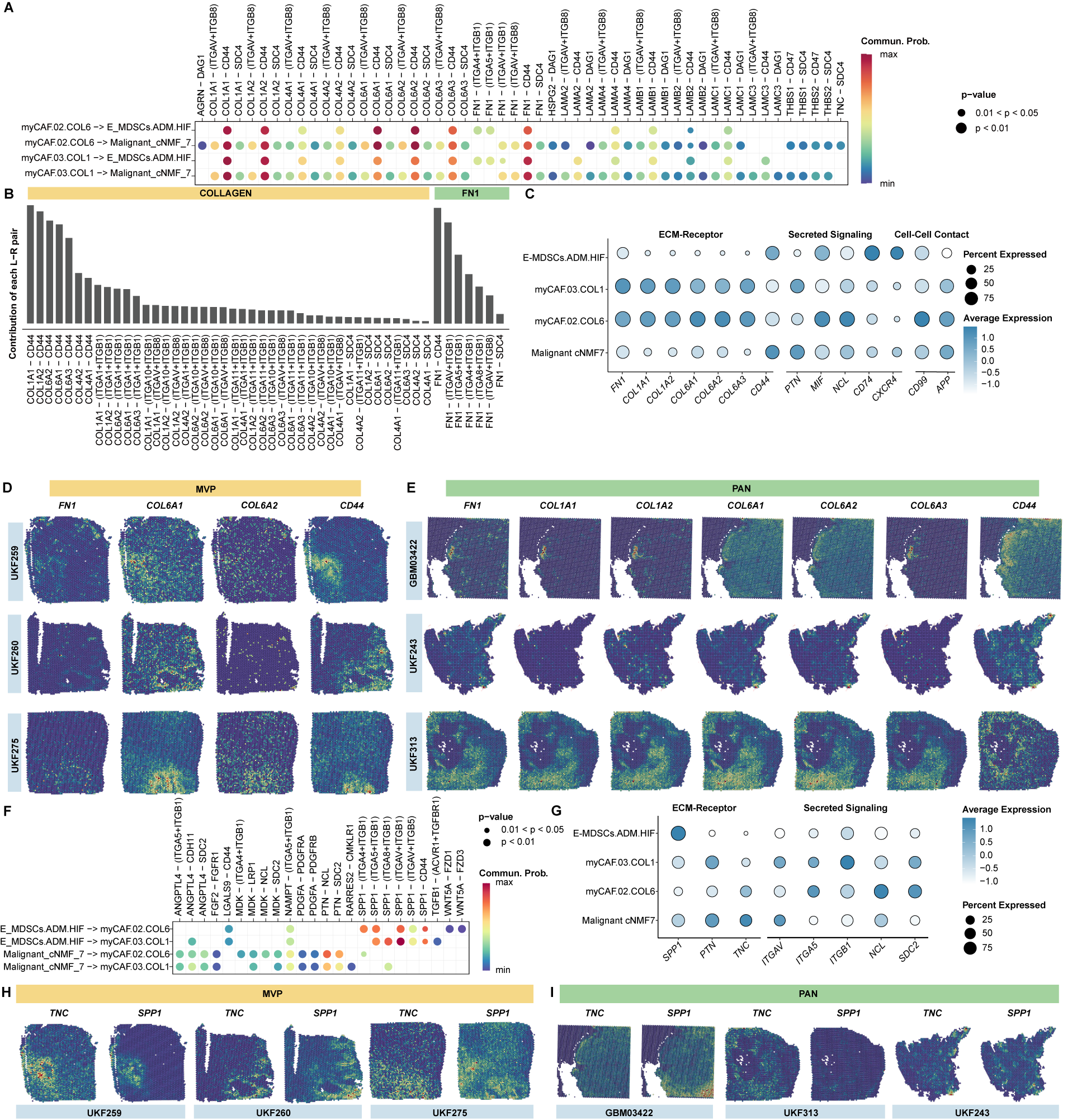

### FigureS9_Bulk.tif

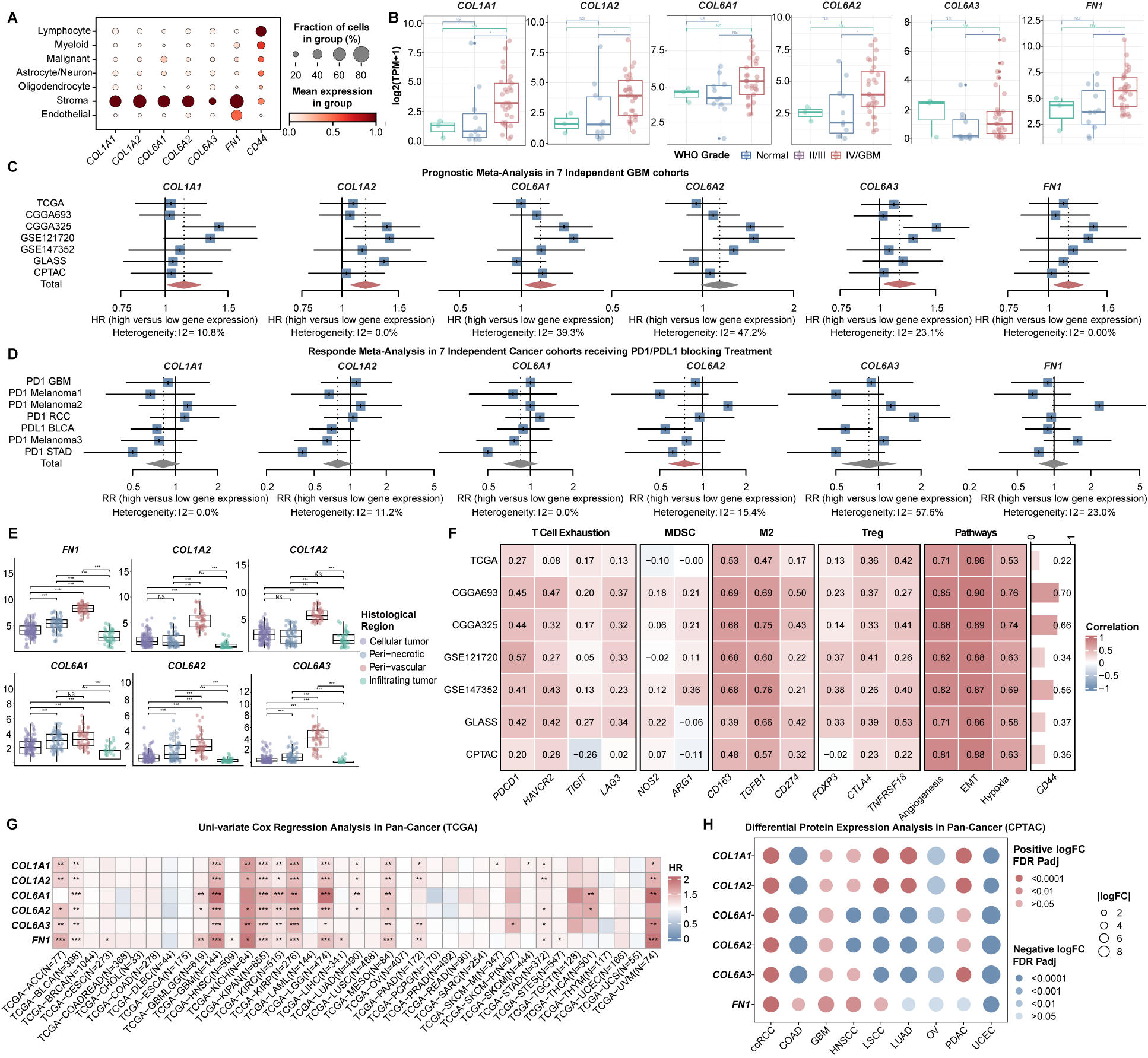

### FigureS10_SMI.tif

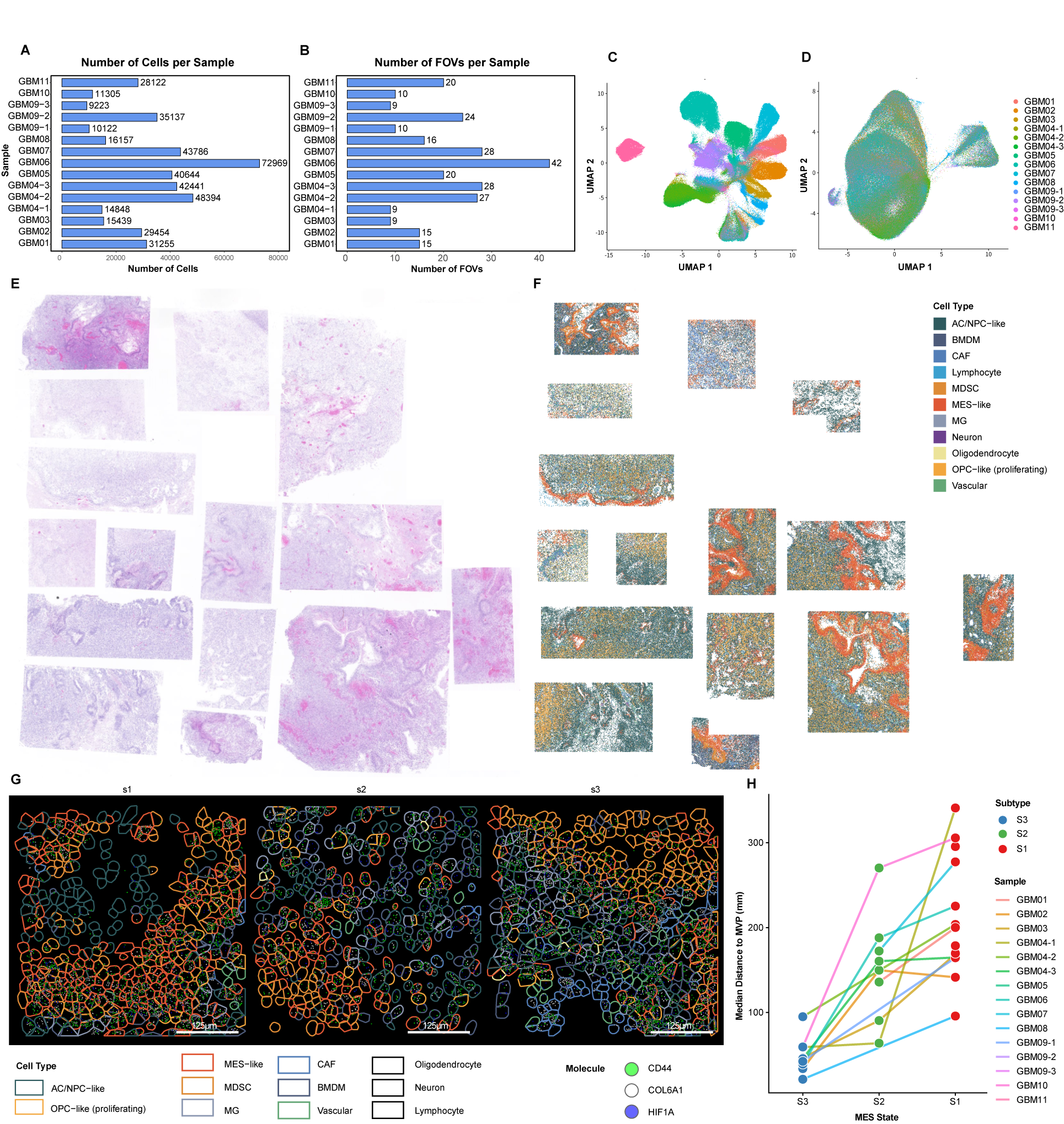

### FigureS11_LAB1.tif

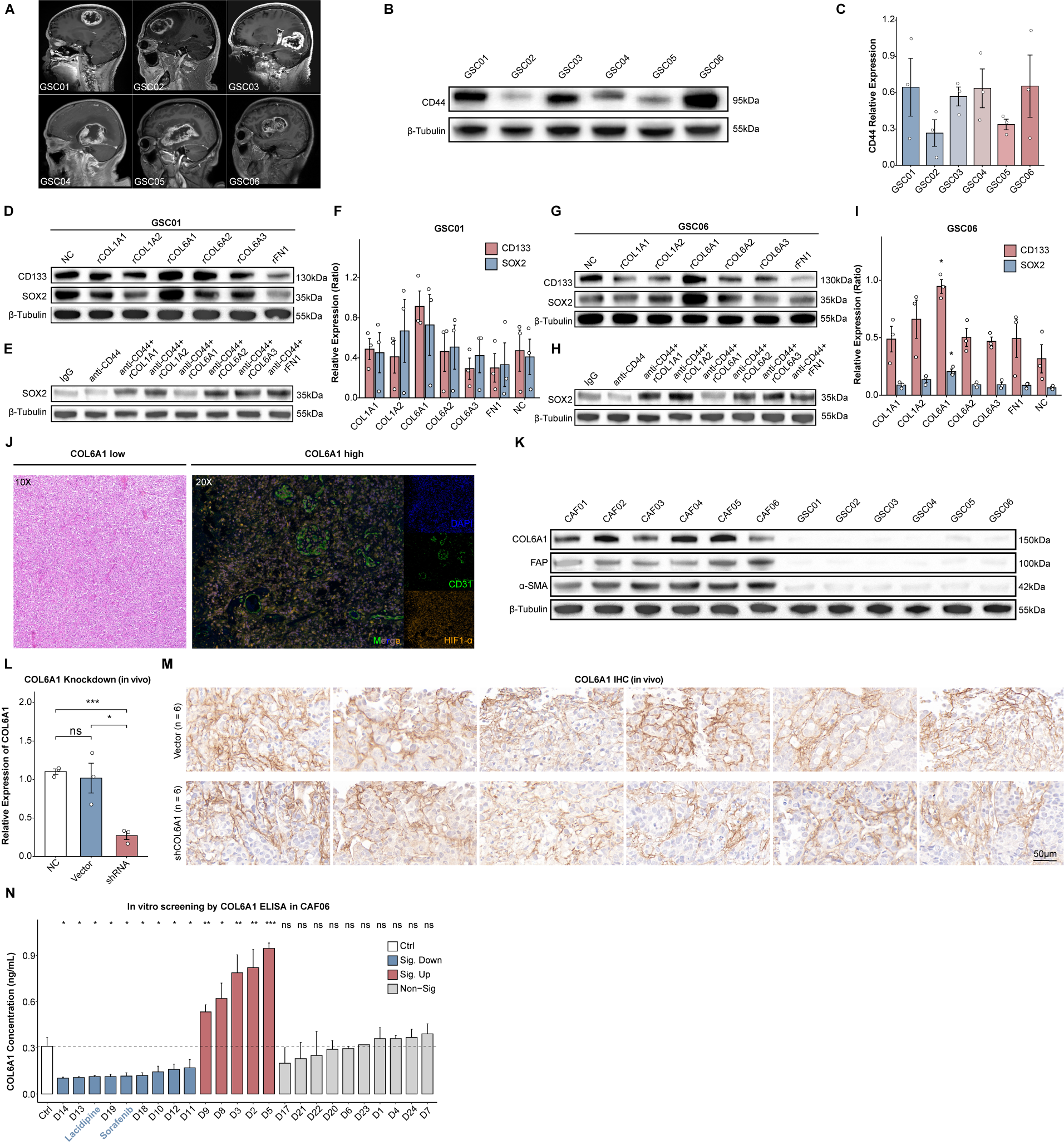

### FigureS12_LAB2.tif

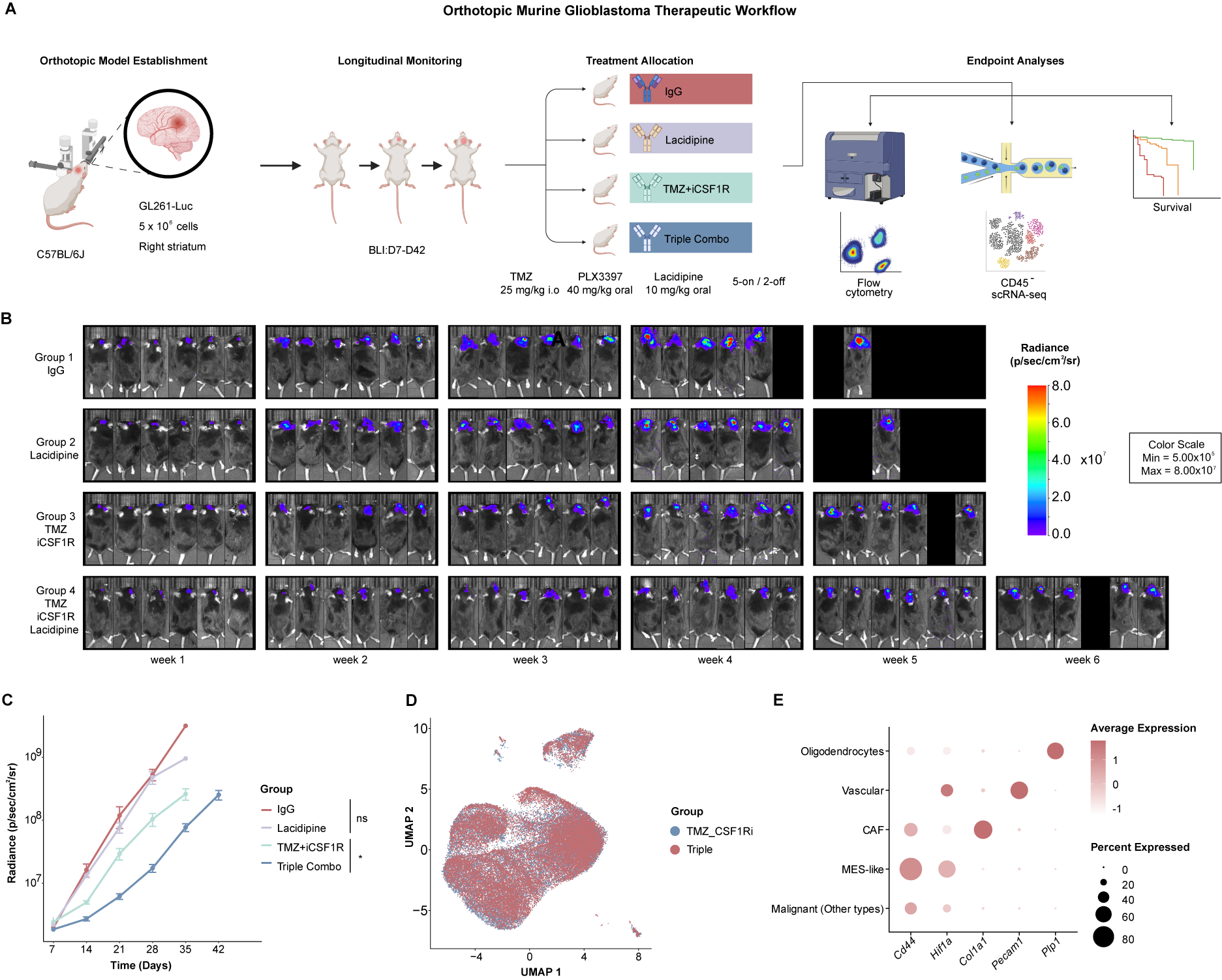
